# A Cheminformatics Workflow for Higher-throughput Modeling of Chemical Exposures from Biosolids

**DOI:** 10.1101/2025.04.03.647109

**Authors:** Paul M. Kruse, Caroline L. Ring

**Affiliations:** Oak Ridge Institute for Science and Education, Oak Ridge, TN; Center for Computational Toxicology and Exposure, Office of Research and Development, United States Environmental Protection Agency, Research Triangle Park. NC

**Keywords:** exposure science, risk assessment, exposure modeling, computational exposure, cheminformatics, biosolids

## Abstract

The U.S. Environmental Protection Agency’s Biosolids Screening Tool can predict potential human and ecological exposures to chemical contaminants in treated sewage sludge biosolids, but large quantities of chemical-specific physico-chemical data are required to parameterize the model. Here, an R workflow is presented that leverages publicly-available databases of chemical information to prepare data for model simulations using the Biosolids Screening Tool. The automated Biosolids Screening Tool workflow (autoBST) reduces the time to gather data necessary to screen hundreds of chemicals from days to just a few minutes and provides transparent and reproducible data retrieval and input into existing models, allowing assessors to defensibly prioritize chemicals in biosolids that may pose a risk to human health or the environment.

## Introduction

The Clean Water Act requires the United States Environmental Protection Agency (US EPA) to establish numerical limits and management practices that protect public health and the environment from the reasonably anticipated adverse effects of toxic pollutants in sewage sludge; and to periodically review existing regulations for the purpose of identifying additional toxic pollutants that may be present in sewage sludge and assesses whether those pollutants may adversely affect public health or the environment based on their toxicity, persistence, concentration, mobility, and potential for exposure.

Based on existing biosolids monitoring data, 726 chemical contaminants have been identified in biosolids (Richman et al. 2022). However, presence in biosolids alone does not necessarily establish a risk to human health or the environment. Potential risk from chemical contaminants in biosolids is estimated by considering both exposure (how much contact a human, animal, or plant has with the chemical resulting from biosolids contamination) and hazard/toxicity (the capability of the chemical to cause adverse effects, and the exposure level at which such adverse effects may occur). Exposures are predicted by modeling the fate and transport of a pollutant through the environment, taking into account different environmental conditions and exposure scenarios. Biosolids-specific exposure models have been implemented in EPA’s publicly-available Biosolids Screening Tool (BST) (US EPA SAB, 2023).

The BST predicts exposures associated with land application of biosolids in agricultural and land reclamation settings or placement in a surface disposal unit, under scenarios of three different meteorologic conditions (wet, average, and dry). It is implemented as a Microsoft Access tool, in which chemical-specific parameters may be entered manually, one chemical at a time, using a pre-programmed graphical interface. Once all necessary parameters for a chemical are entered into the BST, then the BST may be run for that one chemical. Required chemical-specific data are pre-loaded into the BST for 77 chemical substances, but for other chemical substances, they must be added by the user.

The BST can be run only for chemicals with known concentrations in biosolids. Of the 726 chemical contaminants with known presence in biosolids, 484 have measured, quantitative concentrations in biosolids from one of the three National Sewage Sludge Surveys (NSSS) published by EPA, which measured chemical contaminant concentrations in samples of sewage sludge from a representative sample of wastewater treatment plants across the United States in 1988, 2001, and 2006 (Richman et al. 2022). Using biosolids concentrations data along with chemical-specific parameters (including physico-chemical properties and environmental fate and transport properties), the BST can predict chemical fate and transport among various environmental media. Detailed information about the required chemical-specific parameters may be found in the BST User Guide Appendices (USEPA, 2023).

With more than 400 chemicals to consider, gathering BST parameter values one chemical at a time is not feasible, nor is entering them into the BST by hand. Fortunately, chemical-specific data are available for hundreds of thousands of substances from EPA’s CompTox Chemicals Dashboard (CCD), a web application that integrates and makes accessible publicly-available cheminformatics data for computational toxicology and exposure (CTX) (Williams et al. 2017). Recently, CTX data have also been made available through Application Programming Interfaces (APIs), which can be accessed using the open-source R package “ctxR” (Kruse and Ring 2025).

Here, a reproducible workflow that operationalizes CCD CTX data and tools to parameterize and run the BST for hundreds of chemicals with biosolids concentration data is presented. This workflow makes it possible for risk assessors, regulators, and other decision-makers to rapidly predict potential exposures from chemical contaminants in biosolids.

## Materials and Methods

The automated BST workflow (“autoBST”, available at https://github.com/USEPA/CompTox-ExpoCast-autoBST, is a workflow written as an R script. It is designed to take as input a list of chemical identifiers for which biosolids-related exposure calculations are desired; to automatically acquire values for the chemical-specific parameters necessary to run the BST; and to automatically produce as output a properly-formatted Microsoft Excel spreadsheet of BST parameters that can be automatically imported into the BST.

The format of this output Excel spreadsheet is defined by the Bulk Upload Tool (available in Supplemental Material). This tool, from the developers of the BST, consists of an Excel workbook that prescribes a special tabular format for user-input of chemical-specific data so that the Microsoft-Access-based BST can import it automatically. The Bulk Upload Tool contains three “import” sheets on which the user must enter chemical data: one sheet listing the chemical identifiers to be imported; one sheet for the chemical-specific parameters; and one sheet for ecological inputs. The chemical-specific parameter import sheet is organized in a table with columns for the chemical identifier; the name of the BST parameter; the value of the BST parameter; and a reference identifier that identifies the source of the value. This is the required format for the output of autoBST.

The Bulk Upload Tool prescribes not only the format, but also the content, of autoBST output. The Bulk Upload Tool lists the required chemical-specific parameters in tabular form on two “template” sheets, one for organic chemicals and one for inorganic chemicals. There is also a third “template” sheet for ecological parameters, for which the same list of parameters is required for both organic and inorganic chemicals. The “template” sheets also specify the acceptable range of values for each parameter. Each chemical must have a valid numeric value for every parameter expected by the BST. These requirements are strict. Chemicals with incomplete or out-of-range data cannot be bulk-uploaded at all. If, for any chemical, any of the required parameters are not present in the table, or have values outside of the accepted range, then either the bulk-upload process or the BST simulations will fail. In addition, data for chemicals already present in the BST cannot be bulk-uploaded. Supplying new data for these chemicals in the Bulk Upload Tool will not overwrite the existing data, nor will it create a duplicate chemical; instead, the bulk-upload process or the BST simulations will fail.

The autoBST workflow automatically handles the process of locating values for the required parameters, checking their validity, and formatting them for bulk uploading. The appropriate “template” tables (chemical-specific and ecological tables) for each input chemical are filled out by autoBST. The only thing the user needs to provide is a list of chemical identifiers of interest, along with the biosolids concentration for each chemical in units of µg/g dry weight biosolids. Everything else is handled by autoBST.

The steps automatically performed by autoBST are summarized in the following list and described in more detail below.

1. Load user-supplied list of chemical identifiers of interest.
2. Load chemical classifications: “inorganic” or “organic.” Chemicals or substances that cannot be classified as either “inorganic” or “organic” are excluded from further analysis.
3. Identify dioxin-like chemicals (DLCs), including dioxins, furans, and PCBs. These are excluded from BST exposure modeling, as recommended in the BST User Guide.
4. Identify chemicals whose parameters are already loaded into the BST.
5. Load a list of the BST’s required chemical-specific parameters, depending on whether the chemical is organic or inorganic (Table 1 and Table 2).
6. Load user-supplied biosolids concentration data and retrieve chemical name and CASRN.
7. Populate required physico-chemical property data by retrieving EPA data using ctxR.
8. Populate required biotransfer factors from pre-existing tables or predict them from physical-chemical properties using equations as implemented in the BST.
9. Populate required hazard/toxicity values.
10. Populate required ecological data.
11. Check that all necessary data are supplied, rejecting chemicals missing any such data.
12. Check that all supplied values are within acceptable operating ranges for the BST.
  a. If discrepancy is small, adjust to the exceeded limit.
  b. If discrepancy is large, reject chemical.
13. Combine accepted data into tables and output into a correctly-formatted Microsoft Excel file.

**Table 1.** BST parameters for organic chemicals.

| Parameter name | Definition | Units | Accepted range | Default |
| --- | --- | --- | --- | --- |
| CAS | Chemical Abstract Service Registry number | Numerals only – no dashes | — | — |
| Chemical Name | Any text string identifying the chemical | — | — | — |
| CTPWasteDry | Chemical concentration in dry biosolids | µg/g dry weight biosolids | — | — |
| Hazard and toxicity parameters |  |  |  |  |
| CSFOral | Oral cancer slope factor (human toxicity) | per mg/kg-day | — | — |
| IUR | Inhalation unit risk (human toxicity, cancer) | per µg/m³ | — | — |
| Ref_BMD_Bird | Reference benchmark dose (bird) | mg chem/kg BW/day | — | — |
| Ref_BMD_Mammal | Reference benchmark dose (mammal) | mg chem/kg BW/day | — | — |
| Ref_BW_Bird | Reference body weight (bird) | kg | — | — |
| Ref_BW_Mammal | Reference body weight (mammal) | kg | — | — |
| RfC | Reference concentration (human toxicity, noncancer) | mg/m3 | — | — |
| RfD | Reference dose (human toxicity, noncancer) | mg/kg-day | — | — |
| Physical-chemical properties |  |  |  |  |
| MW | Molecular weight | g/mol | 6.0–1000 | — |
| logKow | Log octanol-water partition coefficient | Unitless; log<sub>10</sub>-scale | -4.0–10.0 | — |
| HLC | Henry’s Law constant | atm-m³/mol | 0.0–10.0 | — |
| Sol | Water solubility | mg/L | 1×10<sup>-12</sup>–1×10<sup>6</sup> | — |
| Koc | Organic carbon partition coefficient | mL/g | 0.0–1×10<sup>9</sup> | — |
| Dw | Diffusivity in water | cm²/s | 0.0–0.01 | — |
| Da | Diffusivity in air | cm²/s | 0.01–1.0 | — |
| Bioconcentration factors: Beef and Milk |  |  |  |  |
| BCF_beef | Beef | [mg/kg beef]/[mg/kg DW] | 0.0–8000 | 0 |
| BCF_milk | Milk | [mg/kg milk]/[mg/kg DW] | 0.0–1×10<sup>4</sup> |  |
| Bioconcentration factors: Plants |  |  |  |  |
| BRExFruit | Exposed fruit | [mg/kg DW plant]/[mg/kg soil] | 0.0–10.0 | 0 |
| BrProFruit | Protected fruit |  |  |  |
| BrExVeg | Exposed vegetables |  |  |  |
| BrProVeg | Protected vegetables |  |  |  |
| BrRoot | Root vegetables |  |  |  |
| BrForage | Forage |  |  |  |
| BrGrain | Grain |  |  |  |
| BrSilage | Silage |  |  |  |
| Bv | Air-to-leaf | ([µg contaminant/g DW])/[µg contaminant/g air] |  |  |
| Fish bioaccumulation factors (BAFs) |  |  |  |  |
| Trophic Level 3 fish filet BAF | Human consumption | [mg/kg fish]/[mg/L water] | 0.0–1×10<sup>9</sup> | 0 |
| Trophic Level 3 fish whole BAF | Eco consumption |  |  |  |
| Trophic Level 4 fish filet BAF | Human consumption |  |  |  |
| Trophic Level 4 fish whole BAF | Eco consumption |  |  |  |
| Ecological bioaccumulation factors (BAFs) |  |  |  |  |
| Soil-to-tissue BAFs |  |  |  |  |
| BAF_Worms | Worms | [mg/kg wet weight]/[mg/kg soil] | 0.0–1×10<sup>9</sup> | 1 |
| BAF_Soilinvert | Soil invertebrates |  |  |  |
| BAF_HerbVert | Herbivorous vertebrates |  |  |  |
| BAF_OmnVert | Omnivorous vertebrates |  |  |  |
| BAF_SmBirds | Small birds |  |  |  |
| BAF_SmMammals | Small mammals |  |  |  |
| BAF_SmHerp | Small herpetofauna |  |  |  |
| Aquatic systems bioconcentration factors (BCFs) |  |  |  |  |
| BCF_WaterVeg | Surface water to aquatic plants | [mg/kg wet weight]/[mg/L water] | 0.0–1×10<sup>9</sup> | 1 |
| BCF_Bff | Sediment to biota | [mg/kg wet weight]/[mg/kg sediment] |  |  |
| Degradation rates |  |  |  |  |
| kaer | Aerobic biodegradation rate in the water column | 1/day | 0–50 | 0 |
| kanaer | Anaerobic biodegradation rate in the sediment | 1/day | 0–2 | 0 |
| kh | Hydrolysis rate | 1/day | 0–50 | 0 |
| kpo | Photolysis rate | 1/day | 0–50 | 0 |
| Ksoil | Soil biodegradation rate | 1/day | 0–100 | 0 |

**Table 2.** BST parameters for inorganic chemicals.

| Parameter name | Definition | Units | Accepted range | Default |
| --- | --- | --- | --- | --- |
| CAS | Chemical Abstract Service Registry number | Numerals only – no dashes | — | — |
| Chemical Name | Any text string identifying the chemical | — | — | — |
| CTPWasteDry | Chemical concentration in dry biosolids | µg/g dry weight biosolids | — | — |
| Hazard and toxicity parameters |  |  |  |  |
| CSFOral | Oral cancer slope factor (human toxicity) | per mg/kg-day | — | — |
| IUR | Inhalation unit risk (human toxicity, cancer) | per µg/m³ | — | — |
| Ref_BMD_Bird | Reference benchmark dose (bird) | mg chem/kg BW/day | — | — |
| Ref_BMD_Mammal | Reference benchmark dose (mammal) | mg chem/kg BW/day | — | — |
| Ref_BW_Bird | Reference body weight (bird) | kg | — | — |
| Ref_BW_Mammal | Reference body weight (mammal) | kg | — | — |
| RfC | Reference concentration (human toxicity, noncancer) | mg/m3 | — | — |
| RfD | Reference dose (human toxicity, noncancer) | mg/kg-day | — | — |
| Physical-chemical properties |  |  |  |  |
| MW | Molecular weight | g/mol | 6.0–1000 | — |
| Density | Density | g/cm³ | 0.0–23 | — |
| Dw | Diffusivity in water | cm²/s | 0.01–0.01 | — |
| Kd | Soil partition coefficient | L/kg | 0.0–1×10⁶ | — |
| Bioconcentration factors: Beef and Milk |  |  |  |  |
| BCF_beef | Beef | [mg/kg beef]/[mg/kg DW] | 0.0–8000 | 0 |
| BCF_milk | Milk | [mg/kg milk]/[mg/kg DW] | 0.0–1×10⁴ |  |
| Bioconcentration factors: Plants |  |  |  |  |
| BRExFruit | Exposed fruit | [mg/kg DW plant]/[mg/kg soil] | 0.0–10.0 | 0 |
| BrProFruit | Protected fruit |  |  |  |
| BrExVeg | Exposed vegetables |  |  |  |
| BrProVeg | Protected vegetables |  |  |  |
| BrRoot | Root vegetables |  |  |  |
| BrForage | Forage |  |  |  |
| BrGrain | Grain |  |  |  |
| BrSilage | Silage |  |  |  |
| Bv | Air-to-leaf | ([µg contaminant/g DW]/[µg contaminant/g air]) | Assumed 0 for inorganic chemicals because these are generally non-volatile |  |
| Fish bioaccumulation factors (BAFs) |  |  |  |  |
| Trophic Level 3 fish filet BAF | Human consumption | [mg/kg fish]/[mg/L water] | 0.0–1×10⁹ | 0 |
| Trophic Level 3 fish whole BAF | Eco consumption |  |  |  |
| Trophic Level 4 fish filet BAF | Human consumption |  |  |  |
| Trophic Level 4 fish whole BAF | Eco consumption |  |  |  |
| Ecological bioaccumulation factors (BAFs) |  |  |  |  |
| Soil to tissue BAFs |  |  |  |  |
| BAF_Worms | Worms | [mg/kg wet weight]/[mg/kg soil] | 0.0–1×10⁹ | 1 |
| BAF_Soilinvert | Soil invertebrates |  |  |  |
| BAF_HerbVert | Herbivorous vertebrates |  |  |  |
| BAF_OmnVert | Omnivorous vertebrates |  |  |  |
| BAF_SmBirds | Small birds |  |  |  |
| BAF_SmMammals | Small mammals |  |  |  |
| BAF_SmHerp | Small herpetofauna |  |  |  |
| **Aquatic systems bioconcentration factors (BCFs)** |  |  |  |  |
| BCF_WaterVeg | Surface water to aquatic plants | [mg/kg wet weight]/[mg/kg sediment] | 0.0–1×10<sup>9</sup> | 1 |
| BCF_Bff | Sediment to biota |  |  |  |

### Step 1: Chemical identifiers

In step 1, the chemical identifiers of interest provided by the user should be DSSTox Substance IDs (DTXSIDs). These are unique identifiers for substances in EPA’s Distributed Searchable Substance-Toxicity database (Grulke et al. 2019; Richard and Williams 2002). Chemical identifiers of other types, such as Chemical Abstract Service Registry Numbers (CASRNs), chemical names, or InChIKeys, should be mapped to DTXSIDs before beginning the workflow; this can be done easily by using either the CCD’s web-based batch search, or using functions from the R package ctxR that retrieve DTXSIDs matching user-input chemical identifiers including full or partial name, CASRN, or InChIKey.

### Step 2: Chemical classifications

In step 2, autoBST automatically assigns the chemical classifications “inorganic” and “organic” using the ClassyFire chemical classification tool, which classifies chemicals based on their structure using the ChemOnt ontology (Djoumbou Feunang et al. 2016). ClassyFire provides a hierarchical chemical classification with levels kingdom, superclass, class, subclass, and so on, up to seven additional levels (levels 5 through 11). At present, the ClassyFire tool available on the web can be queried by supplying the chemical’s InChIKey, which identifies its structure. Chemical identifiers such as DTXSIDs can be mapped to InChIKeys using the CTX APIs described above. If the chemical does not have an InChIKey, or if ClassyFire does not return a classification of “inorganic” or “organic,” then the chemical is excluded from further analysis.

### Step 3: Dioxin-like compounds

In step 3, autoBST identifies dioxin-like compounds (DLCs) using their ClassyFire classifications: if their class is either “Benzodioxins” or “Benzofurans,” or their level-6 classification is “Polychlorinated biphenyls,” then they are designated DLCs and excluded from further analysis. DLCs are excluded from BST exposure modeling, as the BST fate and transport algorithms are not appropriate for this class of chemicals, according to the BST User Guide (USEPA, 2023).

### Step 4: Pre-loaded chemicals

In step 4, autoBST compares the user-input list of chemicals to the list of 77 chemicals pre-loaded into the BST, and removes any user-input chemicals that are already pre-loaded.

### Step 5: Determine required chemical-specific parameters

In step 5, autoBST determines the chemical-specific parameters required to run the BST, depending on whether the chemical is organic (Table 1) or inorganic (Table 2).

### Step 6: Biosolids concentration and chemical identifiers

CTPWasteDry is the chemical concentration in biosolids in units of µg/g dry weight biosolids. It must be supplied to autoBST as a table of DTXSIDs and concentrations.

CAS and ChemicalName are the chemical identifiers used within the BST. autoBST acquires them automatically from the user-supplied DTXSID by querying CTX databases using the R package ctxR (described above) with the function ctxR::get_chemical_details().

### Step 7: Physical-chemical properties

In this step, autoBST acquires values for physico-chemical properties using the CTX APIs.

Molecular weight (MW), octanol-water partition coefficient (log K_ow_), Henry’s Law Constant (HLC), water solubility (Sol), and organic-carbon partition coefficient (K_oc_) are all acquired from CTX databases, based on the DTXSID, by using ctxR.

If experimental values are available for a property, then autoBST uses them. (If more than one experimental value is available for a property, then the average is taken.) If no experimental values are available for one of these properties, then QSAR-predicted values are used. (If more than one QSAR-predicted value is available for a property, then the average is taken.)

For MW, log K_ow_, HLC, and Sol, if neither experimental nor QSAR-predicted values are available, then autoBST assigns the value as NA (Not Available). (Any chemicals with missing data at this stage, or any stage in steps 7 and 8, could simply skip the rest of steps 7 and 8 and have NA assigned to all of their chemical-specific properties, since they will be excluded from analysis anyhow in step 9. However, the autoBST continues to attempt to find values for other chemical-specific parameters, so that it can identify the full set of parameters that are missing values for each chemical.)

For K_oc_, if neither experimental nor QSAR-predicted values are available, then autoBST predicts the value using an equation that estimates K_oc_ based on chemical ionization at pH 7.0, pKa, pKb, and log K_ow_. This equation is the same one implemented in the well-established wastewater treatment fugacity model SimpleTreat 4.0 (Lautz et al. 2017). The chemical ionization at pH 7.0 is determined using the method and physicochemical data implemented in the R package httk (function httk::calc_ionization()) (Pearce et al. 2017). The pKa and pKb values were those predicted by the OPEn structure–activity/property Relationship App (OPERA) QSAR, version 9.2 (Mansouri et al. 2018); they were predicted by running the desktop OPERA application, rather than being queried from CTX databases, because CTX APIs did not yet include pKa and pKb at the time this analysis was performed.

For Kd (the soil partition coefficient), used only for inorganic chemicals, chemical-specific data are extremely limited. In the absence of chemical-specific data, autoBST uses a constant value of 1×105, reflecting the conservative assumption that all of the chemical binds to the solid component of sewage sludge and will not wash out.

Air and water diffusivity constants are estimated by autoBST using methods developed by Tucker and Nelken (1982). Air diffusivity constant Da is predicted based on temperature (assumed to be 25 degrees Celsius), pressure (assumed to be 1 atm), the molecular weights of air and the chemical of interest, and the diffusion volume (molar volume) of air and the chemical of interest. The molecular weight of air is assumed to be approximately 28.97 g/mol, and the diffusion volume of air is assumed to be approximately 20.1 cm3/mol. Water diffusivity constant Dw is predicted based on the viscosity of water (assumed to be 0.8904 mPa-s) and the LaBas estimate of diffusion volume of the chemical of interest. The molar volume is used in place of the diffusion volume (or its LaBas estimate). The molar volume of the chemical of interest is determined by querying CTX databases: if experimental values are available, they are used; otherwise, predicted values are used if available; if neither are available, then the molar volume, and therefore Da and Dw, are assigned as NA.

For this analysis, all degradation rates (used for organic chemicals only, Table 1) were set to their default value of zero, reflecting a conservative assumption that no degradation occurs.

### Step 8: Biotransfer factors

autoBST predicts bioconcentration factors (BCFs) for beef, milk, and plants from log K_ow_ for each chemical, using equations originally implemented in the BST (see BST User’s Guide, Appendix C) (USEPA, 2023). A single soil-to-plant BCF is used for all aboveground vegetation (all plant BCFs except for root vegetables), and a root concentration factor is calculated for root vegetables. The air-to-leaf biotransfer factor is calculated from log K_ow_ and Henry’s Law constant for organic chemicals, but is set to 0 for inorganic chemicals, following the assumptions of the BST. If log K_ow_ is NA, then BCFs are all set to NA as well. If the Henry’s Law constant is NA, then the air-to-leaf biotransfer factor is set to NA as well.

autoBST draws fish bioaccumulation factors (BAFs) from the National Bioaccumulation Factor (NBAF) tables when available (USEPA, 2016). If no values are available from the NBAF tables for Tier 3, Tier 4, or both, then autoBST gap-fills the missing value(s) by using CTX APIs to query the CCD for bioconcentration factor values available under the “Fate” endpoint, which may include measured values from the ECOTOX database (Olker et al. 2022) and/or QSAR-predicted values from OPERA (Mansouri et al. 2018). In this case, if one or more measured aquatic BCF values are available from the ECOTOX database, then autoBST uses the geometric mean of available values; if no measured values are available, but one or more OPERA-predicted values are available, then autoBST uses the geometric mean of those values. Any fish BAFs with no values available from any of these sources are assigned as NA.

### Step 9: Hazard/toxicity related values

For the purposes of this analysis, which were to predict human average daily dose (ADD) from all ingestion exposures, all hazard/toxicity related values (reference doses, reference concentrations, cancer slope factors, and inhalation unit risks) were assigned a placeholder value of 1. Placeholder values were used for two reasons. First, the process for selecting the most appropriate values for these hazard/toxicity parameters is complicated enough to be outside the scope of autoBST, requiring expert input particularly for the large number of chemicals that do not have a reference dose (RfD) established through EPA’s Integrated Risk Information System (IRIS). (See the BST User Guide Appendix D for more details.) Second, and more practically, a placeholder value of 1 enables the BST to output the human ADD directly. The BST output does not include ADD in mg/kg/day; it includes only the hazard quotient (HQ) calculated from that ADD (HQ = ADD/RfD). When RfD is given a placeholder value of 1, then the BST-output “HQ” corresponds to the ADD itself.

### Step 10: Ecological data

At present, all ecological BAFs/BCFs and ecological toxicity values are assigned placeholder values of 1. These placeholder values allow all BST simulations to run, but the resulting estimates of ecological exposure and risk are not meaningful and are not used in this case study. Placeholder ecological values are used because the goal of the present BST simulations was to predict biosolids-specific human exposures. However, functionality to format the required table of ecological data was built into autoBST to facilitate future modeling efforts that may include simulation of ecological exposures.

### Step 11: Check for missing data

The BST will not run if any chemicals are missing one or more required parameters. autoBST therefore determines which, if any, information is missing for each chemical on each type of BST input sheet: organic, inorganic, and ecological. If there are missing data, then autoBST outputs the names of the pieces of data that are missing.

### Step 12: Check for data within BST operating range

In this step, autoBST checks whether chemical data are within the BST’s pre-specified operating ranges (see Table 1 and Table 2 for ranges). Similarly, there are helper functions for organic and inorganic chemicals that check to see whether input data fall within the operating range for the BST. These take in a data frame of physico-chemical property data and check that each input value is within the range of allowable values. If the values exceed the maximum value or are below the minimum value (for non-zero lower bounds), the function tests whether the difference is greater than 1% of the violated bound. If so, the property name is recorded, and if not, the property value is set to the value of the boundary it has exceeded. For lower bounds of zero, the values are not set to zero and are instead recorded as out of bounds.

### Step 12: Export correctly-formatted spreadsheets

In step 12, autoBST generates a list of data frames, where each list item corresponds to one of the required sheets in the Bulk Upload Tool. This list can be written to an Excel workbook using the `writexl` package, function `writexl::write_xlsx()`.

### Perform Bulk Upload and run the BST

After running autoBST, the following steps must be performed by the user to bulk-upload the chemicals into the BST and run the BST. To use the Bulk Upload Tool, the workbook that was output by autoBST must be saved with the filename “ChemicalImport.xlsx” in the same directory as the BST .mdb file. Then, the user can open the BST, click the “Configure” button, go to the “Chemicals” tab, and click the “Bulk Import Chemicals” button. Without any further confirmation or interaction required from the user, all the chemicals in the prepared Excel workbook will be imported to the BST, and will appear as new items on the Chemicals tab.

The bulk-upload process has limitations that are inherent to the BST’s Microsoft Access implementation. Once uploaded, chemicals cannot be bulk-deleted from the BST. To remove user-added chemicals, they must be deleted manually, one at a time. Also, once uploaded, chemicals cannot be bulk-overwritten.

For example, if the user makes a change to autoBST that affects a parameter value for all chemicals, they cannot bulk-re-upload the chemicals and overwrite their data to reflect the changes. For this reason, it is recommended to keep a “clean” version of the BST .mdb file, containing only the default preloaded chemicals. To bulk-upload chemicals, first create a copy of the .mdb file, and save it with a different name. Then open the new copy of the BST and bulk-upload chemicals to that copy. To make a change to autoBST and re-upload chemicals with new parameter values, go back to the “clean” version, make a new copy, and start again.

Once the bulk-upload process is complete, the user can run the BST as usual. See the BST User Guide for instructions (USEPA, 2023); see also further instructions and tips in the R Markdown file containing the autoBST code (Supplementary Material).

### Case study

To demonstrate the use of autoBST, it was run for the curated list of 726 chemicals plus 6 nutrients that have been detected in biosolids (Richman et al. 2022). For biosolids concentrations of these chemicals (parameter CTPWasteDry), two main scenarios were considered: the mean and 95th percentile concentrations reported from the most recent available NSSS (calculated with non-detects set equal to half of the Minimum Reporting Limit) (Richman et al. 2022). All NSSS concentrations were converted from their original reported units into units of µg/g dry weight biosolids to match the input units required by the BST. If a chemical on the curated biosolids list was not monitored in any NSSS, or if it was monitored but was not detected in a sufficient number of samples to calculate a mean or 95th percentile concentration, then it was excluded from further analysis. The following chemicals were excluded: chromium (III), chromium (VI), mercury (divalent), selenium (IV), selenium (VI), albuterol, chlortetracycline, oxolinic acid, PFOA, PFOS, and sulfathiazole. (In the NSSS, chromium, mercury, and selenium were measured but not speciated; PFOA and PFOS were not measured; and the other chemicals were measured, but were not detected in enough samples to compute a mean or 95th percentile concentration.)

Additionally, a third scenario was considered where biosolids concentrations were set equal to 1 ppb (0.001 µg/g dry weight biosolids) for all chemicals, which provides a relative ranking of potential for exposure. In this scenario, all chemicals on the curated list were included.

Some chemicals were already pre-loaded into the BST. Pre-loaded chemicals cannot be edited or overwritten using autoBST, so they were edited manually. All pre-loaded chemicals were manually edited within the BST to set their reference dose (RfD) values equal to 1, the same as for autoBST. Pre-loaded chemicals were also manually edited to set their biosolids concentration (CTPWasteDry) equal to the appropriate concentration: the most recent NSSS mean concentration, the most recent NSSS 95th percentile concentration, or 1 ppb (1 × 10-3 mg/kg dry weight).

The BST was run under three land application use (LAU) scenarios (Crop, Pasture, and Reclamation), assuming average rainfall (as opposed to dry or wet conditions). Other than the chemical-specific parameters, all other parameters of the BST were left at their default values; see the BST User Guide for details (USEPA, 2023). All three LAU scenarios simulate exposures to a family living and farming on the land, including ingestion of drinking water, soil, and fish caught on the farm. For the crop scenario, the family eats contaminated produce grown on the farm; for the pasture and reclamation scenarios, the family eats beef and milk grown on the farm, where the cattle eat contaminated forage and soil. (The BST simulates inhalation exposures as well, but ingestion exposures were of primary interest in this analysis.) The BST also predicts exposures for surface-disposal scenarios (in which biosolids are placed in landfills); however, for the sake of tractability of this case study, only LAU scenario results are presented here.

The BST simulation results were saved as an Microsoft Excel file and then imported into R. Then, for each LAU scenario, and each biosolids concentration scenario, the chemicals were ranked according to the BST-predicted average daily dose through the oral (ingestion) pathway for an adult farmer. Linear regression analyses were performed to assess the relation between log_10_ ADD and log_10_ biosolids concentration. Additional linear regression analyses added log_10_ K_ow_ as a second regression term.

## Results

From inputting the list of chemicals at the start to generating the bulk upload file at the end, the entire BST Workflow process took approximately 15 minutes on a Dell laptop with 13th Gen Intel(R) Core(TM) i7-1365U 1.80 GHz processor and 16 gigabytes of RAM. BST simulations took roughly 30 hours to complete.

When biosolids concentrations were assumed to be 1 ppb for all 726 chemicals plus 6 nutrients on the curated biosolids list, the BST was able to make exposure predictions for 434 of those chemicals. The data filtering steps are illustrated in Figure 1. 71 of the curated biosolids chemicals were pre-loaded into the BST, and therefore were excluded from further BST Workflow filtering steps. 41 chemicals were excluded because they could not be classified as either organic or inorganic by ClassyFire. 30 of these could not be classified because they did not have InChIKeys; mainly, these were mixtures without a single defined structure (e.g., aroclors, polycarbonates, butylated hydroxyanisole, branched 4-nonylphenol, tert-butylphenyl diphenyl phosphates, tricresyl phosphate). Eleven other chemicals had InChIKeys and defined structures, but ClassyFire did not have a classification available when queried by InChIKey: these included aldrin, dieldrin, endrin, lindane, endosulfan I, endosulfan II, heptachlor epoxide B, dl-norgestrel, alpha-1,2,3,4,5,6-Hexachlorocyclohexane, beta-Hexachlorocyclohexane, and delta-Hexachlorocyclohexane.

**Figure 1:**
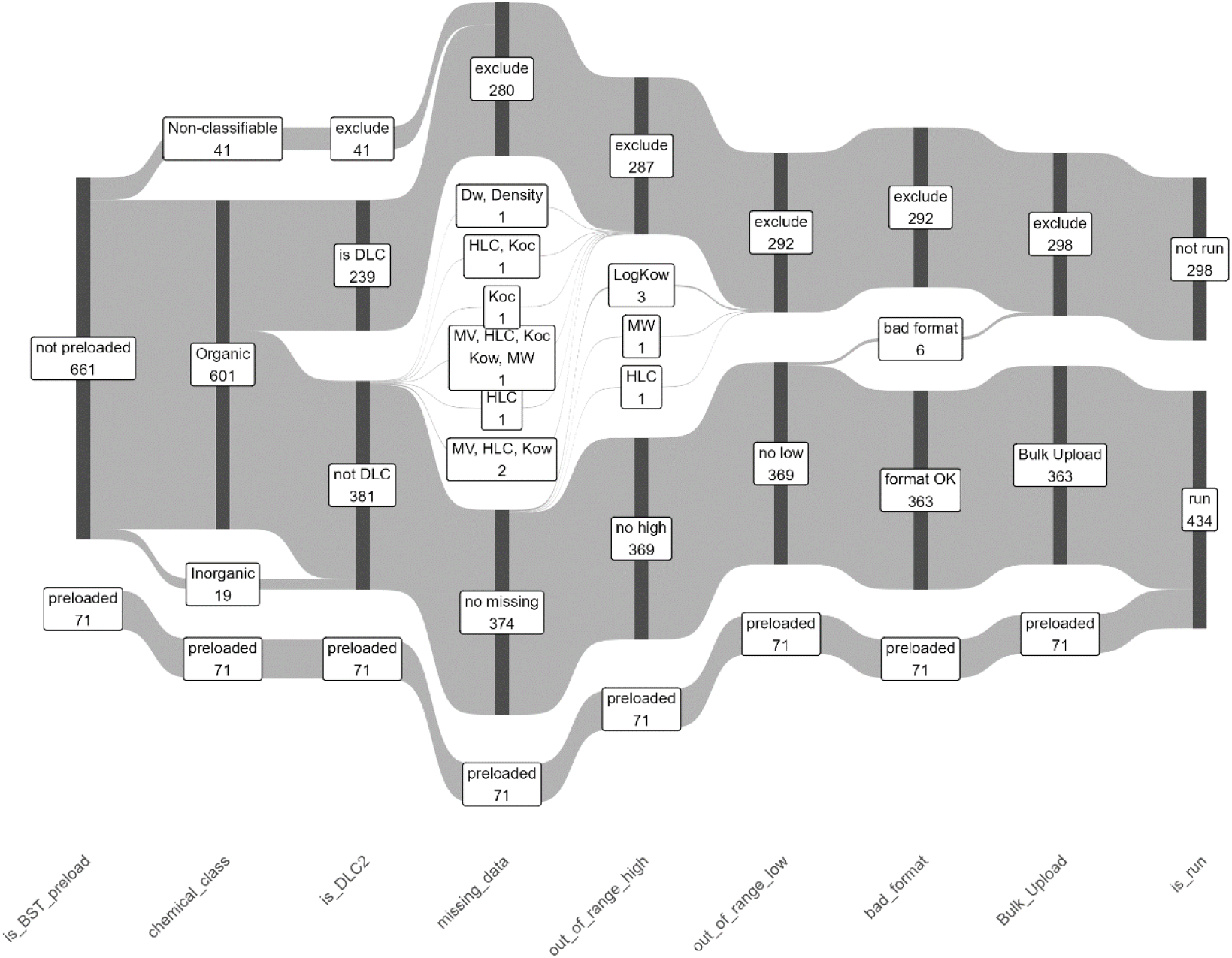
Sankey diagram of data filtering steps for 727 DTXSIDs in the curated list of chemicals detected in biosolids. The flow proceeds from left to right across the stages that are labeled along the bottom. “is_BST_preload”: whether the DTXSID corresponds to a CASRN already in the BST. “chemical_class”: Is the chemical organic, inorganic, or non-classifiable (usually UVCB). “is_DLC2”: Whether the chemical is dioxin-like or not. “missing_data”: Whether the chemical is still missing values for any of the required BST parameters after the hierarchical process described in the main text. “out_of_range_high” and “out_of_range_low”: Whether any of the required BST parameter values were more than 1% out of range on the high or low end, respectively. “bad_format”: Whether the DTXSID is on the list of chemicals that trigger a bug in the BST. “Bulk_Upload”: whether chemical info is to be imported using the Bulk Upload Tool (chemicals which passed all previous stages). “is_run”: whether the chemical is included in BST simulation runs (the combination of bulk-uploaded chemical and preloaded chemicals).

Of the chemicals that could be classified, 239 were excluded because they were DLCs. Seven chemicals were missing values for one or more physico-chemical properties. Three more chemicals were excluded because their log K_ow_ values were more than 1% greater than the high end of the BST operating range (i.e., 12), and one was excluded because its molecular weight was above the high end of the BST operating range (i.e., 1000 g/mol). Six more chemicals were excluded because, during testing of the autoBST, it was discovered that they trigger a bug in the BST that causes it to crash. (A reliable fix for this bug could not be ascertained, even in consultation with BST developers.) This left 363 chemicals to be uploaded using the Bulk Upload Tool. Combined with the 71 pre-loaded chemicals, the BST was able to make exposure predictions for 434 total chemicals.

When chemicals with no NSSS concentration data were excluded, the BST was able to make predictions for 234 chemicals (Figure 2). 256 of the 727 chemicals were excluded because they did not have NSSS mean or 95th percentile concentrations. 66 more chemicals were pre-loaded into the BST. Fourteen chemicals were excluded because they could not be classified as organic or inorganic. 221 more chemicals were excluded because they were DLCs. Of the remaining 175 chemicals, none were missing physico-chemical data; however, three were excluded because their log K_ow_ values were more than 1% greater than the high end of the BST operating range (i.e., 12). Four more chemicals were excluded because they trigger the aforementioned BST bug. Combined with the 66 pre-loaded chemicals, the BST was able to make exposure predictions for 234 total chemicals.

**Figure 2.**
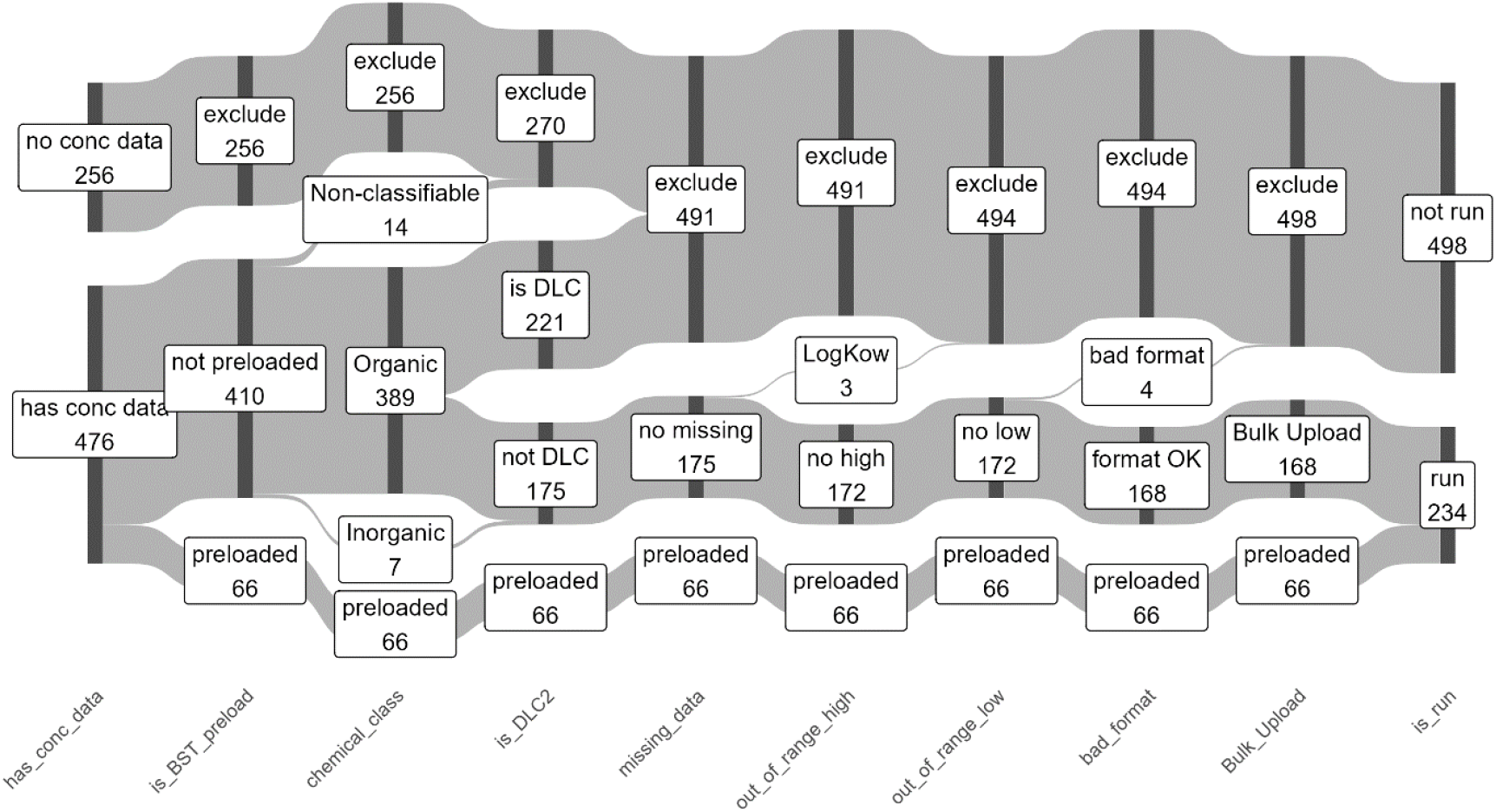
Sankey diagram of data filtering steps for 727 DTXSIDs in the curated list of chemicals detected in biosolids. As for Figure 1, but Includes an additional data filtering step at the beginning: whether chemicals have mean or 95^th^ percentile concentration values in the NSSS.

The largest source of exclusions was dioxin-like compounds (DLCs) (dioxins, furans, and PCBs), which are excluded not because of a lack of data, but because the guidance for use of the BST excludes this class of chemicals. Only seven chemicals were excluded because they were missing chemical-specific data. For the vast majority of substances detected in biosolids, autoBST was successful at locating chemical-specific parameter values from publicly available databases.

The Microsoft Excel files generated by autoBST for the Bulk Upload Tool for each scenario are provided in Supplementary Tables 1-3. The output of the BST is provided in Supplementary Tables 4-6. BST-predicted oral average daily doses (ADD) for an adult farmer spanned a wide range of values across chemicals (Figure 3). For simulations made using NSSS mean and 95^th^ percentile concentrations, the range was approximately 1 × 10^-10^ – 1 × 10^5^ mg/kg/day, with mode around 1 × 10^-4^ mg/kg/day (100 ng/kg/day). For simulations made using 1 ppb concentrations, the range was somewhat narrower, approximately 1 × 10^-12^ – 1 mg/kg/day, with mode around 1 × 10^-6^ mg/kg/day (1 ng/kg/day).

**Figure 3.**
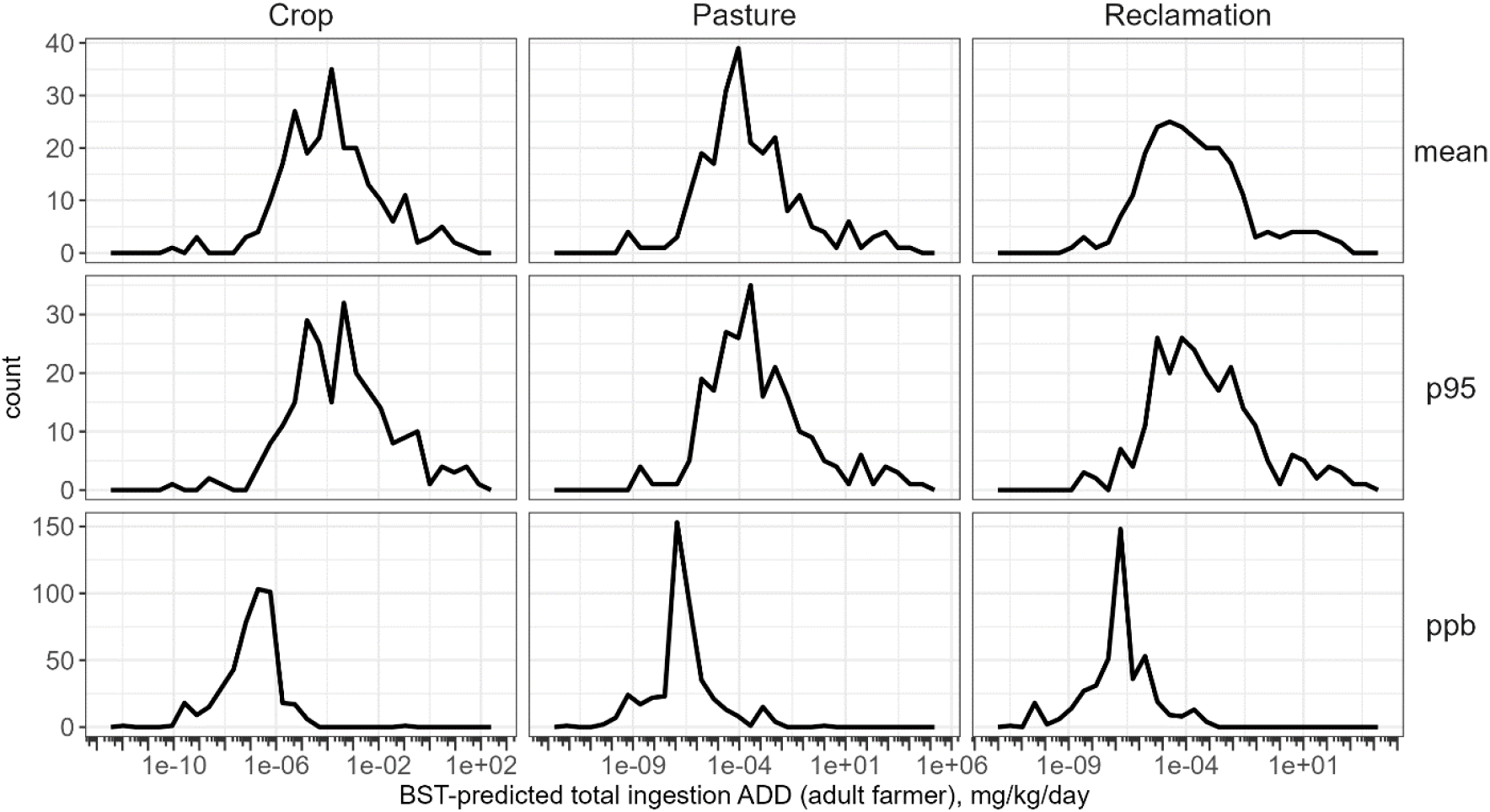
Frequency polygons (similar to histograms) of adult farmer ADD (mg/kg/day), by LAU scenario (columns) and by which biosolids concentration was used as input to simulations (rows): NSSS mean (“mean”), NSSS 95^th^ percentile (“p95”), or 1 ppb (“ppb”). Count (vertical axis) represents number of chemicals. Note log_10_ scale on horizontal axis and note differing scales on vertical axis for each row.

Chemicals were ranked by BST-predicted oral average daily dose (ADD) for an adult farmer, under each of the three land-application use (LAU) scenarios, for each biosolids concentration considered (NSSS mean, NSSS 95^th^ percentile, and 1 ppb) (Supplementary Table 7). For BST predictions based on NSSS mean biosolids concentrations, the top ten chemicals by the adult farmer ADD for the crop LAU scenario were coprosterol, (3alpha,5beta)-cholestan-3-ol, calcium, (+)-dihydrocholesterol, cholesterol, iron, (-)-beta-sitosterol, stigmasterol, squalene, and stigmastan-3beta-ol. For BST predictions based on NSSS 95th percentile biosolids concentrations, the top ten chemicals were the same, except that (-)-beta-sitosterol moved down two places. (In both cases, the top ten rankings for the pasture and reclamation LAU scenarios were similar.) These substances were all in the top 20 for highest NSSS mean and 95th percentile biosolids concentrations (except squalene, which was number 22). The occurrence of sterols at high levels in biosolids is expected; sterols occur naturally in animal feces as the end product of cholesterol metabolism— in fact, they are often used as a biomarker to detect human or animal fecal contamination of water (Fulke et al. 2025). Sterols are lipophilic, so the potential for higher exposures is reasonable.

NSSS biosolids concentration explained about half of variability in predicted adult farmer ADD in a log_10_-log_10_ regression for all three LAU scenarios, both for the NSSS mean and NSSS 95^th^-percentile concentration scenarios (Figure 4). (Crop scenario: R^2^ 0.651 for NSSS mean, 0.673 for NSSS 95th percentile. Pasture scenario: R^2^ 0.508 for NSSS mean, 0.530 for NSSS 95th percentile. Reclamation scenario: R^2^ 0.497 for NSSS mean, 0.528 for NSSS 95^th^ percentile.) Higher adult farmer ADDs were also correlated with higher log K_ow_ values for the pasture and reclamation scenarios, but not for the crop scenario. When log K_ow_ was added to the regression as a second term, its coefficient was not statistically significant for the crop scenario, and R2 values did not improve (crop R^2^ 0.663 for NSSS mean, 0.685 for NSSS 95th percentile), but its coefficient was statistically significant (and positive) for the pasture and reclamation scenarios, and R^2^ values improved (pasture R^2^ 0.711 for NSSS mean, 0.723 for NSSS 95^th^ percentile; reclamation R^2^ 0.777 for NSSS mean, 0.792 for NSSS 95^th^ percentile). The difference in relation between log ADD and log K_ow_ across land-application scenarios may be explained by the different exposure pathways considered by the BST exposure models in each scenario. Pasture and reclamation scenarios include dietary exposures only from beef and milk from exposed animals and fish from an exposed pond, whereas the crop scenario also includes dietary exposures from produce grown in contaminated soil (USEPA, 2023). The autoBST workflow estimates beef and milk bioconcentration factors based on log K_ow_. Beef and milk constitute a larger proportion of dietary exposures in the pasture and reclamation scenarios than in the crop scenario, so it makes sense that exposure would depend more strongly on log Kow in pasture and reclamation scenario.

**Figure 4.**
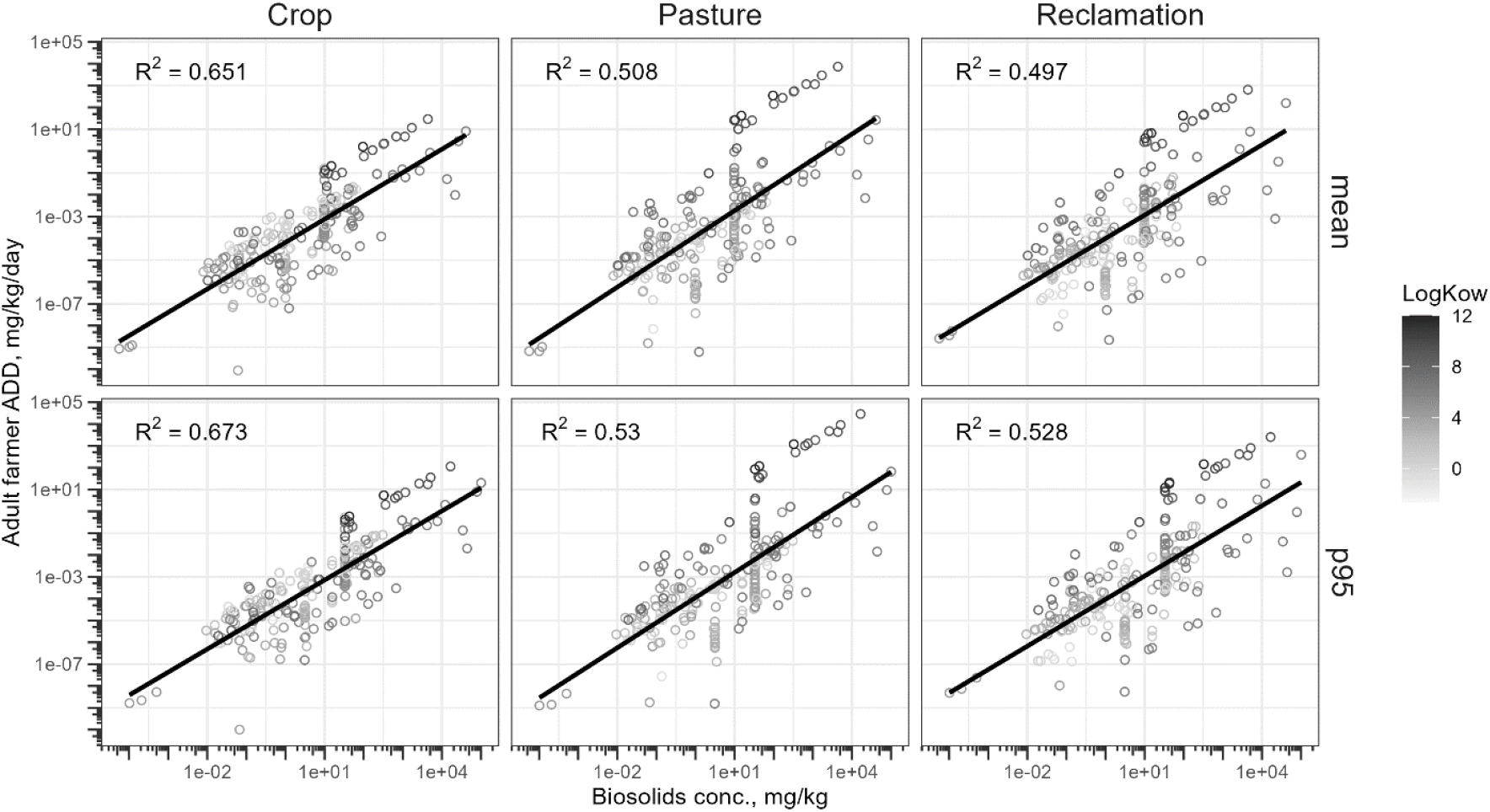
Adult farmer ADD vs. biosolids concentration by LAU scenario (columns) and by which NSSS concentration was used (rows): mean or 95^th^ percentile (“p95”). Each circle represents one chemical. Circles are grayscale-coded by log Kow: darker gray represents higher log K_ow_. Black lines represent best-fit line for log_10_ ADD regressed on log10 biosolids concentration in each panel. R^2^ for each linear regression is shown in upper left corner of each panel. Note that log K_ow_ was not included in the linear regressions shown here; see text for more details.

The 1 ppb biosolids concentration scenario results were informative about the exposure potential of chemicals that did not have NSSS concentration information. The top ten chemicals by adult farmer ADD for the crop scenario included mercury (divalent), squalene, dimethyl phthalate, n-tetracosane, selenium (VI), docosane, di-n-octyl phthalate, eicosane, 4,4’-methylenebis(2,6-di-t-butylphenol), and (-)-beta-sitosterol. (The top ten chemicals for pasture and reclamation scenarios were similar: coprosterol, (3alpha,5beta)-cholestan-3-ol and (+)-dihydrocholesterol joined the top ten, and dimethyl phthalate and selenium (VI) dropped out of the top ten; for the pasture scenario, eicosane also dropped out of the top ten.) Mercury (divalent) and selenium (VI) are chemicals pre-loaded into the BST; their chemical parameters were not determined according to autoBST, but were pre-determined (see the BST User Guide for more information about parameters for these chemicals). Selenium (VI) is considered bioaccumulative in fish (the BST uses a level-3 BCF of 490 and a level-4 BCF of 1700), and divalent mercury is considered extremely bioaccumulative in fish (the BST uses level-3 BCF 1.6 × 106 and level-4 BCF 6.8 × 106), which may explain the high rankings of these substances.

## Discussion

AutoBST is an automated, reproducible workflow that uses information from publicly available cheminformatics databases and tools to rapidly parameterize a model to predict biosolids exposures. AutoBST’s data collection and internal checks are rapid and thus lend themselves to higher-throughput analysis of hundreds or thousands of chemical contaminants of interest. A case study has been presented wherein autoBST was used to parameterize the BST for hundreds of chemicals with existing biosolids monitoring data from National Sewage Sludge Surveys. However, autoBST is not limited to use with the chemical list included in this case study. Because of its modular, programmatic structure and its integration of chemical data APIs, autoBST can be applied — with little or no modification — to parametrize and run the BST to predict exposures for large numbers of new chemicals as additional biosolids occurrence data become available, for example through the biennial review process required under the Clean Water Act, or through new approaches such as non-targeted screening (Newmeyer et al. 2024).

Because autoBST pulls chemical data from public databases, it is both limited and strengthened by the availability of data in its sources. For example, because autoBST currently relies on ClassyFire to categorize chemicals as organic or inorganic, eleven chemicals had to be excluded from the case study because ClassyFire did not have a classification available, even though they had defined structures.

Future improvements to autoBST will include making organic/inorganic categorizations using available structural identifiers, without relying on an outside service. However, autoBST is also able to make use of new data as they are made available through ongoing updates to the source databases, because it uses APIs to programmatically pull data from public databases rather than relying on static data dumps.

In the case study presented here, the BST was used to predict average oral daily doses resulting from land applications of biosolids. However, potential for exposure does not necessarily imply potential for hazard or risk. To predict non-cancer hazard and cancer risk for humans, as well as risks for ecological receptors, the BST includes the capability to compare predicted exposures to user-provided toxicity reference values. While selection of the most appropriate toxicity reference values is beyond the scope of autoBST in its current state, autoBST could easily be extended to automate the process of selecting reference values by drawing on publicly-available databases of toxicity and hazard information such as ToxValDB (Judson 2018).

Our results indicate that, for 234 chemicals with monitoring data from the National Sewage Sludge Surveys, biosolids concentrations explained approximately half of the variability in BST-predicted exposures. In other words, biosolids occurrence alone is not the whole story of exposure: modeling environmental fate and transport remains important. The autoBST workflow provides much-needed capability to compile the chemical-specific data needed to model fate and transport in the BST in a manner suitable for automated, high-throughput uses, thereby enabling researchers, risk assessors, and other decision-makers to quickly and easily to predict human exposures from chemical contaminants in biosolids.

## Supporting information

BST Bulk Upload

BST Bulk Chemical Import Tool

autoBST workflow files

## Acknowledgments

The authors are deeply grateful to the US EPA Office of Water Biosolids Team for expert advice and user testing, including David Tobias, Tess Richman, Lisa Weber, and Sophie Greene. The authors also gratefully acknowledge the team at RTI International responsible for developing the Biosolids Screening Tool and its bulk-upload capabilities, including Ted Lillys, Anne Lutes, and Donna Womack.

## Supplementary Material

BST_BulkUpload.mdb: Microsoft Access implementation of the Biosolids Screening Tool, in a version that has the capability for bulk-uploading chemicals.

BST_BulkChemicalImportTool.xlsm: Microsoft Excel workbook, providing the template used by autoBST to produce output suitable for bulk upload into the BST.

autobst.zip: A ZIP archive of the autoBST code and data files, also available on GitHub at https://github.com/USEPA/CompTox-ExpoCast-autoBST.

Supplementary Table 1: AutoBST workflow output (Excel workbook of BST parameters suitable for uploading using the Bulk Upload Tool) when biosolids concentrations are set to National Sewage Sludge Survey mean concentrations.

Supplementary Table 2: AutoBST workflow output (Excel workbook of BST parameters suitable for uploading using the Bulk Upload Tool when biosolids concentrations are set to National Sewage Sludge Survey 95th percentile concentrations.

Supplementary Table 3: AutoBST workflow output (Excel workbook of BST parameters suitable for uploading using the Bulk Upload Tool) when biosolids concentrations are set to 1 ppb for all chemicals.

Supplementary Table 4: BST simulation results when biosolids concentrations are set to National Sewage Sludge Survey mean concentrations (i.e., when Supplementary Table 1 is used as input). See BST User Guide for details.

Supplementary Table 5: BST simulation results when biosolids concentrations are set to National Sewage Sludge Survey 95th percentile concentrations (i.e., when Supplementary Table 2 is used as input). See BST User Guide for details.

Supplementary Table 6: BST simulation results when biosolids concentrations are set to 1 ppb for all chemicals (i.e., when Supplementary Table 3 is used as input). See BST User Guide for details.

Supplementary Table 7: Chemical rankings by BST-predicted oral average daily dose (ADD) for an adult farmer, under each of the three land-application use (LAU) scenarios (Crop, Pasture, and Reclamation), for each biosolids concentration scenario considered (NSSS mean, NSSS 95th percentile, and 1 ppb).

