## Supplementary material for "A Cheminformatics Workflow for Higher-throughput Modeling of Chemical Exposures from Biosolids": autoBST workflow files: BST BulkLoader Workflow Manuscript.Rmd

````
---
title: "autoBST"
author: "Paul Kruse and Caroline Ring"
date: "2025-03-24"
output: html_document
---

```{r setup, include=FALSE}
knitr::opts_chunk$set(echo = TRUE)
```

# Setup 

Ensure your working directory in R is the directory where this R Markdown document lives (using commands `getwd()` to check the current working directory, and `setwd()` to change it if needed).

Load necessary packages. All are available on CRAN except for `ggsankey`.

```{r}
library(tidyverse)
library(data.table)
library(httk)
library(ctxR)
library(ggplot2)
library(ggsankey) #use devtools::install_github("davidsjoberg/ggsankey") to install
#or if that doesn't work, clone the github repo yourself and use devtools::install_local() to install from the locally-cloned repo
```

## Register your CTX API key

To use the `ctxR` package to access EPA's Computational Toxicology and Exposure (CTX)
APIs, an API key is required. Before continuing, please use `ctxR::register_ctx_api_key()` to register your API key for this R session: in the following command, replace `<YOUR API KEY>` with your actual API key, and then evaluate at the R command line. Please do not paste your actual API key into this document, or anywhere where it might inadvertently be shared with other people.

```{r, eval = FALSE}
ctxR::register_ctx_api_key(key = '<YOUR API KEY>',
                           write = TRUE)
```

# Bulk Uploader Tool template sheets

Load in the template sheets from the Bulk Uploader Tool: templates for organic chemicals, inorganic chemicals, and ecological data associated with each chemical.

If you open the template file `data/BST_BulkChemicalImportTool.xlsm` in Microsoft Excel, you will find a good deal of information and context about the various sheets in the template. You will also find instructions to use the template file by manually inputting chemical information into the spreadsheet and then clicking buttons to run certain macros. You can ignore these instructions if you are using autoBST. autoBST reproduces what would be contained in the file after manual input of data and running all macros; you won't need to do anything. autoBST only uses `data/BST_BulkChemicalImportTool.xlsm` as a template to format its output. 

```{r}
organic_template <- readxl::read_excel(path = 'data/BST_BulkChemicalImportTool.xlsm',
                                       sheet = 'Template_Chem_Org',
                                       range = 'A4:M60')

inorganic_template <- readxl::read_excel(path = 'data/BST_BulkChemicalImportTool.xlsm',
                                       sheet = 'Template_Chem_Inorg',
                                       range = 'A4:M48')

eco_template <- readxl::read_excel(path = 'data/BST_BulkChemicalImportTool.xlsm',
                                       sheet = 'Template_Eco',
                                       range = 'A4:M58')
```

# Biosolids concentration data

Load in the biosolids concentration data. Note units: the Bulk Uploader Tool requires measurements in ug/g (equivalent to mg/kg).

```{r}
NSSS <- readxl::read_excel(path = 'data/Richman_et_al_2022_SI_Table4_Biosolids Concentration Data from NSSSs.xlsx',
                                         sheet = 'Chemical Concentration Data',
                                         range = 'A2:AJ486')

#convert to data.table
NSSS <- data.table(NSSS)

#Rename columns to reflect years
#1988 
setnames(NSSS,
         names(NSSS)[5:15],
         paste("1988",
               names(NSSS)[5:15]))
#2001
setnames(NSSS,
         names(NSSS)[16:27],
         paste("2001",
               names(NSSS)[16:27]))
#2009
setnames(NSSS,
         names(NSSS)[28:36],
         paste("2009",
               names(NSSS)[28:36]))

#remove the "...[number]" suffixes that were originally added during data import
setnames(NSSS,
         names(NSSS),
         gsub(pattern = "\\.\\.\\.\\d+$",
              replacement = "",
              x = names(NSSS)))

#also read in nutrients sheet
NSSS2 <- readxl::read_excel(path = 'data/Richman_et_al_2022_SI_Table4_Biosolids Concentration Data from NSSSs.xlsx',
                                         sheet = 'Nutrient Concentration Data',
                                         range = 'A2:U5')

NSSS2 <- data.table(NSSS2)

#Rename columns to reflect years
#1988 
setnames(NSSS2,
         names(NSSS2)[5:11],
         paste("1988",
               names(NSSS2)[5:11]))
#2009
setnames(NSSS2,
         names(NSSS2)[13:21],
         paste("2009",
               names(NSSS2)[13:21]))

NSSS2[, `...12` := NULL]

#remove the "...[number]" suffixes that were originally added during data import
setnames(NSSS2,
         names(NSSS2),
         gsub(pattern = "\\.\\.\\.\\d+$",
              replacement = "",
              x = names(NSSS2)))

setnames(NSSS2,
         names(NSSS2),
         gsub(x = names(NSSS2),
              pattern = "1/2 ML",
              replacement = "1/2 MRL",
              fixed = TRUE))

setnames(NSSS2,
         "MasterList Chemical Name",
          "Chemical Name Reported in Biennial Reviews and Sewage Sludge Surveys")

NSSS <- data.table::rbindlist(list(NSSS,
                       NSSS2),
                  use.names = TRUE,
                  fill = TRUE)
```

# ClassyFire chemical classifications

ClassyFire classifications can be queried from the ClassyFire API, but the InCHiKey is required. Get the InCHiKey for each DTXSID in the NSSS data, using the `ctxR` function `ctxR::get_chemical_details_batch()`.

```{r}
biosolids_curated <- readxl::read_excel("data/Richman_et_al_2022_SI_Table1_Master List of Chemicals Detected in Biosolids.xlsx",
                                        "Biosolids Master List",
                                        "A1:D728")
#remove extra blank row
biosolids_curated <- biosolids_curated[2:nrow(biosolids_curated), ]

#also read in nutrients
biosolids_curated2 <- readxl::read_excel("data/Richman_et_al_2022_SI_Table1_Master List of Chemicals Detected in Biosolids.xlsx",
                                        "Nutrients Detected in Biosolids",
                                        "A1:D8")
#remove extra blank row
biosolids_curated2 <- biosolids_curated2[2:nrow(biosolids_curated2), ]

biosolids_curated <- data.table::rbindlist(list(biosolids_curated,
                                                biosolids_curated2),
                                           use.names = FALSE)

chem_details <- ctxR::get_chemical_details_batch(DTXSID = biosolids_curated$DTXSID)
```

Source helper functions to query the ClassyFire API.

```{r}
source("classyfireAPIs.R")
```

Because querying the ClassyFire API can be slow, we highly recommend only running this section once, and *saving* the result to re-load later. The ClassyFire API is limited to a maximum of 12 requests per minute; with 727 unique InCHiKeys in the biosolids curated list, that translates to approximately 61 minutes. The results of the query have been pre-saved in the `data/` folder.

```{r, eval = FALSE}
system.time(
  chem_classes <- classify_inchikeys(inchikeys = chem_details[!is.na(inchikey),
                                                      unique(inchikey)])
)

chem_classes <- as.data.table(chem_classes)

saveRDS(chem_classes,
        "data/biosolids_curated_ClassyFire.Rds")
```

Read in the saved ClassyFire classifications.

```{r}
chem_classes <- readRDS("data/biosolids_curated_ClassyFire.Rds")
```

# BST pre-loaded chemicals

Load a list of chemicals that are pre-loaded into the BST (these will be removed from analysis). These are identified by CASRN (but stripped of hyphens).

```{r}
previously_loaded_casrns <- data.table::fread("data/BST previously loaded CASRNs.csv")
```

# List of CASRNs whose data is known to crash the BST

```{r}
bad_chems <- fread("data/Bad format chemicals BST 2023_10_6.txt",
                   col.names = c("casrn_stripped", "chemical_name"))
```


# Sources of physical-chemical data other than the Comptox Chemicals Dashboard

Load in a table of density values for elements in the periodic table. This was sourced from https://pubchem.ncbi.nlm.nih.gov/periodic-table/density/.

```{r}
element_densities <- read.csv('data/PubChemElements_all.csv')
```

# Bio-transfer factors

## National Bioaccumulation Factors

Load in table of National Bioaccumulation Factors

```{r}
reported_BAF <- readxl::read_excel(path = 'data/national-bioaccumulation-factors-supplemental-information.xlsx',
                                   sheet = 'National BAFs',
                                   range = 'A2:T99')
reported_BAF <- data.table::as.data.table(reported_BAF)
names(reported_BAF)[16:18] <- c('National BAF TL3', 'National BAF TL4', 'Alternative BCF')
reported_BAF[, CASRN := gsub(pattern = '-', replacement = '', x = `CAS Number`)]
# Remove duplicate data row (only differ in name)
reported_BAF <- reported_BAF[-11,]
```

Functions to predict biotransfer factors:

## Bio-transfer factors for beef and milk

The first is $\text{BTF}_\text{fat}$, which is used to calculate the quantities $\text{BCF}_\text{beef}$ and $\text{BCF}_\text{milk}$.

These equations are as implemented in the BST. THey are here because the BST may be inconsistent in whether it properly recalculates the transfer factors. So we do it here.

```{r}
btf_fat <- function(log_kow){
  # log_kow must be between -0.67 and 8.2. If it exceeds either bound, it is set
  # to the appropriate bound it exceeded
  log_kow <- min(8.2, max(-0.67, log_kow))
  
  log_btf <- -0.099*log_kow^2 + 1.07*log_kow - 3.56
  return(10^(log_btf))
}

btf_beef <- function(log_kow){
  btf_fat_value <- btf_fat(log_kow = log_kow)
  return(0.19*btf_fat_value)
}

btf_milk <- function(log_kow){
  btf_fat_value <- btf_fat(log_kow = log_kow)
  return(0.04*btf_fat_value)
}

bcf_beef <- function(log_kow){
  btf_beef_value <- btf_beef(log_kow = log_kow)
  return(12.27*btf_beef_value)
}

bcf_milk <- function(log_kow){
  btf_milk_value <- btf_milk(log_kow = log_kow)
  return(20.7*btf_milk_value)
}
```

## Soil-to-plant bioaccumulation/bioconcentration factors

Next we calculate soil to plant BAFs/BCFs

```{r}
bioconcentration_factor_vegetation_org <- function(log_kow){
  # The allowable range for log_kow is 1.15 to 9.35. If it exceeds either bound,
  # it is set to the bound it exceeds
  log_kow <- min(9.35, max(1.15, log_kow))
  
  log_br <- 1.588 - 0.578*log_kow
  return(10^log_br)
}
```

## Air-to-plant bioconcentration factor

Next we calculate the air-to-plant BCF.

```{r}
vol_air_to_leaf <- function(log_kow, HLC){
  # The allowable range for log_kow is 1.2 to 8.2, and for HLC is 6.1E-8 to 
  # 8.3E-3. If either bound is exceeded, the value is assigned to the 
  # associated bound.
  log_kow <- min(8.2, max(1.2, log_kow))
  HLC <- min(8.3E-3, max(6.1E-8, HLC))
  
  # Set constant values
  R_ideal <- 8.205E-5
  TEMP <- 298.1
  log_bvol <- 1.065*log_kow - log10(HLC/(R_ideal*TEMP)) - 1.654
  return(10^log_bvol)
}

air_to_leaf_biotransfer_factor <- function(log_kow, HLC){
  # Set constant values
  rho_air <- 1.19
  f_water <- 0.85
  rho_plant <- 770
  
  B_vol <- vol_air_to_leaf(log_kow = log_kow, HLC = HLC)
  
  B_v <- (rho_air * B_vol)/((1 - f_water)*rho_plant*100)
  
  return(B_v)
}
```

## Root concentration factor

Now we calculate the root concentration factor.

```{r}
root_concentration_factor_org <- function(log_kow) {
  # The allowable range for log_kow is -0.57 to 8.2. If it exceeds either bound,
  # it is set to the bound it exceeds.
  log_kow <- min(8.2, max(-0.57, log_kow))
  
  if (log_kow < 2){
    log_rcf_offset <- 0.77*log_kow - 1.52
    rcf_offset <- 10^log_rcf_offset
    rcf <- rcf_offset + 0.82
  } else {
    log_rcf <- 0.77*log_kow - 1.52
    rcf <- 10^log_rcf
  }
  
  return(rcf)
}
```

# Physical-chemical properties

Most physical-chemical properties will be gathered from the CCD. However, there are a few that either may not be available for all chemicals from the CCD, or are known to not be available at all from the CCD.

## Organic carbon partition coefficient Koc

The Organic Carbon Partition Coefficient is one of the required inputs for the BST. As with other physico-chemical properties, we first try to gather this data from the Dashboard using ctxR. If a value for the Organic Carbon Partition Coefficient is not available, we can calculate it using the methods from SimpleTreat.

```{r}
# This takes in the chemical type (acid, base, or neutral), the pK data, and the Kow data and uses the
# SimpleTreat 4.0 method for calculating the Koc value.

calc_koc <- function(DTXSID,
                     pH = 7,
                     Kow){
  
  chemical_type <- get_ionization(DTXSID = DTXSID,
                           pH = pH)
  
    pKa_data <- get_pka(DTXSID = DTXSID)
    
    if(length(pKa_data[[1]])> 0 &
       length(pKa_data[[2]]) > 0){
    if(is.finite(pKa_data[[1]]) & 
       is.finite(pKa_data[[2]])){
       #If there is both a pKa_acidic and a pKa_basic,
    #we use the one matching the predominant species at pH 7.0
    #if it's an acid, then we use pKa_a
    #if it's a base, then we use pKa_b
    if(chemical_type %in% "acid"){
      pKa <- pKa_data[[1]]
    }else if(chemical_type %in% "base"){
      pKa <- pKa_data[[2]]
    }
      #if we only have one pKa value, then use it as-is
    }else if(is.finite(pKa_data[[1]]) & 
       !is.finite(pKa_data[[2]])){
      pKa <- pKa_data[[1]]
    }else if(!is.finite(pKa_data[[1]]) & 
       is.finite(pKa_data[[2]])){
      pKa <- pKa_data[[2]]
    }else{
      #if we don't have a pKa value, then use NaN
      #in this case it will be neutral and pKa will not be used anyway
      pKa <- NaN
    }
    }else{
      pKa <- NA_real_
    }
  
  Koc <- NA_real_
if(chemical_type %in% "neutral"){
  #the following Koc is for primarily hydrophobic chemicals
  print('Neutral')
  Koc <- 1.26* Kow^(0.81)
  return(Koc)
  # "Unlike rules for estimating the bio-concentration potential from Kow, which is
  # limited by the molecular weight of 700 (TGD, 2003), the applicability range of
  # the relationship between Kp and Kow is wide. Nevertheless, special structural
  # properties related to, for example, amphiphilic substances (surfactants) or
  # dissociating substances (organic acids, bases) may lead to multiple equilibrium
  # processes." -- from SimpleTreat 4.0 doc, PDF page 24/marked page 23
}else{ #for acids and bases
  #default pH = 7
  pH <- 7
  if(chemical_type == "acid"){
    gamma <- 1
    Fn <- 1/(1 + 10^(gamma * (pH - pKa)))
    #Kaw <- Kaw * Fn
    Fn_opt <- 1/(1 + 10^(gamma * ((pH - 0.6) - pKa)))
    if(is.na(Koc)){
      Koc <- Fn_opt*10^(0.54 * log10(Kow)+1.11) +
        (1-Fn_opt)*10^(0.11*log10(Kow) + 1.54)
    }
  }else if(chemical_type %in% "base"){
    gamma <- -1
    Fn <- 1/(1 + 10^(gamma * (pH - pKa)))
    Dow <- Fn * Kow
    
    Koc <- 10^(0.31*log10(Dow)+2.78)
    
  }
 
}
    return(Koc) 
}
```

### pKa

Because pKa is required in order to predict Koc, we also need a way to get pKa. pKa is currently not available via CTX APIs. Therefore, we get pKa values by running OPERA. The OPERA predictions were saved.

```{r}
opera <- data.table::fread('data/biosolids_chemicals-txt_OPERA2.9Pred.csv')

#remove any DTXSIDs for which predictions could not be made
opera <- opera[!(FoundBy %in% c("NotFound",
                                "Error: Inorganic")), ]

```

(Incidentally, all of the NSSS DTXSIDs without OPERA predictions are inorganic or UVCB. OPERA made pKa predictions for all of the NSSS organic chemicals.)


This is a helper function to retrieve the OPERA-predicted pKa values. OPERA predicts two pKa values: an acidic pKa and a basic pKa. (These are not the same as pKa and pKb.)

```{r}
get_pka <- function(DTXSID){
  pka_pred <- opera[MoleculeID %in% c(DTXSID),
                    .(pKa_a_pred, pKa_b_pred)]
  return(pka_pred)
}
```

### Chemical type: acid or base

To predict Koc, we need to know whether the chemical is acid, base, or neutral at the specified pH (in this case, pH 7.0). To predict ionization, we can use the method implemented in the `httk` R package. This is a helper function to get the majority ion species in solution at the specified pH, and therefore to tell whether the chemical is acting as an acid, a base, or neutral.

```{r}
get_ionization <- function(DTXSID,
                           pH = 7){
  
  opera_pka <- get_pka(DTXSID)
  #if opera_pka is an empty data.table (0 rows) then it means OPERA did not make a prediction for this DTXSID. Substitute NULL, because httk::calc_ionization() uses NULL as a special value to mean "no value supplied," and uses NA as a special value to mean "no ionization occurs."
  if(nrow(opera_pka) %in% 0){
    pka_pred <- list(pKa_a_pred = NULL,
                     pKa_b_pred = NULL)
  }
  
   chemical_type <- tryCatch(
      {
      which(httk::calc_ionization(pH = 7,
                                  pKa_Donor = opera_pka[[1]],
                                  pKa_Accept = opera_pka[[2]]) > .5)
      }, error = function(cond){
        message(cond$message)
        return(list())
      })

      chemical_type <- intersect(names(chemical_type), c('fraction_neutral', 'fraction_negative',
                                                         'fraction_positive', 'fraction_zwitter'))
     
      if (is.null(chemical_type)){
        chemical_type <- ''
      } else if (chemical_type %in% 'fraction_neutral'){
      chemical_type <- 'neutral'
    } else if (chemical_type %in% 'fraction_negative'){
      chemical_type <- 'acid'
    } else if (chemical_type %in% 'fraction_positive'){
      chemical_type <- 'base'
    } else {
      chemical_type <- 'zwitter'
    }
      
      return(chemical_type)
}
```

## Air and water diffusivity constants

To determine the air and water diffusivity coefficients, we use two functions developed to estimate these quantities. These can be found at [https://www3.epa.gov/ceampubl/learn2model/part-two/onsite/estdiffusion.html].

We describe the equations and then code them as functions.

The air diffusivity coefficient is given by $$D_{\text{air}} = \frac{0.001T^{1.75}M_r^{\frac{1}{2}}}{\left(P\left(V_A^{\frac{1}{3}} + V_B^{\frac{1}{3}}\right)^2\right)}$$

Where $D_{\text{air}}$ is the diffusion coefficient in cm^2/sec, $T$ is the temperature in K, $M_r$ is the function of molecular weight $$M_r = \frac{(M_A + M_B)}{M_AM_B}$$ with $M_A$ the molecular weight of air, approximately 28.97 b/mol, and $M_B$ is the molecular weight of the compound of interest. Additional parameters are $V_A$ the diffusion volume of air (approximately 20.1 cm^3/mol), $V_B$ the diffusion volume of the compound of interest, and $P$ is the pressure in atm.

```{r}
# This assumes operating at 1 atm and 25 celsius
air_diffusivity <- function(Temp = 298.15,
                            M_a = 28.97,
                            M_b = NULL,
                            V_a = 20.1,
                            V_b = NULL,
                            P = 1){
  M_r <- (M_a + M_b)/(M_a*M_b)
  D_air <- 0.001 * Temp^{1.75}*M_r^{0.5}/(P*(V_a^{1/3} + V_b^{1/3})^2)
  return(D_air)
}
```

The water diffusivity coefficient is given by $$D_{\text{water}} = \frac{13.26 \times 10^{-5}}{h^{1.14} (V^{'}_B)^{0.589}}$$
In this equation, $h$ is the viscosity of water and $V^{'}_B$ is the LaBas estimate of diffusion volume.

```{r}
# This assumes operating at 1 atm and 25 celsius
water_diffusivity <- function(h = 0.8904,
                              V_lebas){
  D_water = (13.26E-5)/(h^{1.14}*(V_lebas)^{0.589})
  return(D_water)
}
```

Next diffusivity in water for inorganic.

```{r}
diff_water_inor <- function(mol_weight, density) {
  # Set constant values
  TEMP <- 25
  diffusion <- (1.518E-4)*((TEMP + 273.16)/298.16)*(mol_weight/density)^(-0.6)
  return(diffusion)
}
```


Convert the concentration in dry-mass biosolids to the concentration in sewage sludge, assuming that solids are 40% of the sludge by mass.

```{r}
ctpwaste <- function(ctp_waste_dry){
  # Set constant value. Solids is assumed to be 40 percent.
  solids = 40
  return(ctp_waste_dry*solids/100)
}
```

# Function to generate organic template table for a chemical

The following function implements the procedure for populating the organic template table by querying the Dashboard and/or drawing from the datasets and/or prediction functions above.

```{r}
generate_organic_entry <- function(DTXSID,
                                   biosolids_concentration = 1){
  #print(DTXSID)
  # Get chemical details
  chemical_details <- ctxR::get_chemical_details(DTXSID = DTXSID)
  
    if(length(chemical_details) == 0){
    chemical_details <- list(casrn = NA_character_,
                             preferredName = NA_character_,
                             monoisotopicMass = NA_real_)
  }
  
  # Get chemical info
  chemical_info <- data.table(ctxR::get_chem_info(DTXSID = DTXSID))
  
    if(length(chemical_info) == 0){
     chemical_info <-  as.data.table(
     expand.grid(name = c('Molar Volume',
                                         "Henry's Law",
                                         'KOC',
                                         'LogKow: Octanol-Water'
                                         ),
                                propType = c('predicted', 'experimental'),
                                value = NA_real_,
                                stringsAsFactors = FALSE)
    )
  }
  
 
  # Get chemical CAS
  chemical_cas <- chemical_details$casrn
  #strip hyphens
  chemical_cas <- gsub('-', '', chemical_cas)
  chemical_cas_ref <- 279
  chemical_cas_notes <- 'CompTox Dashboard chemical details'
  
  # Get chemical name
  chemical_name <- chemical_details$preferredName
  chemical_name_ref <- 279
  chemical_name_notes <- 'CompTox Dashboard chemical details'
  
  # Get molar volume and molecular weight for diffusivity calculations
  chemical_mw <- as.numeric(chemical_details$monoisotopicMass)
  chemical_mw_ref <- 279
  chemical_mw_notes <- 'CompTox Dashboard chemical details'
  
  #if experimental values available, use them
  chemical_molar_volume <- chemical_info[name %in% 'Molar Volume' &
                                           propType %in% 'experimental',
                                         mean(value, na.rm = TRUE)]
  chemical_molar_volume_ref <- 279
  chemical_molar_volume_notes <- 'CompTox Dashboard, chemical info endpoint, mean of experimental values'
  
  #if experimental values not available, use predicted
  if (is.nan(chemical_molar_volume)){
    chemical_molar_volume <- chemical_info[name %in% 'Molar Volume' &
                                             propType %in% 'predicted',
                                           mean(value, na.rm = TRUE)] 
    chemical_molar_volume_notes <- 'CompTox Dashboard, chemical info endpoint, mean of QSAR values'
  }
  
  chemical_air_diff_notes <- ''
  chemical_water_diff_notes <- ''
  
  if (is.nan(chemical_molar_volume)){
    chemical_molar_volume <- NA_real_
    chemical_air_diff <- NA_real_
    chemical_water_diff <- NA_real_
    chemical_molar_volume_ref <- NA_real_
    chemical_air_diff_ref <- NA_real_
    chemical_water_diff_ref <- NA_real_
  } else {
    chemical_air_diff <- air_diffusivity(M_b = chemical_mw,
                                         V_b = chemical_molar_volume)
    chemical_water_diff <- water_diffusivity(V_lebas = chemical_molar_volume)
    chemical_air_diff_ref <- 190
    chemical_water_diff_ref <- 190
    chemical_air_diff_notes <- "FSG method: https://www3.epa.gov/ceampubl/learn2model/part-two/onsite/ed-background.html"
    chemical_water_diff_notes <- "HL method: https://www3.epa.gov/ceampubl/learn2model/part-two/onsite/ed-background.html"
  }
  
  # Get Henry's Law Constant
  #start by taking mean experimental value if available
  chemical_hlc <- chemical_info[name %in% "Henry's Law" &
                                  propType %in% 'experimental',
                                mean(value, na.rm = TRUE)]
  chemical_hlc_ref <- 279
  chemical_hlc_notes <- 'CompTox Dashboard, chemical info endpoint, mean of experimental values'
  #if exp value not avail, take mean predicted value (QSAR)
  if (is.nan(chemical_hlc)){
    chemical_hlc <- chemical_info[name %in% "Henry's Law" &
                                    propType %in% 'predicted',
                                  mean(value, na.rm = TRUE)]
    chemical_hlc_notes <- 'CompTox Dashboard, chemical info endpoint, mean of QSAR values'
  }
  if (is.nan(chemical_hlc)){
    chemical_hlc <- NA_real_
    chemical_hlc_ref <- NA_real_
  }
  
  # Get logkow and koc
  chemical_logkow <- chemical_info[name %in% 'LogKow: Octanol-Water' &
                                     propType %in% 'experimental',
                                   mean(value, na.rm = TRUE)]
  chemical_logkow_ref <- 279
  chemical_logkow_notes <- 'CompTox Dashboard, chemical info endpoint, mean of experimental values'
  if (is.nan(chemical_logkow)){
    chemical_logkow <- chemical_info[name %in% 'LogKow: Octanol-Water' &
                                       propType %in% 'predicted',
                                     mean(value, na.rm = TRUE)]
    chemical_logkow_notes <- 'CompTox Dashboard, chemical info endpoint, mean of QSAR values'
  }
  if (is.nan(chemical_logkow)){
    chemical_logkow <- NA_real_
    chemical_log_kow_ref <- NA_real_
  }
  
  chemical_koc <- chemical_info[name %in% 'KOC' &
                                  propType %in% 'experimental',
                                mean(value, na.rm = TRUE)]
  chemical_koc_ref <- 279
  chemical_koc_notes <- 'CompTox Dashboard, chemical info endpoint, mean of experimental values'
  chemical_koc_notes <- ''
  if (is.nan(chemical_koc)){
    chemical_koc <- chemical_info[name %in% 'KOC' &
                                    propType %in% 'predicted',
                                  mean(value, na.rm = TRUE)]
    chemical_koc_notes <- 'CompTox Dashboard, chemical info endpoint, mean of QSAR values'
  } 
  if (is.nan(chemical_koc)){
    # Get chemical fate
    #look first in chemical details, but if it's not there, sometimes it's available through fate endpoint instead
    chemical_fate <- data.table(ctxR::get_fate_by_dtxsid(DTXSID = DTXSID))
    if (length(chemical_fate) > 0){
      chemical_koc <- chemical_fate[endpointName %in%
                                      'Soil Adsorp. Coeff. (Koc)' &
                                      valueType %in% 'experimental',
                                    mean(resultValue, na.rm = TRUE)]
      chemical_koc_ref <- 279
      chemical_koc_notes <- 'CompTox Dashboard fate, mean of experimental values'
      #if no experimental try predicted
      if (is.nan(chemical_koc)){
        chemical_koc <- chemical_fate[endpointName %in%
                                        'Soil Adsorp. Coeff. (Koc)' &
                                        valueType %in% 'predicted',
                                      mean(resultValue, na.rm = TRUE)] 
        chemical_koc_ref <- 279
        chemical_koc_notes <- 'CompTox Dashboard fate, mean of predicted values'
      }
    }
  }
  
  #if still missing, impute using SimpleTreat equation
  if (is.nan(chemical_koc)){
    if (!is.na(chemical_logkow)){
      chemical_koc <- calc_koc(DTXSID = DTXSID,
                               pH = 7,
                               Kow = 10^chemical_logkow)
      chemical_koc_ref <- 190
      chemical_koc_notes <- 'koc method: https://www.rivm.nl/publicaties/simpletreat-40-a-model-to-predict-fate-and-emission-of-chemicals-in-wastewater'
    } else {
      chemical_koc <- NA_real_
      chemical_koc_ref <- NA_real_
    }
  }
  
  # Grab bioconcentration factor for Trophic level 3, 4
  
  tr_fish_3 <- 1
  tr_fish_3_ref <- 143
  tr_fish_3_notes <- ''
  
  tr_fish_4 <- 1
  tr_fish_4_ref <- 143
  tr_fish_4_notes <- ''
  
  #look in National Bioaccumulation Factors table first
  if (chemical_cas %in% reported_BAF$CASRN){
    BAF_index <- which(reported_BAF$CASRN %in% chemical_cas)
    BAF_tr_3 <- reported_BAF[BAF_index, `National BAF TL3`]
    BAF_tr_4 <- reported_BAF[BAF_index, `National BAF TL4`]
    BAF_tr_alt <- reported_BAF[BAF_index, `Alternative BCF`]
    
    if (BAF_tr_3 != 'ND'){
      tr_fish_3 <- BAF_tr_3
      tr_fish_3_notes <- 'National bioaccumulation factors supplemental information'
      tr_fish_3_ref <- 190
    } else if (BAF_tr_alt != 'ND'){
      tr_fish_3 <- BAF_tr_alt
      tr_fish_3_notes <- 'National bioaccumulation factors supplemental information'
      tr_fish_3_ref <- 190
    }
    
    if (BAF_tr_4 != 'ND'){
      tr_fish_4 <- BAF_tr_3
      tr_fish_4_notes <- 'National bioaccumulation factors supplemental information'
      tr_fish_4_ref <- 190
    } else if (BAF_tr_alt != 'ND'){
      tr_fish_4 <- BAF_tr_alt
      tr_fish_4_notes <- 'National bioaccumulation factors supplemental information'
      tr_fish_4_ref <- 190
    }
    
  }
  
  #if not available through NBAF table, then look on Dashboard under fate endpoint
  if (tr_fish_3_ref != 190 || tr_fish_4_ref != 190){
    chemical_fate_bcf <- data.table::data.table(ctxR::get_fate_by_dtxsid(DTXSID = DTXSID))
    
    if (dim(chemical_fate_bcf)[[1]] > 0){ #if fate data were available
      
      #take geometric mean of values by model source
      bcf_fish_temp <- copy(
        chemical_fate_bcf[
          endpointName %in% 'Bioconcentration Factor' &
            !is.na(resultValue),
                                              .(
                                                Mean = exp(mean(log(resultValue)))
                                                ),
          by = .(modelSource)
          ]
        )
      #if there's an ECOTOX value use it first; if not, proceed to OPERA
      if (dim(bcf_fish_temp[modelSource %in% 'ECOTOX: aquatic',])[1] == 1){
        if (tr_fish_3_ref != 190){
          tr_fish_3 <- bcf_fish_temp[modelSource %in% 'ECOTOX: aquatic', Mean]
          tr_fish_3_ref <- 279
          tr_fish_3_notes <- 'CompTox fate endpoint, geometric mean of ECOTOX: aquatic values.'
        }
        if (tr_fish_4_ref != 190){
          tr_fish_4 <- bcf_fish_temp[modelSource %in% 'ECOTOX: aquatic', Mean]
          tr_fish_4_ref <- 279
          tr_fish_4_notes <- 'CompTox fate endpoint, geometric mean of ECOTOX: aquatic values.'
        }
      } else if (dim(bcf_fish_temp[modelSource %in% 'OPERA',])[1] == 1) {
        if (tr_fish_4_ref != 190){
          tr_fish_3 <- bcf_fish_temp[modelSource %in% 'OPERA', Mean]
          tr_fish_3_ref <- 280
          tr_fish_3_notes <- 'CompTox fate endpoint, geometric mean of OPERA values.'
        }
        if (tr_fish_4_ref != 190){
          tr_fish_4 <- bcf_fish_temp[modelSource %in% 'OPERA', Mean]
          tr_fish_4_ref <- 280
          tr_fish_4_notes <- 'CompTox fate endpoint, geometric mean of OPERA values.'
        }
      }
    }
  }
  
  # bcf_beef and bcf_milk
  #if we don't have log KoW then we can't do anything
  if (is.na(chemical_logkow)){
    bcf_beef_value <- NA_real_
    bcf_beef_ref <- NA_real_
    bcf_beef_notes <- ''
    
    bcf_milk_value <- NA_real_
    bcf_milk_ref <- NA_real_
    bcf_milk_notes <- ''
  } else { #otherwise calculate BCFs using log Kow
    bcf_beef_value <- bcf_beef(log_kow = chemical_logkow)
    bcf_beef_ref <- 49
    bcf_beef_notes <- 'RTI, 2005'
    
    bcf_milk_value <- bcf_milk(log_kow = chemical_logkow)
    bcf_milk_ref <- 49
    bcf_milk_notes <- 'RTI, 2005'
  }
  
  # BrExfruit, BrExveg, BrForage, BrGrain, BrProfruit, BrProveg, BrSilage
  #similarly, use log Kow if available
  if (is.na(chemical_logkow)){
    br_crops_value <- NA_real_
    br_crops_ref <- NA_real_
    br_crops_notes <- ''
  } else {
    br_crops_value <- bioconcentration_factor_vegetation_org(log_kow = chemical_logkow)
    br_crops_ref <- 58
    br_crops_notes <- 'Travis & Arms, 1988'
  }
  
  # Biotransfer factor vapor phase air to plant
  #similarly, use logKow and Henry's law const to calculate if available
  if (is.na(chemical_logkow) | is.na(chemical_hlc)){
    bv_value <- NA_real_
    bv_ref <- NA_real_
    bv_notes <- ''
  } else {
    bv_value <- air_to_leaf_biotransfer_factor(log_kow = chemical_logkow,
                                               HLC = chemical_hlc)
    bv_ref <- 5
    bv_notes <- 'Bacci et al., 1992'
  }
  
  #similarly use log Kow to calculate root concentration factor if available
  if (is.na(chemical_logkow)){
    root_cf_value <- NA_real_
    root_cf_ref <- NA_real_
    root_cf_notes <- ''
  } else {
    root_cf_value <- root_concentration_factor_org(log_kow = chemical_logkow)
    root_cf_ref <- 9
    root_cf_notes <- 'Briggs et al., 1982'
  }
  
  # CTPWaste
  #dilution factor from rehydrating biosolids before application
  #this equation is from the BST itself (see BST manual)
  #section C3 of Appendix C
  #LAU model requires conc to be on a wet weight basis, not a dry weight basis
  #again, we calculate it here because the BST is sometimes inconsistent in whether it properly recalculates these things
  #so we recalculate everything here
  ctpwaste_wet_value <- ctpwaste(ctp_waste_dry = biosolids_concentration)
  ctpwaste_wet_ref <- 145
  ctpwaste_wet_notes <- 'Calculated'
  organic_entry <- copy(organic_template)
  
  #These "magic numbers" are the row and column indexes of each piece of information from the Bulk Upload Tool template sheet
  organic_entry[1, 5] <- chemical_cas
  organic_entry[1, 6] <- chemical_cas_ref
  organic_entry[1, 8] <- chemical_cas_notes
  
  organic_entry[2, 5] <- substr(chemical_name, 1, 255)
  organic_entry[2, 6] <- chemical_name_ref
  organic_entry[2, 8] <- chemical_name_notes
  
  organic_entry[7, 5] <- as.character(ctpwaste_wet_value)
  organic_entry[7, 6] <- ctpwaste_wet_ref
  organic_entry[7, 8] <- ctpwaste_wet_notes
  
  organic_entry[14, 5] <- as.character(bcf_beef_value)
  organic_entry[14, 6] <- bcf_beef_ref
  organic_entry[14, 8] <- bcf_beef_notes
  
  organic_entry[15, 5] <- as.character(bcf_milk_value)
  organic_entry[15, 6] <- bcf_milk_ref
  organic_entry[15, 8] <- bcf_milk_notes
  
  organic_entry[16:22, 5] <- as.character(br_crops_value)
  organic_entry[16:22, 6] <- br_crops_ref
  organic_entry[16:22, 8] <- br_crops_notes
  
  organic_entry[23, 5] <- as.character(bv_value)
  organic_entry[23, 6] <- bv_ref
  organic_entry[23, 8] <- bv_notes
  
  organic_entry[24, 5] <- as.character(root_cf_value)
  organic_entry[24, 6] <- root_cf_ref
  organic_entry[24, 8] <- root_cf_notes
  
  organic_entry[25:26, 5] <- as.character(tr_fish_3)
  organic_entry[25:26, 6] <- tr_fish_3_ref
  organic_entry[25:26, 8] <- tr_fish_3_notes
  
  organic_entry[27:28, 5] <- as.character(tr_fish_4)
  organic_entry[27:28, 6] <- tr_fish_4_ref
  organic_entry[27:28, 8] <- tr_fish_4_notes
  
  #Hazard/tox related values:
  #These parameters are assigned a placeholder value of 1,
  #because the BST requires them to be present, 
  #even though we are not using them,
  #and they don't affect the calculated exposure values.
  
  # CSFOral
  organic_entry[30, 5] <- as.character(1)
  organic_entry[30, 6] <- 190
  organic_entry[30, 8] <- 'Default positive values to generate dose values in model output'
  
  # IUR
  organic_entry[32, 5] <- as.character(1)
  organic_entry[32, 6] <- 190
  organic_entry[32, 8] <- 'Default positive values to generate dose values in model output'
  
  # Ref_BMD_Bird
  organic_entry[38, 5] <- as.character(1)
  organic_entry[38, 6] <- 190
  organic_entry[38, 8] <- 'Default positive values to generate dose values in model output'
  
  # Ref_BMD_Mammal
  organic_entry[39, 5] <- as.character(1)
  organic_entry[39, 6] <- 190
  organic_entry[39, 8] <- 'Default positive values to generate dose values in model output'
  
  # Ref_BW_Bird
  organic_entry[40, 5] <- as.character(1)
  organic_entry[40, 6] <- 190
  organic_entry[40, 8] <- 'Default positive values to generate dose values in model output'
  
  # Ref_BW_Mammal
  organic_entry[41, 5] <- as.character(1)
  organic_entry[41, 6] <- 190
  organic_entry[41, 8] <- 'Default positive values to generate dose values in model output'
  
  # RfC
  organic_entry[42, 5] <- as.character(1)
  organic_entry[42, 6] <- 190
  organic_entry[42, 8] <- 'Default positive values to generate dose values in model output'
  
  # RfD
  organic_entry[43, 5] <- as.character(1)
  organic_entry[43, 6] <- 190
  organic_entry[43, 8] <- 'Default positive values to generate dose values in model output'
  
  # RfD_fish
  organic_entry[44, 5] <- as.character(1)
  organic_entry[44, 6] <- 190
  organic_entry[44, 8] <- 'Default positive values to generate dose values in model output'
  
  # RfD_food
  organic_entry[45, 5] <- as.character(1)
  organic_entry[45, 6] <- 190
  organic_entry[45, 8] <- 'Default positive values to generate dose values in model output'
  
  # RfD_soil
  organic_entry[46, 5] <- as.character(1)
  organic_entry[46, 6] <- 190
  organic_entry[46, 8] <- 'Default positive values to generate dose values in model output'
  
  # RfD_water
  organic_entry[47, 5] <- as.character(1)
  organic_entry[47, 6] <- 190
  organic_entry[47, 8] <- 'Default positive values to generate dose values in model output'
  
  
  #Other chemical-specific parameters
  organic_entry[50, 5] <- as.character(biosolids_concentration)
  
  organic_entry[51, 5] <- as.character(chemical_air_diff)
  organic_entry[51, 6] <- chemical_air_diff_ref
  organic_entry[51, 8] <- chemical_air_diff_notes
  
  organic_entry[52, 5] <- as.character(chemical_water_diff)
  organic_entry[52, 6] <- chemical_water_diff_ref
  organic_entry[52, 8] <- chemical_water_diff_notes
  
  organic_entry[53, 5] <- as.character(chemical_hlc)
  organic_entry[53, 6] <- chemical_hlc_ref
  organic_entry[53, 8] <- chemical_hlc_notes
  
  organic_entry[54, 5] <- as.character(chemical_koc)
  organic_entry[54, 6] <- chemical_koc_ref
  organic_entry[54, 8] <- chemical_koc_notes
  
  organic_entry[55, 5] <- as.character(chemical_logkow)
  organic_entry[55, 6] <- chemical_logkow_ref
  organic_entry[55, 8] <- chemical_logkow_notes
  
  organic_entry[56, 5] <- as.character(chemical_mw)
  organic_entry[56, 6] <- chemical_mw_ref
  organic_entry[56, 8] <- chemical_mw_notes
  
  casrn_df <- data.frame('CAS' = rep(chemical_cas, 56))
  
  organic_entry <- cbind(casrn_df, organic_entry)
  
  return(organic_entry)
}
```

# Function to generate inorganic template entry for one chemical

The following function implements the procedure for populating the inorganic template table by querying the Dashboard and/or drawing from the datasets and/or prediction functions above.

```{r}
generate_inorganic_entry <- function(DTXSID,
                                     biosolids_concentration = 1){
  #print(DTXSID)
  # Get chemical details
  chemical_details <- ctxR::get_chemical_details(DTXSID = DTXSID)
  
  if(length(chemical_details) == 0){
    chemical_details <- list(casrn = NA_character_,
                             preferredName = NA_character_,
                             monoisotopicMass = NA_real_)
  }
  
  # Get chemical info
  chemical_info <- data.table(ctxR::get_chem_info(DTXSID = DTXSID))
  
  if(length(chemical_info) == 0){
       chemical_info <- as.data.table(
         expand.grid(name = c('Molar Volume',
                                         "Henry's Law",
                                         'KOC',
                                         'LogKow: Octanol-Water'
                                         ),
                                propType = c('predicted', 'experimental'),
                                value = NA_real_,
                                stringsAsFactors = FALSE)
       )
  }
  
  # Get chemical CAS
  chemical_cas <- chemical_details$casrn
  chemical_cas <- gsub('-', '', chemical_cas)
  chemical_cas_ref <- 279
  chemical_cas_notes <- 'CompTox Dashboard chemical details'
  
  # Get chemical name
  chemical_name <- chemical_details$preferredName
  chemical_name_ref <- 279
  chemical_name_notes <- 'CompTox Dashboard chemical details'
  
  # Get molar volume and molecular weight for density calculations
  chemical_mw <- as.numeric(chemical_details$monoisotopicMass)
  chemical_mw_ref <- 279
  chemical_mw_notes <- 'CompTox Dashboard chemical details'
  
  chemical_molar_volume <- chemical_info[name %in% 'Molar Volume' &
                                           propType %in% 'experimental',
                                         mean(value, na.rm = TRUE)]
  chemical_molar_volume_ref <- 279
  chemical_molar_volume_notes <- 'CompTox Dashboard, chemical info endpoint, mean of experimental values'
  if (is.nan(chemical_molar_volume)){
    chemical_molar_volume <- chemical_info[name %in% 'Molar Volume' &
                                             propType %in% 'predicted',
                                           mean(value, na.rm = TRUE)] 
    chemical_molar_volume_notes <- 'CompTox Dashboard, chemical info endpoint, mean of QSAR values'
  }
  
  #get chemical denisty from  CCD if available,
  #otherwise from table
  density_marker <- FALSE
  chemical_density_notes <- ''
  chemical_density <- chemical_info[name %in% 'Density' &
                                      propType %in% 'experimental',
                                    mean(value, na.rm = TRUE)]
  chemical_density_ref <- 279
  chemical_density_notes <- 'CompTox Dashboard, chemical info endpoint, mean of experimental values'
  if (is.nan(chemical_density)){
    chemical_density <- chemical_info[name %in% 'Density' &
                                        propType %in% 'predicted',
                                      mean(value, na.rm = TRUE)]
    chemical_density_notes <- 'CompTox Dashboard, chemical info endpoint, mean of QSAR values'
  }
  if (is.nan(chemical_density)){
    chemical_density <- element_densities[element_densities$Name %in% chemical_name,
                                          "Density"]
    chemical_density_ref <- 190
    chemical_density_notes <- 'https://pubchem.ncbi.nlm.nih.gov/periodic-table/density/'
    
  }
  if (length(chemical_density) == 0){
    chemical_density <- NA_real_
    chemical_density_ref <- NA_real_
  }
  
  # Get lowkow and koc
  chemical_logkow <- chemical_info[name %in% 'LogKow: Octanol-Water' &
                                     propType %in% 'experimental',
                                   mean(value, na.rm = TRUE)]
  chemical_logkow_ref <- 279
  chemical_logkow_notes <- 'CompTox Dashboard, chemical info endpoint, mean of experimental values'
  if (is.nan(chemical_logkow)){
    chemical_logkow <- chemical_info[name %in% 'LogKow: Octanol-Water' &
                                       propType %in% 'predicted',
                                     mean(value, na.rm = TRUE)]
    chemical_logkow_notes <- 'CompTox Dashboard, chemical info endpoint, mean of QSAR values'
  }
  if (is.nan(chemical_logkow)){
    chemical_logkow <- NA_real_
    chemical_log_kow_ref <- ''
  }
  
  #Default value for Kd
  #very conservative value to make it all bind to biosolids
  #so we can assume most of the chemical is in the biosolids and won't just wash out
  chemical_kd <- 10000
  chemical_kd_ref <- 143
  chemical_kd_notes <- ''
  
  # Grab bioconcentration factor for Trophic level 3, 4\
  
  #as for organics:
  #look up in National BAF table if available
  #otherwise, look up in CCD Fate endpoint
  
  tr_fish_3 <- 1
  tr_fish_3_ref <- 143
  tr_fish_3_notes <- ''
  tr_fish_4 <- 1
  tr_fish_4_ref <- 143
  tr_fish_4_notes <- ''
  
  if (chemical_cas %in% reported_BAF$CASRN){
    BAF_index <- which(reported_BAF$CASRN %in% chemical_cas)
    BAF_tr_3 <- reported_BAF[BAF_index, `National BAF TL3`]
    BAF_tr_4 <- reported_BAF[BAF_index, `National BAF TL4`]
    BAF_tr_alt <- reported_BAF[BAF_index, `Alternative BCF`]
    
    if (BAF_tr_3 != 'ND'){
      tr_fish_3 <- BAF_tr_3
      tr_fish_3_notes <- 'National bioaccumulation factors supplemental information'
      tr_fish_3_ref <- 190
    } else if (BAF_tr_alt != 'ND'){
      tr_fish_3 <- BAF_tr_alt
      tr_fish_3_notes <- 'National bioaccumulation factors supplemental information'
      tr_fish_3_ref <- 190
    }
    
    if (BAF_tr_4 != 'ND'){
      tr_fish_4 <- BAF_tr_3
      tr_fish_4_notes <- 'National bioaccumulation factors supplemental information'
      tr_fish_4_ref <- 190
    } else if (BAF_tr_alt != 'ND'){
      tr_fish_4 <- BAF_tr_alt
      tr_fish_4_notes <- 'National bioaccumulation factors supplemental information'
      tr_fish_4_ref <- 190
    }
    
  }
  
  if (tr_fish_3_ref != 190 || tr_fish_4_ref != 190){
    # print('Pinging API for fate...')
    chemical_fate_bcf <- data.table::data.table(
      ctxR::get_fate_by_dtxsid(DTXSID = DTXSID)
    )
    
    if (dim(chemical_fate_bcf)[[1]] > 0){
      
      bcf_fish_temp <- copy(
        chemical_fate_bcf[
          endpointName %in% 'Bioconcentration Factor' &
            !is.na(resultValue),
          .(
            Mean = exp(mean(log(resultValue)))
          ),
          by = .(modelSource)
        ]
      )
      if (dim(bcf_fish_temp[modelSource %in% 'ECOTOX: aquatic',])[1] == 1){
        if (tr_fish_3_ref != 190){
          tr_fish_3 <- bcf_fish_temp[modelSource %in% 'ECOTOX: aquatic', Mean]
          tr_fish_3_ref <- 279
          tr_fish_3_notes <- 'CompTox Dashboard, fate endpoint, geometric mean of ECOTOX: aquatic values.'
        }
        if (tr_fish_4_ref != 190){
          tr_fish_4 <- bcf_fish_temp[modelSource %in% 'ECOTOX: aquatic', Mean]
          tr_fish_4_ref <- 279
          tr_fish_4_notes <- 'CompTox Dashboard, fate endpoint, geometric mean of ECOTOX: aquatic values.'
        }
      } else if (dim(bcf_fish_temp[modelSource %in% 'OPERA',])[1] == 1) {
        if (tr_fish_3_ref != 190){
          tr_fish_3 <- bcf_fish_temp[modelSource %in% 'OPERA', Mean]
          tr_fish_3_ref <- 280
          tr_fish_3_notes <- 'CompTox Dashboard, fate endpoint, geometric mean of OPERA values.'
        }
        if (tr_fish_4_ref != 190){
          tr_fish_4 <- bcf_fish_temp[modelSource %in% 'OPERA', Mean]
          tr_fish_4_ref <- 280
          tr_fish_4_notes <- 'CompTox Dashboard, fate endpoint, geometric mean of OPERA values.'
        }
      }
    }
  }
  
  
  # diffusivity in water
  if (is.na(chemical_density)){
    diff_water_value <- NA_real_
    diff_water_ref <- NA_real_
    diff_water_notes <- ''
  } else {
    diff_water_value <- diff_water_inor(mol_weight = chemical_mw,
                                        density = chemical_density)
    diff_water_ref <- 145
    diff_water_notes <- 'Calculated'
  }
  
  # CTPWaste
  ctpwaste_wet_value <- ctpwaste(ctp_waste_dry = biosolids_concentration)
  ctpwaste_wet_ref <- 145
  ctpwaste_wet_notes <- 'Calculated' 
  
  #Generating table
  inorganic_entry <- copy(inorganic_template)
  
  #Filling out table
  #These "magic numbers" are the row and column indexes of each piece of information from the Bulk Upload Tool template sheet
  inorganic_entry[1, 5] <- chemical_cas
  inorganic_entry[1, 6] <- chemical_cas_ref
  inorganic_entry[1, 8] <- chemical_cas_notes
  
  inorganic_entry[2, 5] <- substr(chemical_name, 1, 255)
  inorganic_entry[2, 6] <- chemical_name_ref
  inorganic_entry[2, 8] <- chemical_name_notes
  
  inorganic_entry[5, 5] <- as.character(ctpwaste_wet_value)
  inorganic_entry[5, 6] <- ctpwaste_wet_ref
  inorganic_entry[5, 8] <- ctpwaste_wet_notes
  
  inorganic_entry[11, 5] <- as.character(diff_water_value)
  inorganic_entry[11, 6] <- diff_water_ref
  inorganic_entry[11, 8] <- diff_water_notes
  
  inorganic_entry[14:15, 5] <- as.character(tr_fish_3)
  inorganic_entry[14:15, 6] <- tr_fish_3_ref
  inorganic_entry[14:15, 8] <- tr_fish_3_notes
  
  inorganic_entry[16:17, 5] <- as.character(tr_fish_4)
  inorganic_entry[16:17, 6] <- tr_fish_4_ref
  inorganic_entry[16:17, 8] <- tr_fish_4_notes
  
  # CSFOral
  inorganic_entry[27, 5] <- as.character(1)
  inorganic_entry[27, 6] <- 190
  inorganic_entry[27, 8] <- 'Default positive values to generate dose values in model output'
  
  # IUR
  inorganic_entry[28, 5] <- as.character(1)
  inorganic_entry[28, 6] <- 190
  inorganic_entry[28, 8] <- 'Default positive values to generate dose values in model output'
  
  # Ref_BMD_Bird
  inorganic_entry[29, 5] <- as.character(1)
  inorganic_entry[29, 6] <- 190
  inorganic_entry[29, 8] <- 'Default positive values to generate dose values in model output'
  
  # Ref_BMD_Mammal
  inorganic_entry[30, 5] <- as.character(1)
  inorganic_entry[30, 6] <- 190
  inorganic_entry[30, 8] <- 'Default positive values to generate dose values in model output'
  
  # Ref_BW_Bird
  inorganic_entry[31, 5] <- as.character(1)
  inorganic_entry[31, 6] <- 190
  inorganic_entry[31, 8] <- 'Default positive values to generate dose values in model output'
  
  # Ref_BW_Mammal
  inorganic_entry[32, 5] <- as.character(1)
  inorganic_entry[32, 6] <- 190
  inorganic_entry[32, 8] <- 'Default positive values to generate dose values in model output'
  
  # RfC
  inorganic_entry[33, 5] <- as.character(1)
  inorganic_entry[33, 6] <- 190
  inorganic_entry[33, 8] <- 'Default positive values to generate dose values in model output'
  
  # RfD
  inorganic_entry[34, 5] <- as.character(1)
  inorganic_entry[34, 6] <- 190
  inorganic_entry[34, 8] <- 'Default positive values to generate dose values in model output'
  
  # RfD_fish
  inorganic_entry[35, 5] <- as.character(1)
  inorganic_entry[35, 6] <- 190
  inorganic_entry[35, 8] <- 'Default positive values to generate dose values in model output'
  
  # RfD_food
  inorganic_entry[36, 5] <- as.character(1)
  inorganic_entry[36, 6] <- 190
  inorganic_entry[36, 8] <- 'Default positive values to generate dose values in model output'
  
  # RfD_soil
  inorganic_entry[37, 5] <- as.character(1)
  inorganic_entry[37, 6] <- 190
  inorganic_entry[37, 8] <- 'Default positive values to generate dose values in model output'
  
  # RfD_water
  inorganic_entry[38, 5] <- as.character(1)
  inorganic_entry[38, 6] <- 190
  inorganic_entry[38, 8] <- 'Default positive values to generate dose values in model output'
  
  inorganic_entry[41, 5] <- as.character(biosolids_concentration)
  
  inorganic_entry[42, 5] <- as.character(chemical_density)
  inorganic_entry[42, 6] <- chemical_density_ref
  inorganic_entry[42, 8] <- chemical_density_notes
  
  inorganic_entry[43, 5] <- as.character(chemical_kd)
  inorganic_entry[43, 6] <- chemical_kd_ref
  inorganic_entry[43, 8] <- chemical_kd_notes
  
  inorganic_entry[44, 5] <- as.character(chemical_mw)
  inorganic_entry[44, 6] <- chemical_mw_ref
  inorganic_entry[44, 8] <- chemical_mw_notes
  
  casrn_df <- data.frame('CAS' = rep(chemical_cas, 44))
  
  inorganic_entry <- cbind(casrn_df, inorganic_entry)
  
  return(inorganic_entry)
}
```

## Generalized function to generate the appropriate entry

```{r}
generate_entry <- function(DTXSID,
                         chemical_class,
                         biosolids_concentration = 1){
  if(chemical_class %in% "Organic"){
    out <- generate_organic_entry(DTXSID = DTXSID,
                         biosolids_concentration = biosolids_concentration)
  }else{
    out <- generate_inorganic_entry(DTXSID = DTXSID,
                         biosolids_concentration = biosolids_concentration)
  }
  return(out)
}
```


# Function to generate ecological template entry for one chemical

The following function implements the procedure for populating the ecological template table. These parameters are all currently given a placeholder value of 1.

```{r}
generate_eco_entry <- function(DTXSID,
                               chemical_cas = NULL,
                               truncate = FALSE){
  
  if(is.null(chemical_cas)){
    #look up CASRN if necessary
  # Get chemical details
  chemical_details <- data.table(ctxR::get_chemical_details(DTXSID = DTXSID))
  
  if (length(chemical_details) == 0){
    return(NA)
  }
  
  chemical_cas <- gsub('-', '', x = chemical_details$casrn)
  }
  
  #substitute a placeholder value of 1 here
  chemical_bcf <- 1
  chemical_bcf_ref <- 190
  chemical_bcf_notes <- 'Default value given by BST default value'
 
  #print('Generating table')
  eco_entry <- copy(eco_template)
  
  #print('Filling out table')
  eco_entry[8, 5] <- chemical_bcf
  eco_entry[8, 6] <- chemical_bcf_ref
  eco_entry[8, 7] <- chemical_bcf_notes
  
  eco_entry[9, 5] <- chemical_bcf
  eco_entry[9, 6] <- chemical_bcf_ref
  eco_entry[9, 7] <- chemical_bcf_notes
  
  if (!truncate){
    eco_entry[10:54, 5] <- 1
    eco_entry[10:54, 6] <- 190
    eco_entry[10:54, 7] <- 'User set to "dummy" value of 1 to run model'
  }
  
  # Remove rows that can be deleted if missing data
  deletion_rows <- c(40:54)[which(which(sapply(eco_entry[, c('Value')], is.na)) >= 40)]
  if (length(deletion_rows)) {
    eco_entry <- eco_entry[-deletion_rows,]
  }
  
  casrn_df <- data.frame('CAS' = rep(chemical_cas, dim(eco_entry)[[1]]))
  
  eco_entry <- cbind(casrn_df, eco_entry)
  
  return(eco_entry)
  
}
```

# Data checks

The following functions implement data checks.

## Check whether each chemical is on the list of pre-loaded BST chemicals

```{r}
check_bst_preloads <- function(entry){
  
  cas <- entry$CAS
is_preload <- ifelse(CASRN_stripped %in% previously_loaded_casrns$CASRN,
                           "Preloaded",
                           "Not preloaded")
return(is_preload)
}
```

## Check organic, inorganic, or other

```{r}
check_class <- function(DTXSIDs,
                        inchikeys = NULL){
  
  #if inchikeys not already provided, query from CCD
  if(is.null(inchikeys)){
    inchikeys <- ctxR::get_chemical_details_batch(DTXSID = DTXSIDs)[, inchikey]
  }
  
   #get previously-saved ClassyFire results for these inchikeys
  this_dt <- data.table(inchikey = inchikeys)
  tmp <- chem_classes[this_dt, on = "inchikey"]

  tmp[, 
             chemical_class := dplyr::case_when(kingdom %in% "Organic compounds" ~ "Organic",
                                               kingdom %in% "Inorganic compounds" ~ "Inorganic",
                                               .default = "Non-classifiable")]
  
  #return chemical class
  return(tmp$chemical_class)
}
```


## Check whether each chemical is a DLC

```{r}
check_dlc <- function(DTXSIDs,
                      inchikeys = NULL){
  
  #if inchikeys not already provided, query from CCD
  if(is.null(inchikeys)){
    inchikeys <- ctxR::get_chemical_details_batch(DTXSID = DTXSIDs)[, inchikey]
  }
  
  #get previously-saved ClassyFire results for these inchikeys
  this_dt <- data.table(inchikey = inchikeys)
  tmp <- chem_classes[this_dt, on = "inchikey"]
  
  is_dlc <- tmp[, dplyr::case_when(class %in% 'Benzodioxins' ~ 'is dioxin',
                                         class %in% 'Benzofurans' ~ 'is furan',
                              level6 %in% 'Polychlorinated biphenyls'~ 'is PCB',
                              !(kingdom %in% c("Organic compounds",
                                               "Inorganic compounds")) ~ "NA",
                            .default = "not DLC")]
  
  return(is_dlc)
}
```


## Check for missing information

Check missing values for an organic template table entry:

```{r}
#give it a sheet and it will tell you what is and is not missing
organic_entry_missing_information <- function(organic_entry){
 
  if(!is.data.table(organic_entry)){
    organic_entry <- as.data.table(organic_entry)
  }
  
  values <- organic_entry[c(1:2, 7, 14:28, 30, 32, 38:47, 50:56),
                          Value]
  missing_values <- which(sapply(values, is.na))
  if (length(missing_values) > 0){
    missing_data <- paste(organic_entry[c(1:2, 7, 14:28, 30, 32, 38:47, 50:56)[missing_values],
                                          ModelCode],
                            collapse = ', ')
  }else{
    missing_data <- "no missing"
  }
  return(missing_data)
}
```

Check missing values for an inorganic template table entry:

```{r}
inorganic_entry_missing_information <- function(inorganic_entry){
  
  
  if(!is.data.table(inorganic_entry)){
    inorganic_entry <- as.data.table(inorganic_entry)
  }
  values <- inorganic_entry[c(1:2, 5, 11, 14:17, 27:38, 41:44), Value]
  missing_values <- which(sapply(values, is.na))
  if (length(missing_values) > 0){
    missing_data <- paste(inorganic_entry[c(1:2, 5, 11, 14:17, 27:38, 41:44)[missing_values], ModelCode], collapse = ', ')
  }else{
    missing_data <- "no missing"
  }
  
  return(missing_data)
}
```

### General function to check missing values

```{r}
check_missing_info <- function(entry,
                               chemical_class){
  if(chemical_class %in% "Organic"){
    missing_data <- organic_entry_missing_information(organic_entry = entry)
  }else if(chemical_class %in% "Inorganic"){
    missing_data <- inorganic_entry_missing_information(inorganic_entry = entry)
  }
  
  return(missing_data)
}
```

## Check that values are within range

We also need to check that values are within the acceptable ranges (as defined by the BST).

Check the ranges for an organic entry:

```{r}
organic_entry_range_check <- function(organic_entry){
  
  organic_entry <- copy(organic_entry)
  
  if(!is.data.table(organic_entry)){
    organic_entry <- as.data.table(organic_entry)
  }
  
  values <- as.numeric(organic_entry[c(7, 14:28, 30, 32, 38:47, 50:56), Value])
  min_values <- as.numeric(organic_entry[c(7, 14:28, 30, 32, 38:47, 50:56), `Minimum Value`])
  max_values <- as.numeric(organic_entry[c(7, 14:28, 30, 32, 38:47, 50:56), `Maximum Value`])
  
  # No upper bound for CTPwaste
  max_values[[1]] <- Inf
  
  high_string <- 'no high'
  low_string <- 'no low'
  
  max_indices <- which(values > max_values)
  if (length(max_indices) > 0){
    #If it is more than 1% out of range, mark it as out of range
    #If it is less than 1% out of range, substitute the bound
    out_of_range_high <- which(((values[max_indices] - max_values[max_indices])/(max_values[max_indices])) > 0.01)
    if (length(out_of_range_high) > 0){
      # print(values[max_indices[out_of_range_high]])
      # print(max_values[max_indices[out_of_range_high]])
      high_string <- paste(organic_entry[c(7, 14:28, 30, 32, 38:47, 50:56)[max_indices[out_of_range_high]],
                                         ModelCode],
                           collapse = ', ')
    } else {
      organic_entry[c(7, 14:28, 30, 32, 38:47, 50:56)[max_indices],
                    Value := as.character(max_values[max_indices])]
    }
  }
  
  min_indices <- which(values < min_values)
  if (length(min_indices > 0)){
       #If it is more than 1% out of range, mark it as out of range
    #If it is less than 1% out of range, substitute the bound
    #but only if the lower bound is not zero
    zero_indices <- intersect(c(1:28, 30:33), min_indices)
    non_zero_indices <- intersect(c(29, 34:35), min_indices)
    #to do: use the range itself to reflect max allowable interval rather than value itself
    out_of_range_low <- which(((min_values[non_zero_indices] - values[non_zero_indices])/(min_values[non_zero_indices])) > 0.01)
    out_of_range_low <- sort(union(non_zero_indices[out_of_range_low], zero_indices))
    if (length(out_of_range_low) > 0){
      low_string <- paste(organic_entry[c(7, 14:28, 30, 32, 38:47, 50:56)[out_of_range_low],
                                        ModelCode],
                          collapse = ', ')
    } else {
      organic_entry[c(7, 14:28, 30, 32, 38:47,50:56)[min_indices],
                    Value := as.character(min_values[min_indices])]
    }
  }
  
 organic_entry[, c("out_of_range_high",
                      "out_of_range_low") := .(high_string,
                                               low_string)]

  return(organic_entry)
  
}
```

Check the ranges for an inorganic entry:

```{r}
inorganic_entry_range_check <- function(inorganic_entry){
 
  inorganic_entry <- copy(inorganic_entry)
  
   if(!is.data.table(inorganic_entry)){
    inorganic_entry <- as.data.table(inorganic_entry)
   }
  
  values <- as.numeric(inorganic_entry[c(5, 11, 14:17, 27:38, 41:44), Value])
  min_values <- as.numeric(inorganic_entry[c(5, 11, 14:17, 27:38, 41:44), `Minimum Value`])
  max_values <- as.numeric(inorganic_entry[c(5, 11, 14:17, 27:38, 41:44), `Maximum Value`])
  
  # No upper bound for CTPwaste
  max_values[[1]] <- Inf
  
  high_string <- 'no high'
  low_string <- 'no low'
  
  max_indices <- which(values > max_values)
  if (length(max_indices) > 0){
        #If it is more than 1% out of range, mark it as out of range
    #If it is less than 1% out of range, substitute the bound
    out_of_range_high <- which(((values[max_indices] - max_values[max_indices])/(max_values[max_indices])) > 0.01)
    if (length(out_of_range_high) > 0){
      high_string <- paste(inorganic_entry[c(5, 11, 14:17, 27:38, 41:44)[max_indices[out_of_range_high]], ModelCode], collapse = ', ')
    } else {
      inorganic_entry[c(5, 11, 14:17, 27:38, 41:44)[max_indices], Value := as.character(max_values[max_indices])]
    }
  }
  
  min_indices <- which(values < min_values)
  if (length(min_indices > 0)){
    zero_indices <- intersect(c(1:18, 20:21), min_indices)
    non_zero_indices <- intersect(c(19, 22), min_indices)
        #If it is more than 1% out of range, mark it as out of range
    #If it is less than 1% out of range, substitute the bound
    out_of_range_low <- which(((min_values[non_zero_indices] - values[non_zero_indices])/(min_values[non_zero_indices])) > 0.01)
    out_of_range_low <- sort(union(non_zero_indices[out_of_range_low], zero_indices))
    if (length(out_of_range_low) > 0){
      low_string <- paste(inorganic_entry[c(5, 11, 14:17, 27:38, 41:44)[out_of_range_low], ModelCode], collapse = ', ')
    } else {
      inorganic_entry[c(5, 11, 14:17, 27:38, 41:44)[min_indices], Value := as.character(min_values[min_indices])]
    }
  }
  
  inorganic_entry[, c("out_of_range_high",
                      "out_of_range_low") := .(high_string,
                                               low_string)]

  return(inorganic_entry)
  
}
```

### General function to check range

```{r}
check_range <- function(entry,
                        chemical_class){
  if(chemical_class %in% "Organic"){
    out <- organic_entry_range_check(organic_entry = entry)
  }else if(chemical_class %in% "Inorganic"){
    out <- inorganic_entry_range_check(inorganic_entry = entry)
  }
  
  return(out)
  
}
```

# Read in Bulk Uploader Tool sheets

```{r}
bulkuploader_Notes <- readxl::read_excel(path = 'data/BST_BulkChemicalImportTool.xlsm',
                                         sheet = 'Notes')

bulkuploader_Template_Chem_Org <- readxl::read_excel(path = 'data/BST_BulkChemicalImportTool.xlsm',
                                         sheet = 'Template_Chem_Org')
bulkuploader_Template_Chem_Inorg <- readxl::read_excel(path = 'data/BST_BulkChemicalImportTool.xlsm',
                                         sheet = 'Template_Chem_Inorg')
bulkuploader_Template_Eco <- readxl::read_excel(path = 'data/BST_BulkChemicalImportTool.xlsm',
                                         sheet = 'Template_Eco')
bulkuploader_References <- readxl::read_excel(path = 'data/BST_BulkChemicalImportTool.xlsm',
                                         sheet = 'References')
```

First, prepare a table of DTXSIDs and concentrations from the concentration data. The following function does this.

```{r}
make_conc_tbl <- function(which_conc = "mean" #or "95th"
                          ){
  #Take the most recent available option
  if(which_conc %in% "mean"){
    vars <- c("2009 Mean with Nondetects Set to 1/2 MRL",
              "2001 Mean with Nondetects Set to 1/2 MRL",
              "1988 Mean with Nondetects Set to MRL")
  }else if(which_conc %in% "95th"){
    #either the non-parameteric or the lognormal one will be NA
    NSSS_tmp <- copy(NSSS)
    NSSS_tmp[, ]
    
    vars <- c("2009 95th Percentile Concentration Estimates with Nondetects Set to 1/2 MRL (Non-Parametric Model)",
              "2009 95th Percentile Concentration Estimates with Nondetects Set to 1/2 MRL (Lognormal Model)",
              "2001 95th Percentile Concentration Estimate with Nondetects Set to 1/2 MRL",
              "1988 95th Percentile Concentration Estimate with Nondetects Set to MRL")
  }
  
  units_vars <- c("2009 Unit",
                  "2001 Unit",
                  "1988 Unit")
  
  DT_long <- melt(NSSS,
                  id.vars = "DTXSID",
                  measure.vars = vars,
                  na.rm = TRUE,
                  variable.factor = FALSE)
  
  DT_long[, year := substr(variable,
                         1,
                         4)]
  
    #the first row within each DTXSID will be the most recent
  DT_out <- DT_long[, .SD[1], by = DTXSID]
  setnames(DT_out,
         c("variable", "value"),
         c("conc_variable", "conc"))
  
  DT_long_unit <- melt(NSSS,
                  id.vars = "DTXSID",
                  measure.vars = units_vars,
                  na.rm = TRUE,
                  variable.factor = FALSE)
  
    DT_long_unit[, year := substr(variable,
                         1,
                         4)]
    
    setorder(DT_long_unit,
             DTXSID,
             -year)
    
    
  setnames(DT_long_unit,
         "value",
         "unit")
  
  DT_long_unit[, variable := NULL]
  
   #the first row within each DTXSID will be the most recent
  DT_out_unit <- DT_long_unit[, .SD[1], by = DTXSID]
  
    DT_out <- DT_out_unit[DT_out, on = c("DTXSID", "year")]
  
  #convert all to mg/kg units
  DT_out[unit %in% "MG/KG", conc_mgkg := conc]
  DT_out[unit %in% "UG/KG", conc_mgkg := conc * 1e-3]
  DT_out[unit %in% "NG/KG", conc_mgkg := conc * 1e-6]
  
  DT_out[, year := NULL]
  
  return(DT_out)
  
}
```


# Main function to generate table entries

This function first identifies DTXSIDs that are excluded due to lack of concentration data, being previously loaded into the BST, being neither organic nor inorganic, or being DLCs. For the non-excluded DLCs, it creates the appropriate chemical-specific table (organic or inorganic), and the ecological table. Then it checks the chemical-specific tables for missing information and out-of-range information. If there is missing information, or if out-of-range information can't be reasonably corrected, then it marks those DTXSIDs so that the corresponding table entries are excluded. Finally, it concatenates the non-excluded tables. It returns the tables in appropriate format to be saved in a workbook for the Bulk Upload Tool. Alongside them, it returns a list of excluded chemicals, with information on their reasons for exclusion.

```{r}
# This function takes in a vector of DTXSIDs.
generate_tbls <- function(DTXSIDs,
                          which_conc = "mean"){
  
  if(which_conc %in% c("mean", "95th")){
  #create concentration data table
  conc_tbl <- make_conc_tbl(which_conc = which_conc)
  }else if(which_conc %in% '1ppb'){
    conc_tbl <- data.table(DTXSID = DTXSIDs,
                           conc_mgkg = 1e-3)
  }else{
    stop("which_conc must be either 'mean', '95th', or '1ppb'")
  }
  
  #initialize output table, starting with the list of DTXSIDs
  out_tbl <- data.table(DTXSID = DTXSIDs)
  #merge in concentration data where available
  #fill with NA where concentration data not available
  
  out_tbl <- conc_tbl[out_tbl,
                     on = "DTXSID",
                     nomatch = NA]
  
  #Does chemical have concentration data?
  out_tbl[, has_conc_data := ifelse(is.na(conc_mgkg),
                                    "no conc data",
                                    "has conc data")]
  
  #Is chemical already preloaded into BST?
  #Check using the CASRN.
  #Query CCD to get CASRN.
   chemical_details <- ctxR::get_chemical_details_batch(DTXSID = DTXSIDs)
   out_tbl <- chem_details[, .(dtxsid,
                                 casrn,
                                 preferredName,
                              inchikey)][out_tbl,
              on = c("dtxsid" =  "DTXSID")]
   setnames(out_tbl,
            "dtxsid",
            "DTXSID")
   
   out_tbl[, casrn_stripped := gsub(x = casrn,
                                    pattern = "-",
                                    replacement = "")]
   
   #compare to table of previously loaded CASRNs
  out_tbl[has_conc_data %in% "has conc data",
          is_BST_preload := ifelse(casrn_stripped %in% 
                                       as.character(previously_loaded_casrns$CASRN),
                                     "preloaded",
                                     "not preloaded")]
  
  out_tbl[has_conc_data %in% "no conc data",
          is_BST_preload := "exclude"]
  
  #Chemical class checks:
  
    
  #Is chemical organic, inorganic, or other?
  out_tbl[is_BST_preload %in% "not preloaded",
          chemical_class := check_class(DTXSID,
                                          inchikeys = inchikey)]
  out_tbl[!(is_BST_preload %in% "not preloaded"),
          chemical_class := "exclude"]
  
  #Is chemical a DLC?
  out_tbl[chemical_class %in% c("Organic",
                                "Inorganic"), is_DLC := check_dlc(DTXSID,
                                inchikeys = inchikey)]
  
   out_tbl[!(chemical_class %in% c("Organic",
                                "Inorganic")),
           is_DLC := "exclude"]

  #determine which DTXSIDs to be excluded a priori
  #don't even bother trying to create table entries for these
  out_tbl[, exclude_prior := (!(has_conc_data %in% "has conc data")) |
            (!(is_BST_preload %in% "not preloaded")) |
           ( !(chemical_class %in% c("Organic",
                                "Inorganic"))) |
            (!(is_DLC %in% "not DLC"))
            ]
  
  #For chemicals which pass all of the above checks:
  
  #Create appropriate entry (organic or inorganic)
  chem_entries <- out_tbl[exclude_prior %in% FALSE, 
    generate_entry(DTXSID = DTXSID,
                         chemical_class = chemical_class,
                   biosolids_concentration = conc_mgkg),
    by = .(DTXSID,
           chemical_class)]
  
    #and create ecological entry
  eco_entries <- out_tbl[exclude_prior %in% FALSE, 
    generate_eco_entry(DTXSID = DTXSID,
                   chemical_cas = casrn_stripped),
    by = DTXSID]
  
  #Check for missing information
  #This table will have one row for each DTXSID
  #with a variable "missing_data" that will be empty if nothing was missing,
  #and contain the names of the missing parameters if any are missing
  chem_entries_missing <- chem_entries[, .(missing_data = check_missing_info(entry = .SD,
                                    chemical_class = unique(chemical_class))),
               by = .(DTXSID,
                      chemical_class)]

  #Mark which entries have missing values (if any)
  chem_entries2 <- chem_entries[chem_entries_missing,
                               on = c("DTXSID",
                                      "chemical_class")]
  
  #For those without missing values:
    #Check for values within operating range,
  #correct if possible,
  #and mark if not possible
  chem_entries_rangecheck <- chem_entries2[missing_data %in% "no missing",
                                         check_range(entry = .SD,
                                    chemical_class = unique(chemical_class)),
               by = .(DTXSID,
                      chemical_class,
                      missing_data)]
  
    #Remove any entries marked for removal due to missing information,
  #or out-of-range values that could not be corrected
  chem_entries_final <- chem_entries_rangecheck[(missing_data %in% "no missing") &
                                             (out_of_range_high %in% "no high") &
                                             (out_of_range_low %in% "no low"),
                                           .SD,
                                         .SDcols = setdiff(names(chem_entries_rangecheck),
                                                           c("chemical_class",
                                                             "missing_data",
                                                             "out_of_range_high",
                                                             "out_of_range_low"))]
  
  #table of check results: one row for each DTXSID
  chem_entries_check <- unique(chem_entries_rangecheck[, .(DTXSID,
                                                    out_of_range_high,
                                                    out_of_range_low)])
  #merge in missing data information
  chem_entries_check <- chem_entries_check[chem_entries_missing[, .(DTXSID, missing_data)],
                                           on = "DTXSID",
                                           nomatch = NA]
   #ensure columns are in the correct order
  chem_entries_check <- chem_entries_check[, .SD, .SDcols = c("DTXSID",
                                                              "missing_data",
                                                              "out_of_range_high",
                                                              "out_of_range_low")]
  chem_entries_check[!(missing_data %in% "no missing"),
                     c("out_of_range_high",
                       "out_of_range_low") := "exclude"]
  
  chem_entries_check[!(out_of_range_high %in% "no high"),
                     out_of_range_low := "exclude"]
  
  #merge checks with exclusions
  out_tbl2 <- chem_entries_check[out_tbl,
                     on = "DTXSID",
                     nomatch = NA]
  #ensure columns are in the correct order
  out_tbl2 <- out_tbl2[, .SD, .SDcols = union(names(out_tbl),
                                            names(chem_entries_check))]
  
     #exclude known bad-format chemicals 
  out_tbl2[casrn_stripped %in% bad_chems$casrn_stripped,
           bad_format := "bad format"]
  out_tbl2[!(casrn_stripped %in% bad_chems$casrn_stripped),
           bad_format := "format OK"]

  
  #if chemical was excluded a priori, set all the post-CCD-query data checks to 'exclude'
  out_tbl2[exclude_prior %in% TRUE,
           c("missing_data",
           "out_of_range_high",
                       "out_of_range_low",
           "bad_format") := "exclude"]
  
    #for anything excluded due to post-CCD data checks, set bad_format to "exclude"
    out_tbl2[out_of_range_low %in% "exclude",
          bad_format := "exclude"]
  
  #for anything already preloaded, change the downstream filtering from "exclude" to "preloaded"
  out_tbl2[is_BST_preload %in% "preloaded",
           c("chemical_class",
             "is_DLC",
             "missing_data",
             "out_of_range_high",
             "out_of_range_low",
             "bad_format") := "preloaded"]

  #Add a "Bulk Upload" data filtering stage
  #immediately after the bad_format filtering stage
  out_tbl2[!(bad_format %in% "format OK"),
           Bulk_Upload := "exclude"]
  out_tbl2[bad_format %in% "format OK",
           Bulk_Upload := "Bulk Upload"]
  out_tbl2[bad_format %in% "preloaded",
           Bulk_Upload := "preloaded"]
  
  out_tbl2[, is_run := dplyr::case_when(Bulk_Upload %in% c("Bulk Upload",
                                                             "preloaded") ~ "run",
                                     Bulk_Upload %in% "exclude" ~ "not run",
                                     .default = "not run")]
  
  #set ordering for data filtering columns
  #this will be used for Sankey diagram later
  out_tbl2[, has_conc_data := factor(has_conc_data,
                                     levels =  c("has conc data",
                                                 "no conc data"))]
  out_tbl2[, is_BST_preload := factor(is_BST_preload,
                                      levels = c("preloaded",
                                                 "not preloaded",
                                                 "exclude"))]
  out_tbl2[, chemical_class := factor(chemical_class,
                                      levels = c("preloaded",
                                      "Inorganic",
                                                 "Organic",
                                                 "Non-classifiable",
                                                 "exclude"))]
  out_tbl2[, is_DLC := factor(is_DLC,
                              levels = c("preloaded",
                              "not DLC",
                                         "is furan",
                                         "is dioxin",
                                         "is PCB",
                                         "exclude"))]
  
  out_tbl2[, missing_data := factor(missing_data,
                                      levels = c("preloaded",
                                      "no missing",
                                                 setdiff(unique(missing_data),
                                                         c("no missing",
                                                           "preloaded",
                                                           "exclude")),
                                                 "exclude"))]
  out_tbl2[, out_of_range_high := factor(out_of_range_high,
                                      levels = c("preloaded",
                                      "no high",
                                                 setdiff(unique(out_of_range_high),
                                                         c("no high",
                                                           "preloaded",
                                                           "exclude")),
                                                 "exclude"))]
    out_tbl2[, out_of_range_low := factor(out_of_range_low,
                                      levels = c("preloaded",
                                      "no low",
                                                 setdiff(unique(out_of_range_low),
                                                         c("no low",
                                                           "preloaded",
                                                           "exclude")),
                                                 "exclude"))]
    
    out_tbl2[, bad_format := factor(bad_format,
                                    levels = c("preloaded",
                                               "format OK",
                                               "bad format",
                                               "exclude"))]
    
    
    out_tbl2[, Bulk_Upload := factor(Bulk_Upload,
                                     levels = c("preloaded",
                                     "Bulk Upload",
                                                "exclude"))]
    
    out_tbl2[, is_run := factor(is_run,
                                levels = c("run",
                                           "not run"))]
  
  
  
  #now create list of chemicals
    #run a final check of DTXSIDs to include
    include_dtxsid <- out_tbl2[Bulk_Upload %in% "Bulk Upload", DTXSID]
  chem_entries_final <- chem_entries_final[DTXSID %in% include_dtxsid, ]
    
  #create table of included chemicals
  new_df <- dcast(
    unique(
      chem_entries_final[ModelCode %in% c("CAS",
                                                      "ChemName",
                                                      "ChemType"),
                         .(DTXSID,
                           ModelCode,
                           Value)]
      ),
                  DTXSID ~ ModelCode,
                  value.var = "Value")
  new_df[, UserAdded := TRUE]
  #sort in the same order as for chem_entries_final and eco_entries
  DTXSID_levels <- unique(chem_entries_final$DTXSID)
  new_df[, DTXSID := factor(DTXSID, levels = DTXSID_levels)]
  setorder(new_df, DTXSID)

  #ensure eco entries is filtered properly
  eco_entries <- eco_entries[DTXSID %in% new_df$DTXSID]
  #and sorted properly
  eco_entries[, DTXSID := factor(DTXSID, levels = DTXSID_levels)]
  setorder(eco_entries, DTXSID)
  
    #now remove superfluous DTXSID columns
  new_df[, DTXSID := NULL]
  chem_entries_final[, DTXSID := NULL]
  eco_entries[, DTXSID := NULL]
  
  setnames(new_df,
           "ChemName",
           "Constituent")
  
  return(list('tbl_Chemicals' = new_df,
              'Chemical_Constant' = chem_entries_final,
              'Ecological_Constant' = eco_entries,
              'Data_Filtering' = out_tbl2))
}
```

# Sankey diagram

A good way to visualize the steps of the data filtering process, and how many chemicals are removed at each step, is to use a Sankey diagram. The following function produces one, using the package `ggsankey`.

```{r}
make_sankey <- function(filt_dt,
                        use_has_conc = TRUE,
                        use_color = FALSE,
                        sankey_size = 3,
                        x_title_size = NA_real_){
  
  #simplified "is DLC" check:
  #yes/no,
  #rather than what kind of DLC it is
  filt_dt[, is_DLC2 := dplyr::case_when(is_DLC %in% c("is PCB",
                                                     "is dioxin",
                                                     "is furan") ~ "is DLC",
                                       .default = is_DLC)]
  filt_dt[, is_DLC2 := factor(is_DLC2,
                             levels = c("preloaded",
                                        "not DLC",
                                        "is DLC",
                                        "exclude"))]
  
  filt_dt <-  filt_dt %>% 
  #for readability, in case there is a long list of values missing or out of range --
#drop the derived ones (Br[...], Bv, BCF_beef, BCF_milk)
  mutate(missing_data = gsub(x = missing_data,
                             pattern = "B[rv]\\w*(,\\s)?",
                             replacement = "",
                             perl = TRUE)) %>%
  mutate(missing_data = gsub(x = missing_data,
                             pattern = "BCF_((beef)|(milk))(,\\s)?",
                             replacement = "",
                             perl = TRUE)) %>%
    mutate(missing_data = gsub(x = missing_data,
                             pattern = "RCF(,\\s)?",
                             replacement = "",
                             perl = TRUE)) %>%
    mutate(missing_data = gsub(x = missing_data,
                               pattern = "LogKow",
                               replacement = "Kow")) %>%
    mutate(missing_data =gsub(
                                               x = missing_data,
                                               pattern = "Da, Dw",
                                               replacement = "MV",
                                               fixed = TRUE
                                             )) %>%
    #if it's more than 3 items, split it onto two lines
    mutate(missing_data = if_else(sapply(strsplit(missing_data,
                                                  split = ", "),
                                         length) > 3,
                                 
                                           true = sapply(strsplit(missing_data,
                                                  split = ", "),
                                                  function(this_tmp){
                                                paste0(
                                                  paste(this_tmp[1:3],
                                                      collapse = ", "),
                                                    "\n",
                                                    paste(this_tmp[4:(length(this_tmp))],
                                                          collapse = ", ")
                                                )
                                                  }
                                                ),
                                  false = missing_data
                                         
                                  )) %>%
  mutate(missing_data = factor(missing_data,
                                      levels = c("preloaded",
                                      "no missing",
                                                 setdiff(unique(missing_data),
                                                         c("no missing",
                                                           "preloaded",
                                                           "exclude")),
                                                 "exclude")))
  
  if(use_has_conc %in% TRUE){
sankey_DT <- filt_dt %>% 
  make_long(has_conc_data,
                                   is_BST_preload,
                                   chemical_class,
                                   is_DLC2,
                                   missing_data,
                                   out_of_range_high,
                                   out_of_range_low,
                                   bad_format,
                                   Bulk_Upload,
                                   is_run)


lvls <- filt_dt[, Reduce(union,
                                            lapply(.SD, levels)),
                                   .SDcols = c("has_conc_data",
                                               "is_BST_preload",
                                               "chemical_class",
                                               "is_DLC2", 
                                               "missing_data",
                                               "out_of_range_high",
                                               "out_of_range_low",
                                               "bad_format",
                                               "Bulk_Upload",
                                               "is_run")]
  }else{
  sankey_DT <- filt_dt  %>% 
    make_long(is_BST_preload,
                                   chemical_class,
                                   is_DLC2,
                                   missing_data,
                                   out_of_range_high,
                                   out_of_range_low,
                                   bad_format,
                                   Bulk_Upload,
                                   is_run)

lvls <- filt_dt[, Reduce(union,
                                            lapply(.SD, levels)),
                                   .SDcols = c("is_BST_preload",
                                               "chemical_class",
                                               "is_DLC2", 
                                               "missing_data",
                                               "out_of_range_high",
                                               "out_of_range_low",
                                               "bad_format",
                                               "Bulk_Upload",
                                               "is_run")]
}
  lvls <- c("preloaded",
            "no missing",
            "no high",
            "no low",
            setdiff(lvls,
                    c("preloaded",
                      "no missing",
                      "no high",
                      "no low",
                      "exclude")),
            "exclude")


sankey_DT <- sankey_DT %>%
  mutate(node = factor(node,
                       levels = lvls),
         next_node = factor(next_node,
                            levels = lvls))


if(use_color %in% TRUE){
  #set up color palette
  if(use_has_conc %in% TRUE){
    cols <- viridis::viridis(n = length(lvls)-1)
    names(cols) <- lvls[!(lvls %in% "exclude")]
    cols <- c(cols,
              "exclude" = unname(cols["no conc data"])
    )}else{
      cols <- viridis::viridis(n = length(lvls))
      names(cols) <- lvls
    }
}

#label each node with its corresponding tally
#see also https://github.com/davidsjoberg/ggsankey/issues/9

node_labels <- sankey_DT %>%
  group_by(x, node) %>%
  tally() %>%
  tidyr::drop_na()

node_labels <- node_labels %>%
  mutate(node_label = paste0(node, "\n", n))

sankey_DT <- sankey_DT %>%
  left_join(node_labels,
            by = c('x', 'node'))

if(use_color %in% FALSE){
  this_plot <- ggplot(sankey_DT,
                      aes(x = x,
                          next_x = next_x,
                          node = node,
                          next_node = next_node,
                          label = node_label)) +
    geom_sankey(fill = "gray70",
                node.fill = "gray30") +
    geom_sankey_label(color = "black",
                      fill = "white",
                      size = sankey_size) 
    # geom_sankey_text(color = "black",
    #                  size = sankey_size)
}else{
this_plot <- ggplot(sankey_DT,
                      aes(x = x,
                          next_x = next_x,
                          node = node,
                          next_node = next_node,
                          fill = node,
                          label = node_label)) +
    geom_sankey() +
    geom_sankey_label(color = "white",
                      size = sankey_size) +
    # geom_sankey_text(color = "white",
    #                  size = sankey_size) +
    scale_fill_manual(values = cols)
}


this_plot <- this_plot +
  theme_sankey() +
  theme(legend.position = "none",
        axis.text.x = element_text(angle = 45,
                                   hjust = 1,
                                   size =  x_title_size),
        axis.title.x = element_blank())

return(this_plot)
}
```

# Run autoBST

## Bulk Upload tables and Sankey diagram for all chemicals with concentration set equal to 1 ppb

```{r}
biosolids_curated <- readxl::read_excel("data/Richman_et_al_2022_SI_Table1_Master List of Chemicals Detected in Biosolids.xlsx",
                                        "Biosolids Master List",
                                        "A1:D728")
#remove extra blank row
biosolids_curated <- biosolids_curated[2:nrow(biosolids_curated), ]

#also read in nutrients
biosolids_curated2 <- readxl::read_excel("data/Richman_et_al_2022_SI_Table1_Master List of Chemicals Detected in Biosolids.xlsx",
                                        "Nutrients Detected in Biosolids",
                                        "A1:D8")
#remove extra blank row
biosolids_curated2 <- biosolids_curated2[2:nrow(biosolids_curated2), ]

biosolids_curated <- data.table::rbindlist(list(biosolids_curated,
                                                biosolids_curated2),
                                           use.names = FALSE)

all_entries1ppb <- generate_tbls(DTXSIDs = biosolids_curated$DTXSID,
                             which_conc = "1ppb")

prepared1ppb <- list('Notes' = bulkuploader_Notes,
                           'tbl_Chemicals_import' = all_entries1ppb$tbl_Chemicals,
                           'Chemical_Constant_import' = all_entries1ppb$Chemical_Constant,
                  'Ecological_Constant_import' = all_entries1ppb$Ecological_Constant,
                           'Template_Chem_Org' = bulkuploader_Template_Chem_Org,
                           'Template_chem_Inorg' = bulkuploader_Template_Chem_Inorg,
                           'Template_Eco' = bulkuploader_Template_Eco,
                           'References' = bulkuploader_References,
                           'Data_Filtering' = all_entries1ppb$Data_Filtering)

writexl::write_xlsx(x = prepared1ppb,
                    path = "results/curated1ppb.xlsx")

my_plot1ppb <- make_sankey(filt_dt = all_entries1ppb$Data_Filtering,
                       use_has_conc = FALSE,
                       use_color = FALSE,
                       sankey_size = 2.7,
                       x_title_size = 8)

ggsave( "results/BST_sankey_1ppb.pdf",
        my_plot1ppb,
       height = 7,
       width = 9,
       units = "in")

ggsave( "results/BST_sankey_1ppb.png",
        my_plot1ppb,
       height = 7,
       width = 9,
       units = "in",
       dpi = 300)
```


## Bulk Upload tables and Sankey diagram for NSSS mean concentration data

```{r}
biosolids_curated <- readxl::read_excel("data/Richman_et_al_2022_SI_Table1_Master List of Chemicals Detected in Biosolids.xlsx",
                                        "Biosolids Master List",
                                        "A1:D728")
#remove extra blank row
biosolids_curated <- biosolids_curated[2:nrow(biosolids_curated), ]

#also read in nutrients
biosolids_curated2 <- readxl::read_excel("data/Richman_et_al_2022_SI_Table1_Master List of Chemicals Detected in Biosolids.xlsx",
                                        "Nutrients Detected in Biosolids",
                                        "A1:D8")
#remove extra blank row
biosolids_curated2 <- biosolids_curated2[2:nrow(biosolids_curated2), ]

biosolids_curated <- data.table::rbindlist(list(biosolids_curated,
                                                biosolids_curated2),
                                           use.names = FALSE)

all_entries_mean <- generate_tbls(DTXSIDs = biosolids_curated$DTXSID,
                             which_conc = "mean")

prepared_mean <- list('Notes' = bulkuploader_Notes,
                           'tbl_Chemicals_import' = all_entries_mean$tbl_Chemicals,
                           'Chemical_Constant_import' = all_entries_mean$Chemical_Constant,
                       'Ecological_Constant_import' = all_entries_mean$Ecological_Constant,
                           'Template_Chem_Org' = bulkuploader_Template_Chem_Org,
                           'Template_chem_Inorg' = bulkuploader_Template_Chem_Inorg,
                           'Template_Eco' = bulkuploader_Template_Eco,
                           'References' = bulkuploader_References,
                           'Data_Filtering' = all_entries_mean$Data_Filtering)

writexl::write_xlsx(x = prepared_mean,
                    path = "results/NSSSmean.xlsx")

 my_plot_mean <- make_sankey(filt_dt = all_entries_mean$Data_Filtering,
                       use_has_conc = TRUE,
                       use_color = FALSE,
                       sankey_size = 3.5,
                       x_title_size = 8)

ggsave("results/BST_sankey_NSSSmean.pdf",
       my_plot_mean, 
       height = 5,
       width = 9,
       units = "in")

ggsave("results/BST_sankey_NSSSmean.png",
       my_plot_mean, 
       height = 5,
       width = 9,
       units = "in")
```

## Bulk Upload tables and Sankey diagram for NSSS 95th percentile concentration data

```{r}
biosolids_curated <- readxl::read_excel("data/Richman_et_al_2022_SI_Table1_Master List of Chemicals Detected in Biosolids.xlsx",
                                        "Biosolids Master List",
                                        "A1:D728")
#remove extra blank row
biosolids_curated <- biosolids_curated[2:nrow(biosolids_curated), ]

#also read in nutrients
biosolids_curated2 <- readxl::read_excel("data/Richman_et_al_2022_SI_Table1_Master List of Chemicals Detected in Biosolids.xlsx",
                                        "Nutrients Detected in Biosolids",
                                        "A1:D8")
#remove extra blank row
biosolids_curated2 <- biosolids_curated2[2:nrow(biosolids_curated2), ]

biosolids_curated <- data.table::rbindlist(list(biosolids_curated,
                                                biosolids_curated2),
                                           use.names = FALSE)

all_entries_95 <- generate_tbls(DTXSIDs = biosolids_curated$DTXSID,
                             which_conc = "95th")

prepared_95 <- list('Notes' = bulkuploader_Notes,
                           'tbl_Chemicals_import' = all_entries_95$tbl_Chemicals,
                           'Chemical_Constant_import' = all_entries_95$Chemical_Constant,
                    'Ecological_Constant_import' = all_entries_95$Ecological_Constant,
                           'Template_Chem_Org' = bulkuploader_Template_Chem_Org,
                           'Template_chem_Inorg' = bulkuploader_Template_Chem_Inorg,
                           'Template_Eco' = bulkuploader_Template_Eco,
                           'References' = bulkuploader_References,
                           'Data_Filtering' = all_entries_95$Data_Filtering)

writexl::write_xlsx(x = prepared_95,
                    path = "results/NSSS95.xlsx")

 my_plot_95 <- make_sankey(filt_dt = all_entries_95$Data_Filtering,
                        use_has_conc = TRUE,
                       use_color = FALSE,
                       sankey_size = 3.5,
                       x_title_size = 8)

ggsave("results/BST_sankey_NSSS95th.pdf",
       my_plot_95, 
       height = 5,
       width = 9,
       units = "in")

ggsave("results/BST_sankey_NSSS95th.png",
       my_plot_95, 
       height = 5,
       width = 9,
       units = "in")
```

# Recommended procedure for using the Bulk Upload Tool and running the BST

Operating system requirements: Windows 7 or later.

Microsoft Access is required: any 32- or 64-bit installation of Access 2013 or later.

## Set up the BST

Download the Biosolids Screening Tool from [https://www.epa.gov/biosolids/risk-assessment-pollutants-sewage-sludge](https://www.epa.gov/biosolids/risk-assessment-pollutants-sewage-sludge). (Direct link: [https://www.epa.gov/system/files/other-files/2023-03/biosolids-screening-tool.zip](https://www.epa.gov/system/files/other-files/2023-03/biosolids-screening-tool.zip).)

Before unzipping the BST, create a directory with as short a pathname as possible. The BST must be run in a directory with a pathname of 48 characters or less, according to the BST User Guide. This probably means you need to create the directory close to the root of `C:\`. For example, Dr. Kruse created the directory at `C:\Users\pkruse\BST`.

Unzip the BST download into your short-pathname directory. A new folder called "BST" will appear.

Within the short-pathname directory, create one subdirectory for each set of input concentrations: 1 ppb; NSSS mean; and NSSS 95th percentile. Name them (for example) `ppb`, `mean`, and `p95`, respectively, to keep track of which is which.

Within each of these subdirectories, copy and paste the *contents* of the `BST` folder. You don't need to copy and paste the `HELP` folder every time, but you do need to copy and paste the other directories: `HWIR`, `RISK_RESULTS`, `SOURCE_RESULTS`, `VVWM`. These directories contain helper files that the Microsoft Access implementation of the BST needs in order to run. 

The MS Access implementation of the BST itself lives in the `.mdb` file. You do not need to copy and paste the `.mdb` file; it will be replaced in the next step anyway.

## Set up the version of the BST with bulk upload capabilities

The MS Access implementation of the BST downloaded from the URL above does not include bulk upload capabilities. The file `BST_BulkUpload.mdb`, in the same directory as this R Markdown file, is the version of the BST with bulk upload capabilities. 

Now, within each subdirectory (`mean`, `p95`, and `ppb`), make a new copy of the file `BST_BulkUpload.mdb`.

Optionally, you may want to rename these files to make it clear which subdirectory they are associated with, e.g., as `BST_BulkUpload_mean.mdb`, `BST_BulkUpload_p95.mdb`, and `BST_BulkUpload_ppb.mdb`.

## Place the Bulk Upload worksheet (output of autoBST)

Now, copy and paste the Bulk Upload workbooks generated by the autoBST into their corresponding folders. By default, the above code will have saved them in the `results/` subdirectory that should be alongside this R Markdown document in the current R working directory. Copy and paste `results/curated1.xlsx` into `ppb`; `results/NSSSmean.xlsx` into `mean`; and `results/NSSS95th.xlsx` into `p95`.

Within each of the subdirectories, rename the `.xlsx` file to `ChemicalImport.xlsx`. The BST requires that the bulk upload file have this exact filename.

## Set up the BST runs

Now, choose a subdirectory to start with: for example, the `mean` subdirectory. Double-click on the `.mdb` file you just created to open it in Microsoft Access.

You may receive a pop-up message saying "The active content in this file is blocked. Review your Trust Center settings or contact your IT administrator." Just click OK.

Just underneath the menu ribbon, you may see a yellow bar that says "SECURITY WARNING: Some active content has been disabled. Click for more details." followed by a button that says "Enable Content." Click the "Enable Content" button. This allows the BST to run the macros it needs to import and run simulations. The yellow bar should then disappear.

Read the message titled "Welcome to the Biosolids Tool." Then click the button that says "Continue" at the bottom.

Next, you will see a window titled "Biosolids Tool (Version 1.02) - Main Menu." At the bottom of this window are three buttons: "Configure," "Run", and "View Results." Click the "Configure" button.

You will see a window titled "Configure Model." At the top, it shows several tabs: Scenarios, Chemicals, Human Exposure, Ecological Exposure, and Inputs.

Click on the "Chemicals" tab (this should be the second tab from the left, after "Scenarios"). You will now see the pre-loaded chemicals. 

Also, make sure you are looking at the version of the BST that includes bulk-upload capabilities: **without clicking any buttons**, look at the column of buttons on the left and check to make sure that you *see* a button labeled "Bulk Import Chemicals" (**do not click it yet**).

### Manually edit the hazard/toxicity values for the pre-loaded chemicals 

Manually edit the hazard/toxicity values for the pre-loaded chemicals: set CSForal, IUR, RfC, and RfD equal to 1 for all pre-loaded chemicals. This makes it so that all the "risk" and "hazard quotient" values output by the BST actually represent exposure doses or concentrations. This is the only way to get the BST to output average daily dose that is the total of all ingestion pathways or inhalation pathways. 

If you want the original pre-loaded values for CSForal, IUR, RfC, and RfD, they are still populated in the original file `BST_BulkUpload.mdb`.

### Edit pre-loaded biosolids concentrations

In the subdirectory `ppb`, for simulations with 1 ppb biosolids concentrations, you will need to edit the preloaded chemicals so that `CTPWasteDry` takes a value of $1 \times 10^{-3}$ mg/kg (equivalent to 1 ppb).

When running simulations with mean NSSS biosolids concentrations, you will need to edit the preloaded chemicals so that `CTPWasteDry` reflects the mean NSSS concentration.

When running simulations with 95th percentile NSSS biosolids concentrations, you do *not* need to edit the preloaded chemicals. For the preloaded chemicals, `CTPWasteDry` by default reflects the 95th percentile NSSS concentration, except for a few chemicals specified in the next section.

## De-select pre-loaded chemicals that do not have NSSS mean or 95th percentile concentrations

When running simulations with mean or 95th percentile biosolids concentrations, there are some preloaded chemicals that do not have an NSSS mean or 95th percentile. All except a few of these were measured in NSSS, but have too few detects to compute a mean or 95th percentile. 

Here are the exceptions: PFOA and PFOS were not measured in NSSS. Chromium (III) and Chromium (VI) were not speciated separately in TNSSS; only total Chromium was measured. Divalent mercury was not speciated separately in TNSSS; only mercury was measured (elemental mercury). Selenium (IV) and Selenium (VI) were not speciated separately in TNSSS; only total Selenium was measured.

Here is the list of chemicals that should be deselected:

- Albuterol
- Chlortetracycline
- Chromium (III)
- Chromium (VI) 
- Mercury (divalent) 
- Oxolinic acid
- PFOA
- PFOS
- Selenium (IV)
- Selenium (VI)
- Sulfathiazole

These chemicals should be deselected when you are running simulations with mean or 95th percentile biosolids concentrations.

## Performing the Bulk Import

First, save the BST file with only the preloaded chemicals, including any edits you have made to the preloaded chemicals, and close out of it.

We highly recommend keeping a "clean" version of the BST file as it was just before you performed the bulk import, including any edits that you have made manually to the pre-loaded chemicals. This is because **you cannot delete or overwrite data for any chemicals once they are imported**. If you realize that you need to change something in the autoBST and re-generate new Bulk Upload worksheets with new data, then it will be much easier if you can simply make a new copy of the file and re-do the bulk import with the new workshets.

Next, create a copy of the "clean" file, and rename it. You can use whatever name you like, as long as it fits into the 48-character path name length limit.

Open this newly-renamed Access file. This is where you will perform the bulk upload.

Go to the "Chemicals" tab. On the left, you will see several buttons arranged vertically. Find the button that says "Bulk Import Chemicals." (You may need to scroll down inside this window before this button is visible.)

Click the button that says "Bulk Import Chemicals." The BST will instantly locate the file `ChemicalImport.xlsx` in the same directory and import all chemical data from that file.

Afterwards, you should see many chemicals added to the list of chemicals showing in this window. All of the new chemicals will have the "User-Added" checkbox checked.

To confirm that import went correctly, you can click the "View/Edit/Delete" button for one or a few chemicals, and confirm that the data shown matches the data in the output of the BST (`ChemicalImport.xlsx`).

## Finalizing run settings

Now, choose the model inputs you want to use on the other tabs. Please see the BST User Guide for more information about the various scenarios that can be run.

To reproduce our simulations, make the following selections:

### "Scenarios" tab
On the "Scenarios" tab, where it says "Select Unit Type," check the box for "LAU." Where it says "Select Climate", click the radio button for "Average." Under "Land Application Options", where it says "Select Scenario," check the boxes for "Crop," "Pasture," and "Reclamation." The dry application rates should take their default values: 10 for Crop, 10 for Pasture, 50 for Reclamation. 

### "Chemicals" tab

The chemicals tab was set up in the previous steps.

### "Human Exposure" tab

Ensure checkboxes for all pathways are checked.

### "Ecological Exposure" tab

Click the button for "Deselect All" (on the left). 

### "Inputs" tab

Leave everything as-is.

## Running the BST

Once you are finished choosing input parameters, click the button in the upper right that says "Close Form" in red text. Your choices will be saved. You will be returned to the main Biosolids Tool window with three buttons at the bottom: "Configure," "Run", and "View Results."

Click the "Run" button. You will be asked whether you are sure you want to run the model. If you are sure (that is, you are sure that all settings are what you want), click OK.

The model will now run. For 174 chemicals, it took several hours on a Dell Latitude 5340 PC with 16 GB RAM and processor 13th Gen Intel(R) Core(TM) i7-1365U 1.80 GHz. 

### Practical note about BST runs

It is OK to keep using your computer to do other things while the BST is running. However, as the model runs for each chemical, it spawns various command-prompt windows that each grab keyboard/mouse focus temporarily. Don't worry, you won't accidentally issue a command that breaks the simulation. But it is mildly annoying, because it will repeatedly interrupt whatever you are typing in another window.

Optionally, you can avoid this minor annoyance by running the BST at a time when no one is using the computer for anything else (for example, overnight).

## Saving the BST run results

When the simulation is done, you will see a new pop-up window inside the BST that says "Done!" Click OK.

Click "View Results", click "LAU Results," and export the results to Excel.

Rename the exported Excel workbook files. For example, for the NSSS mean runs for the LAU scenarios, you might want to name the file "BST_results_LAU_mean.xlsx."

## Moving the BST run results for analysis

Copy and paste the exported BST result spreadsheets into the `results/` subdirectory that should be in the same directory alongside this R Markdown file.

That directory already contains the exported BST result spreadsheets from the BST runs performed for "A Cheminformatics Workflow for Higher-throughput Modeling of Chemical Exposures from Biosolids" (Kruse et al. 2025). The output file for the NSSS mean concentration scenario is in `results/BST_results_LAU_mean.xlsx`; the output file for the NSSS 95th percentile concentration scenario is in `results/BST_results_LAU_mean_p95.xlsx`; and the output file for the 1 ppb concentration scenario is in `results/BST_results_LAU_mean_ppb.xlsx`.

The following code refers to those pre-existing output files.

# Analysis of BST results

Make an `analysis/` directory to hold the results of the analysis, if it does not already exist.

```{r}
if(!dir.exists("analysis")){
  mkdir("analysis")
}
```

Read in all output data.

```{r}
  df_list <- sapply(c("mean",
           "p95",
           "ppb"),
         function(which_conc){
  this_out <- readxl::read_excel(paste0(
    "results/BST_results_LAU_",
    which_conc,
    ".XLSX"),
                               sheet = "Human_Results")
  
  #Filter to adult daily dose
  this_add <- this_out %>%
  dplyr::filter(Receptor %in% 'Adult Farmer' &
                  `Health Benchmark` %in% "Noncancer" &
                  Pathway %in% 'Total Ingestion')
  return(this_add)
         },
  simplify = FALSE,
  USE.NAMES = TRUE)
  
  df <- dplyr::bind_rows(df_list,
                         .id = "which_conc")
```

Ensure any chemicals that should have been removed from the NSSS scenarios are removed.

```{r}
df <- df %>%
  filter(!(which_conc %in% c("mean", "p95") &
             `Chemical Name` %in% c(
               "Chromium (III)",
               "Chromium (VI)",
               "Mercury (divalent)",
               "Selenium (IV)",
               "Selenium (VI)",
               "Albuterol",
               "Chlortetracycline",
               "Oxolinic acid",
               "PFOA",
               "PFOS",
               "Sulfathiazole"
             )
           )
         )
```

  
Plot frequency polygons (like histograms) for the adult farmer ADDs, under the three LAU scenarios and the three biosolids concentration scenarios.

```{r}
  
  #frequency polygon of ADDs
  ggplot(df) +
  geom_freqpoly(aes(x = Value),
                bins = 30,
                linewidth = 1) +
    facet_grid(rows = vars(which_conc),
               cols = vars(Scenario),
               scales = "free") +
  scale_x_log10(guide = "axis_logticks") +
  xlab("BST-predicted total ingestion ADD (adult farmer), mg/kg/day") +
  theme_bw() +
  theme(strip.background = element_blank(),
        strip.text = element_text(size = 14),
        strip.text.y.right = element_text(size = 14,
                                          angle = 0),
        axis.text = element_text(size = 12),
        axis.title = element_text(size = 12))

ggsave(paste0("analysis/BST_ADD_freqpoly.pdf"),
        width = 9.75,
       height = 5.4)
ggsave(paste0("analysis/BST_ADD_freqpoly.png"),
       width = 9.75,
       height = 5.4,
       units = "in",
       dpi = 300)
```

Range of concentrations

```{r}
df %>%
  group_by(which_conc, Scenario) %>%
  summarise(min_ADD = min(Value, na.rm = TRUE),
            max_ADD = max(Value, na.rm = TRUE))
```


Compare biosolids concentrations to adult farmer ADDs for each LAU scenario and each biosolids concentration scenario (excluding 1 ppb, since concentration is constant for that scenario). Make plots and run linear regression (on a log10-log10 scale).

```{r}
#read in mapping between DTXSID and stripped CASRN
#pull this from the 1 ppb spreadsheet
mapping <- readxl::read_excel("results/curated1ppb.xlsx",
                   sheet = "Data_Filtering") %>%
  dplyr::select(c(DTXSID, casrn_stripped))

conc_tbl_list <- sapply(c("mean",
                     "p95",
                     "ppb"),
                   function(which_conc){
                     print(which_conc)
                     if(which_conc %in% "mean"){
                       
                       this_tbl <- make_conc_tbl(which_conc = "mean") %>%
                         dplyr::select(DTXSID, conc_mgkg) %>%
                         left_join(mapping) %>%
                         dplyr::rename(CAS = casrn_stripped) %>%
                         dplyr::mutate(CAS = as.numeric(CAS))
                     }else if(which_conc %in% "p95"){
                       this_tbl <- make_conc_tbl(which_conc = "95th") %>%
                         dplyr::select(DTXSID, conc_mgkg) %>%
                         left_join(mapping) %>%
                         dplyr::rename(CAS = casrn_stripped) %>%
                         dplyr::mutate(CAS = as.numeric(CAS))
                     }else{ #if which_conc == "ppb"
                       this_tbl <- mapping %>%
                         dplyr::mutate(conc_mgkg = 1e-3) %>%
                         dplyr::rename(CAS = casrn_stripped)
                     }
                     
                     this_tbl <- this_tbl %>%
                       dplyr::mutate(CAS = as.numeric(CAS))
                     
                     return(this_tbl)
                   },
simplify = FALSE,
USE.NAMES = TRUE)

conc_tbl <- dplyr::bind_rows(conc_tbl_list,
                             .id = "which_conc")


#get concentration rankings
conc_tbl <- conc_tbl %>%
  dplyr::arrange(which_conc,
                 -conc_mgkg) %>%
  dplyr::group_by(which_conc) %>%
  dplyr::mutate(conc_rank = 1:dplyr::n()) %>%
  dplyr::ungroup()

#merge with ADD info
df_conc <- df %>%
  dplyr::left_join(conc_tbl,
                   relationship = "many-to-one") %>%
  dplyr::mutate(conc_mgkg = dplyr::if_else(which_conc %in% "ppb" &
                                             is.na(conc_mgkg),
                                           0.001,
                                           conc_mgkg)
                )

#Regression of log10 ADD on log10 conc.
df_lm <- df_conc %>%
  dplyr::filter(!(which_conc %in% "ppb")) %>%
  dplyr::group_by(which_conc,
                  Scenario) %>%
  dplyr::summarise(Rsq = summary(
    lm(
      log10(Value) ~ log10(conc_mgkg)
      )
    )$r.squared)

write.csv(df_lm,
          "analysis/Rsquared_logADD_logconc.csv")
df_lm <- df_lm %>%
  dplyr::mutate(label = sprintf("R^2 == %.3f",
                         Rsq))

#plot ADD vs. concentration

ggplot(df_conc %>%
         filter(!(which_conc %in% "ppb" &
                    !is.na(Value))),
       aes(x = conc_mgkg,
                    y= Value)) +
  geom_point(shape = 21) +
    geom_text(data = df_lm,
            aes(x = 0,
                y = Inf,
                label = label),
            parse = TRUE,
            hjust = -0.3,
            vjust = 1.8) +
  facet_grid(rows = vars(which_conc),
               cols = vars(Scenario)) +
  scale_y_log10(guide = "axis_logticks") +
  scale_x_log10(guide = "axis_logticks") +
  geom_smooth(method = "lm", se = FALSE, color = "gray50") +
  xlab("Biosolids conc., mg/kg") +
  ylab("Adult farmer ADD, mg/kg/day") +
  theme_bw() + 
  theme(strip.background = element_blank(),
        strip.text = element_text(size = 14))

ggsave("analysis/ADD_vs_conc.pdf",
       width = 9.75,
       height = 5.4,
       units = "in")
ggsave("analysis/ADD_vs_conc.png",
        width = 9.75,
       height = 5.4,
       units = "in",
       dpi = 300)
```

Add Log Kow to the regression of adult farmer ADD on concentration (by LAU scenario and by biosolids concentration scenario).

```{r}

#plot ADD vs. concentration gray-scaled by log Kow
#get log Kow information
#this will be pulled from the mean results workbook,
#but it doesn't matter which results workbook we use
this_input <- readxl::read_excel(
  "results/BST_results_LAU_mean.XLSX",
  sheet = "Run_ChemicalInputs") %>%
  dplyr::filter(Model_Code %in% "LogKow") %>%
  dplyr::select(CAS, Value) %>%
  dplyr::rename(LogKow = Value) %>%
  dplyr::distinct()
 
 df_conc2 <- df_conc %>%
   dplyr::left_join(this_input,
                    by = "CAS") %>%
   dplyr::mutate(LogKow = as.numeric(LogKow))
 
 ggplot(df_conc2 %>%
         filter(!(which_conc %in% "ppb"))) +
  geom_point(shape = 21,  aes(x = conc_mgkg,
                    y= Value,
           color = LogKow)) +
    geom_text(data = df_lm,
            aes(x = 0,
                y = Inf,
                label = label),
            parse = TRUE,
            hjust = -0.3,
            vjust = 1.8) +
  facet_grid(rows = vars(which_conc),
               cols = vars(Scenario)) +
  scale_y_log10(guide = "axis_logticks") +
  scale_x_log10(guide = "axis_logticks") +
   scale_color_distiller(palette = "Greys", trans = "reverse",
                      guide = guide_colorbar(reverse = TRUE)) +
  geom_smooth( aes(x = conc_mgkg,
                    y= Value), method = "lm", se = FALSE, color = "black") +
  xlab("Biosolids conc., mg/kg") +
  ylab("Adult farmer ADD, mg/kg/day") +
  theme_bw() + 
  theme(strip.background = element_blank(),
        strip.text = element_text(size = 14))

ggsave("analysis/ADD_vs_conc_logkow.pdf",
        width = 9.75,
       height = 5.4)
ggsave("analysis/ADD_vs_conc_logkow.png",
       width = 9.75,
       height = 5.4,
       units = "in",
       dpi = 300)

df_lm2 <- df_conc2 %>%
    dplyr::filter(!(which_conc %in% "ppb")) %>%
    dplyr::group_by(which_conc,
                    Scenario) %>%
    dplyr::summarise(Rsq = summary(
        lm(
            log10(Value) ~ log10(conc_mgkg) + LogKow
        )
    )$r.squared)

write.csv(df_lm2,
          "analysis/Rsquared_logADD_logconc_logKow.csv")
```

Get rankings of chemicals by adult farmer ADD within LAU scenario and within biosolids concentration scenario.

```{r}
df_rank <- df_conc %>%
  dplyr::arrange(which_conc,
           Scenario,
          -Value) %>%
  dplyr::group_by(which_conc,
           Scenario) %>%
  mutate(Rank = 1:n()) %>%
  ungroup() %>%
  rename(ADD = Value) %>%
  select(c(DTXSID, CAS, `Chemical Name`, which_conc, conc_mgkg, conc_rank, Scenario, ADD, Rank)) %>%
  rename(Adult_Farmer_ADD = ADD) %>%
 tidyr::pivot_wider(id_cols = c(DTXSID, CAS, which_conc, `Chemical Name`, conc_mgkg, conc_rank),
              names_from = Scenario,
              values_from = c(Adult_Farmer_ADD,
                              Rank)) %>%
  dplyr::mutate(conc_rank = dplyr::if_else(which_conc %in% "ppb",
                                           NA_real_,
                                           conc_rank))

  write.csv(df_rank, "analysis/ADD_rank.csv")
```
````
