## Supplementary material for "A Cheminformatics Workflow for Higher-throughput Modeling of Chemical Exposures from Biosolids": autoBST workflow files: classyfireAPIs.R

```
#'InChIKey classification
#'
#'This function uses the ClassyFire API to classify chemicals from an input
#'data.table using the InChIKey chemical identifier.
#'
#'This function queries ClassyFire's lookup table of pre-classified InChiKeys
#'
#'@param inchikeys A vector of InCHiKeys to be classified
#'@param tax_level_labels By default, the list of taxonomy levels for
#'  ClassyFire: \code{kingdom, superclass, class, subclass, level5, ...
#'  level11}.
#'@param requests_min How many queries per minute can be sent to the ClassyFire
#'  API server. At present, the ClassyFire API limits queries to 12 per minute,
#'  so the default value of this argument is 12.
#'@return A data.frame with the following variables: \itemize{
#'  \item{identifier: The input InCHiKey that was queried. For example,
#'  "XGQJGMGAMHFMAO-UHFFFAOYSA-N"}
#'  \item{smiles: The corresponding SMILES returned by ClassyFire, if any}
#'  \item{inchikey: The InCHiKey returned by ClassyFire, if any. Will be of
#'  format "InCHiKey=XGQJGMGAMHFMAO-UHFFFAOYSA-N"}
#'  \item{classification_version: The version number returned by ClassyFire.
#'  For example, "2.1"}
#'  \item{level: The names of all levels in this InCHiKey's classification.
#'  Will be elements of \code{tax_level_labels}.}
#'  \item{name: The name or label of each level in this InCHiKey's
#'  classification.}
#'  \item{report: A text string reporting the status of the classification,
#'  "ClassyFire returned a classification" if successful; otherwise the report
#'  returned by ClassyFire, or a report about an internal error.}
#'  }
#'
#'  Note that the data.frame is in "long" format, with multiple rows for each
#'  InCHiKey. There is one row for each level of classification for each
#'  InCHiKey. For example, the classification for InCHiKey
#'  "XGQJGMGAMHFMAO-UHFFFAOYSA-N" terminates at level 5, so there will be five
#'  rows for that InCHiKey. The classification for InCHiKey
#'  "PZNXLZZWWBSQQK-UHFFFAOYSA-N" terminates at level 4 (subclass), so there
#'  will be four rows for that InCHiKey.
#'
#'  For use with \code{treecompareR} functions that expect a `data.frame` of
#'  classified entities, this `data.frame` will need to be reshaped into wider
#'  format, with one row for each InCHiKey and one column for each level of
#'  classification. This can be done, e.g., using \code{tidyr::pivot_wider(dat,
#'  names_from = "level", values_from = "name")} (where \code{dat} is the
#'  returned `data.frame`.) However, be on the lookout for pathological cases
#'  where an InCHiKey is listed with two different labels at the same level.
#'  These occur rarely, but they do occur. \code{tidyr::pivot_wider()} will
#'  throw a warning if this happens -- pay attention to it!
#'
#'  Note also that the returned data.frame includes only unique, valid
#'  InCHiKeys. Any duplicates, blanks, NAs, or anything that is not a valid
#'  InCHiKey according to `webchem::is.inchikey()`is not queried, and is not
#'  included in the output.
#'
#'  If you have a source `data.frame` with duplicate, missing, or invalid
#'  InCHiKeys, you can merge the returned `data.frame` with it (or pivot wider,
#'  then merge). For example, if your source `data.frame` is called
#'  \code{source_dat} with variable \code{"INCHIKEY"} containing the InCHiKeys,
#'  and the returned `data.frame` is in variable \code{dat}, the following code
#'  will do the merge: \code{dplyr::left_join(source_dat, dat, by = "INCHIKEY" =
#'  "identifier")}. In that case, classification columns will be filled with NA
#'  for any missing or invalid InCHiKeys.
#'
#'@export
#'@references Djoumbou Feunang, Y., Eisner, R., Knox, C., Chepelev, L.,
#'  Hastings, J., Owen, G., . . . Wishart, D. S. (2016). ClassyFire: automated
#'  chemical classification with a comprehensive, computable taxonomy. J
#'  Cheminform, 8, 61. https://doi.org/10.1186/s13321-016-0174-y
#'
#'

classify_inchikeys <- function(inchikeys,
                               tax_level_labels = c('kingdom',
                                                    'superclass',
                                                    'class',
                                                    'subclass',
                                                    'level5',
                                                    'level6',
                                                    'level7',
                                                    'level8',
                                                    'level9',
                                                    'level10',
                                                    'level11'),
                               requests_per_min = 12){


  INCHIKEYS <- unique(inchikeys) #save time by removing duplicates
  
  wait_min <- 60/requests_per_min #minimum time to wait between requests (in seconds)

  #get classifications as a list of data.tables, one for each INCHIKEY
  entities_list <- lapply(
    INCHIKEYS,
    function(this_inchikey){
      #if inchikey is NA, blank, or not in valid InChiKey format,
      #do not query API; return NA classification instead
      if(is.na(this_inchikey) || #if InChiKey is NA
         !nzchar(trimws(this_inchikey)) || #if InChiKey is blank
         !webchem::is.inchikey(this_inchikey, #if not valid InChiKey format
                               type = "format")
         ){
        #placeholder blank output list with all expected elements
        output <- list()
      }else{ #if inchikey is valid, query ClassyFire API
        Sys.sleep(wait_min) #pause minimum number of seconds before querying again
        output <- query_classyfire_inchikey(inchikey = this_inchikey,
                                            wait_min = wait_min)
        output$identifier <- this_inchikey
      }
      return(output)
    } #end function to apply to each INCHIKEY
  )

  class_list <- lapply(entities_list,
                       parse_classified_entities,
                       tax_level_labels = tax_level_labels)

  #now rowbind the list of data.frames to get one big data.frame
  inchi_class <- dplyr::bind_rows(class_list)

  #coerce to wide format
  
  new_class <- inchi_class %>%
    tidyr::pivot_wider(id_cols = c(identifier,
                       classification_version,
                       report),
                       names_from = level,
                       values_from = c(name)
    ) 
  
  #reorder as input, re-inserting any duplicates
  inchi_tbl <- data.frame(identifier = inchikeys)
  
  new_class <- inchi_tbl %>%
    dplyr::left_join(new_class,
                     by = "identifier") %>%
    dplyr::rename(inchikey = identifier)
  
  return(new_class)
}

#' Parse classified entities
#'
#' @param entities JSON element of classified entities as returned by ClassyFire
#'   API query
#' @param tax_level_labels ChemOnt taxonomy level labels 
# @param
#' @return A data.frame consisting of rows corresponding to each classified
#' entry from the `entities` input. A data.frame with one row for each entity identifier, and variables
#'  `identifier`, `smiles`, `inchikey`, `kingdom`, `superclass`, `class`,
#'   `subclass`, `level5`, `level6`, ... `level11`, `classification_version`,
#'    and `report`.
parse_classified_entities <- function(entities,
                                      tax_level_labels = c('kingdom',
                                                           'superclass',
                                                           'class',
                                                           'subclass',
                                                           'level5',
                                                           'level6',
                                                           'level7',
                                                           'level8',
                                                           'level9',
                                                           'level10',
                                                           'level11')){
  identifier <- NULL
  level <- NULL
  name <- NULL
  #check to see whether classifications actually exist for these
  if(length(entities)==0){
    #if no classifications, return empty data.frame,
    #but with the expected variable names
    classified_entities <- data.frame(identifier = character(0),
                                      smiles = character(0),
                                      inchikey = character(0),
                                      #kingdom = character(0),
                                      #superclass = character(0),
                                      #class = character(0),
                                      #subclass = character(0),
                                      #level5 = character(0),
                                      #level6 = character(0),
                                      #level7 = character(0),
                                      #level8 = character(0),
                                      #level9 = character(0),
                                      #level11 = character(0),
                                      classification_version = character(0),
                                      level = character(0),
                                      name = character(0),
                                      report = character(0))
  }else{ #if length(entities)>0
    #expected named items in entities and their expected classes:
    # identifier                    smiles                  inchikey
    # "character"               "character"               "character"
    # kingdom                superclass                     class
    # "data.frame"              "data.frame"              "data.frame"
    # subclass        intermediate_nodes             direct_parent
    # "data.frame"                    "list"              "data.frame"
    # alternative_parents       molecular_framework              substituents
    # "list"               "character"                    "list"
    # description      external_descriptors                 ancestors
    # "character"                    "list"                    "list"
    # predicted_chebi_terms predicted_lipidmaps_terms    classification_version
    # "list"                    "list"               "character"


    #get identifiers, smiles, inchikey
    #these will be character vectors identifying entities

    #note that some of these may be character(0) -- handle
    #by filling with NAs
    for(item in c("identifier",
                  "smiles",
                  "inchikey",
                  "classification_version")){
      if(length(entities[[item]])==0){
        entities[[item]] <- NA_character_
      }
    }

    cf <- as.data.frame(entities[c("identifier",
                         "smiles",
                         "inchikey",
                         "classification_version")])

    #kingdom, superclass, class, subclass are all data.frames
    #with one row for each identifier,
    #and variables name, description, chemont_id, url
    #giving the relevant ChemOnt taxonomy label for each identifier
    #(i.e. "name" in the "kingdom" element gives the kingdom for each identifier,
    #"name" in the "superclass" element gives the superclass for each identifier,
    #etc.)
    #rowbind these

    #note: if no classification for one of these four levels, the item may be NULL.
    #handle accordingly.
    for(item in c("kingdom",
                  "superclass",
                  "class",
                  "subclass")){
      #check if NULL
      if(is.null(entities[[item]])){
        #replace with list with NAs
        entities[[item]] <- list(name = NA_character_,
                                 description = NA_character_,
                                 chemont_id = NA_character_,
                                 url = NA_character_)
      }

      #check if character(0) and replace with NA
      for (subitem in names(entities[[item]])){
        if(length(entities[[item]][[subitem]])==0){
          entities[[item]][[subitem]] <- NA_character_
        }
      }
    }

    cf_class1 <- dplyr::bind_rows(as.list(entities[c("kingdom",
                                                       "superclass",
                                                       "class",
                                                       "subclass")]))
    #add a column for the identifiers -- repeat for each taxonomy level
    cf_class1$identifier <- rep(entities$identifier,
                                4)

    #"intermediate_nodes"
    #is a list with one element for each item in "input",
    #a 1-row data.frame with variables name, description, chemont_id, url
    #giving more-specific levels of classification, if any.
    #if no intermediate nodes for an entity,
    #its corresponding element in "intermediate_nodes" is an empty data.frame.
    #if it is an empty list (as happens for single-InChiKey queries without intermediate nodes)
    #then handle accordingly.
    cf_class2 <- entities$intermediate_nodes
    if(length(cf_class2)>0){
    names(cf_class2) <- entities$identifier
    cf_class2_df <- dplyr::bind_rows(cf_class2, .id = "identifier")
    }else{
    cf_class2_df <- data.frame()
    }
    rm(cf_class2)

    #rowbind the kingdom-thru-superclass labels,
    #and the "intermediate nodes" labels.
    cf_class <- dplyr::bind_rows(cf_class1,
                                 cf_class2_df)

    # element 'direct_parent' is a data.frame
    #with one row for each identifier,
    #and variables name, description, chemont_id, url
    direct <- entities$direct_parent
    direct$identifier <- entities$identifier
    cf_class <- dplyr::bind_rows(cf_class, direct)

    #"direct parent" is the terminal label,
    #and may be a duplicate of kingdom, superclass, class, subclass,
    #or something in "intermediate nodes",
    #or may not be a duplicate at all.
    #If it duplicates another row, remove it;
    #otherwise, don't.

      cf_class <- cf_class %>%
        dplyr::distinct() %>%
        dplyr::filter(!is.na(name))

    #Add taxonomy level names
    #e.g. 1, 2, 3, 4 -> kingdom, superclass, class, subclass
    cf_class <- cf_class %>%
      dplyr::mutate(level = tax_level_labels[1:nrow(cf_class)])

    classified_entities <- cf_class
    #add a "report" column
    if(!"report" %in% names(entities)){
    classified_entities$report <- "ClassyFire returned a classification"
    }else{
      classified_entities$report <- entities$report
    }

    #merge in smiles and inchikey
    classified_entities <- dplyr::left_join(cf,
                                            classified_entities,
                                            by = "identifier") %>%
      as.data.frame()
  } #end if length(entities)>0

  return(classified_entities)
}


#' Query ClassyFire InChIKey
#'
#' This is a helper function that is used to communicate with the ClassyFire API
#' in order to retrieve classifications.
#'
#' @param inchikey A character string encoding the InChIKey of the chemical
#'   under query.
#' @param retry_get_times The number of times to retry the query, a positive
#'   integer with default value 3.
#' @param wait_min The number of seconds to pause in between attempts, used in
#'   the \code{\link{httr}{RETRY}} function.
#'
#' @return A JSON file with the classification data.
query_classyfire_inchikey <- function(inchikey,
                                      retry_get_times = 3,
                                      wait_min = 0.5){

  base_url <- "http://classyfire.wishartlab.com/entities"
  url <- paste0(base_url,
                "/",
                inchikey,
                ".json")
  #try getting results for query
  resp <- httr::RETRY(verb = "GET",
                      url = url,
                      encode = "json",
                      times = retry_get_times,
                      pause_base = wait_min,
                      pause_min = wait_min,
                      terminate_on = c(404))

  json_res <- httr::content(resp, "text")
  #json_res will be NULL if no content was returned
  if(!is.null(json_res)){
    json_parse <- jsonlite::fromJSON(json_res, simplifyDataFrame = FALSE)
  }else{
    json_parse <- NULL
  }

  #initialize a placeholder json_parse in case everything else fails
  #the items that are named lists are usually actually data.frames,
  #but lists are coerceable to data.frames and making it a list
  #helps with making it more general.
  json_parse_default <- list("identifier" = character(0),
                                   "smiles" = character(0),
                                   "inchikey" = character(0),
                                   "kingdom" = list("name" = character(0),
                                                          "description" = character(0),
                                                          "chemont_id" = character(0),
                                                          "url" = character(0)),
                                   "superclass" = list("name" = character(0),
                                                             "description" = character(0),
                                                             "chemont_id" = character(0),
                                                             "url" = character(0)),
                                   "class" = list("name" = character(0),
                                                        "description" = character(0),
                                                        "chemont_id" = character(0),
                                                        "url" = character(0)),
                                   "subclass" = list("name" = character(0),
                                                           "description" = character(0),
                                                           "chemont_id" = character(0),
                                                           "url" = character(0)),
                                   "intermediate_nodes" = list(),
                                   "direct_parent" = list("name" = character(0),
                                                                "description" = character(0),
                                                                "chemont_id" = character(0),
                                                                "url" = character(0)),
                                   "classification_version" = character(0))

  #fill in any elements in json_parse_default not in json_parse
  #otherwise, keep elements in json_parse
  json_parse <- c(
    json_parse,
    json_parse_default[
      setdiff(
        names(json_parse_default),
        names(json_parse)
      )
    ]
  )

  #assign identifier as inchikey
  json_parse$identifier <- inchikey


  return(json_parse)
}
```
