## Supplementary material for "A Cheminformatics Workflow for Higher-throughput Modeling of Chemical Exposures from Biosolids": autoBST workflow files: README.md

# autobst

This repository contains files associated with the AutoBST workflow associated with the manuscript "A Cheminformatics Workflow for Higher-throughput Modeling of Chemical Exposures from Biosolids" (Kruse et al., 2025).

The main script for autoBST is in the R Markdown document `BST BulkLoader Workflow Manuscript.Rmd`.

That script uses data files found in subdirectory `data/` and helper functions found in `classyfireAPIs.R`.

The script prepares the BST Bulk Upload files described in the manuscript and writes them to the subdirectory `results/`, along with Sankey diagrams (PDF and PNG) illustrating the data-filtering steps.

The script does not perform the BST runs using the Microsoft Access BST implementation (since these cannot be automated). However, it includes detailed instructions about how to perform them, including where to download the MS Access BST implementation, how to set up the runs, and how to use the Bulk Upload capability. 

A version of the BST Microsoft Access implementation that includes bulk upload capability is included in this repository, in file `BST_BulkUpload.mdb`. By itself, this file will not run the BST; you will need to download the additional helper files provided with the original BST at  [https://www.epa.gov/biosolids/risk-assessment-pollutants-sewage-sludge](https://www.epa.gov/biosolids/risk-assessment-pollutants-sewage-sludge). Detailed instructions can be found in `BST BulkLoader Workflow Manuscript.Rmd`.

The output files of the BST runs described in the manuscript are already included, in subdirectory `results/`.

The script then performs analyses of the BST output files as described in the manuscript. If the subdirectory `analysis/` does not already exist, it will be created, and the results of the analysis (tables and figures) will be written to it.
