## Supplementary figures and images for "A Cheminformatics Workflow for Higher-throughput Modeling of Chemical Exposures from Biosolids"

### ADD_vs_conc.pdf

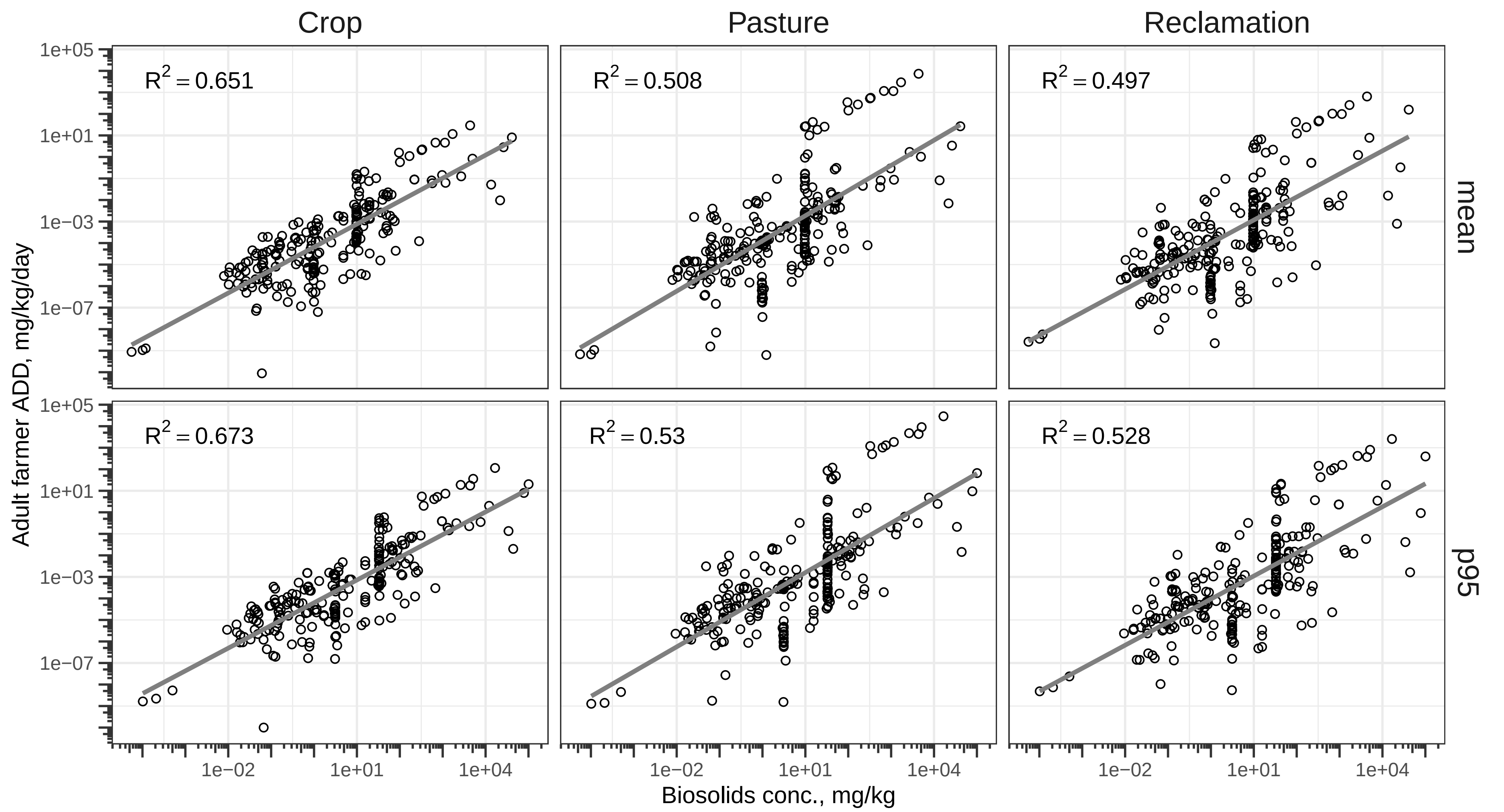

### ADD_vs_conc.png

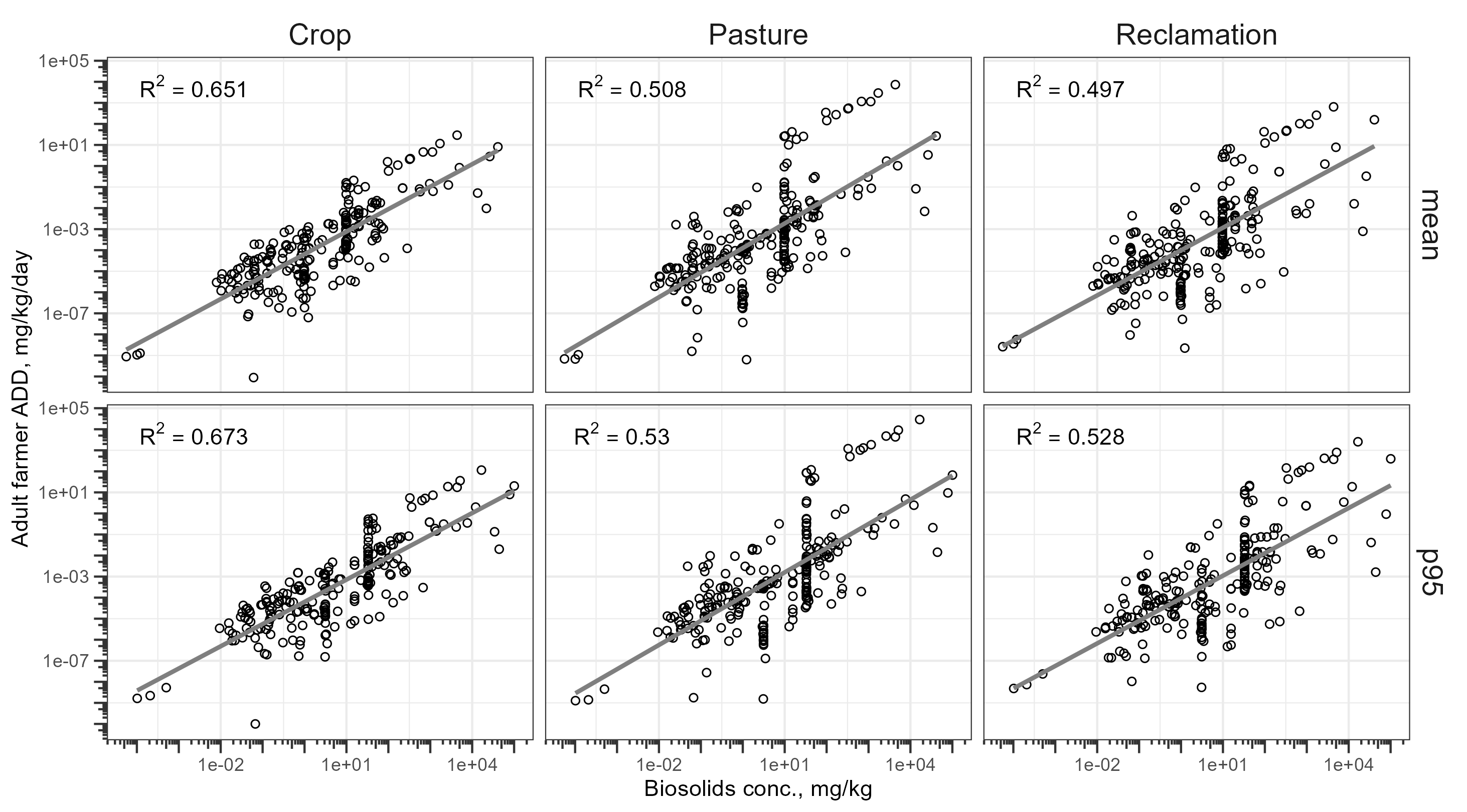

### ADD_vs_conc_logkow.pdf

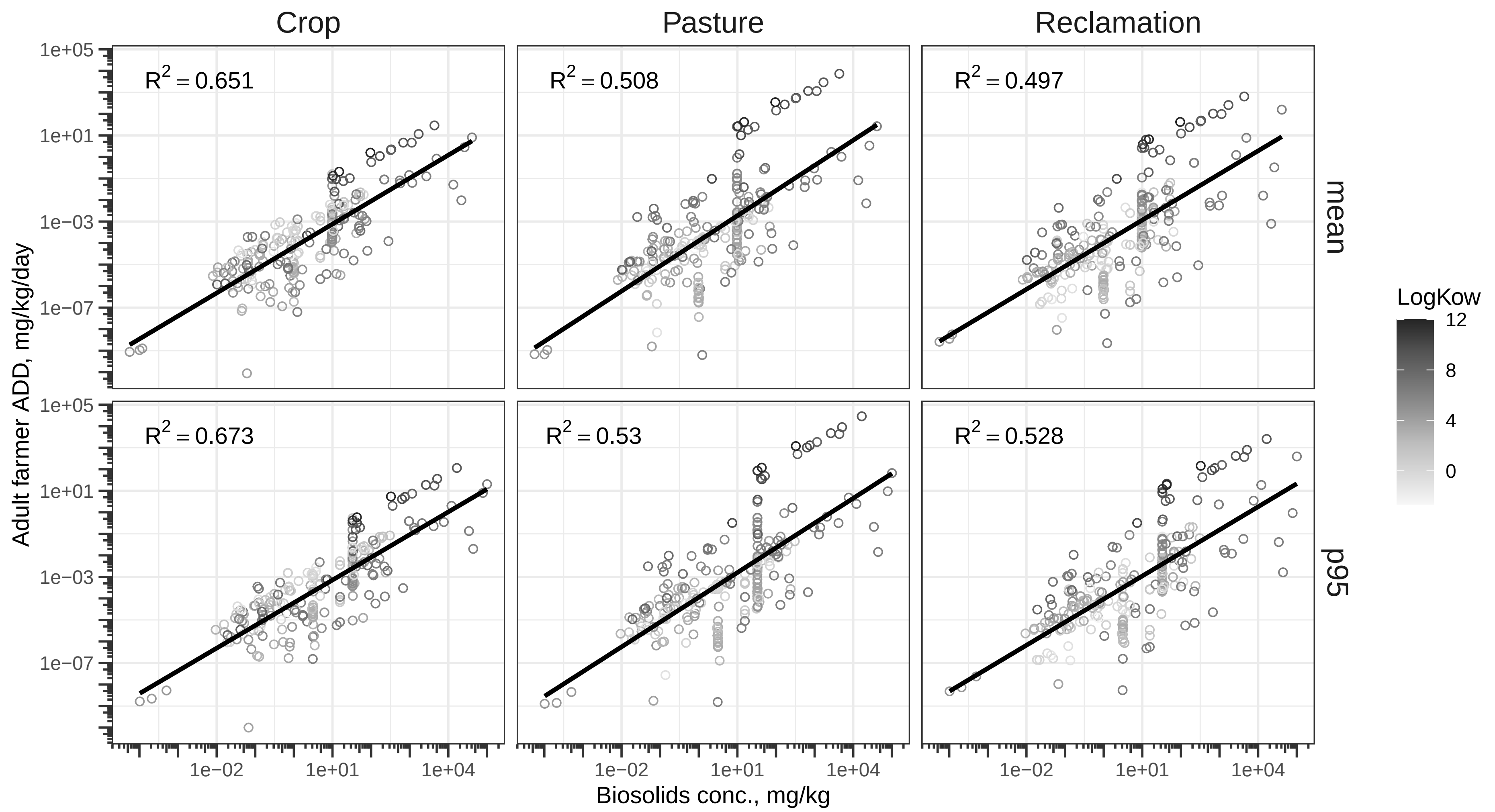

### ADD_vs_conc_logkow.png

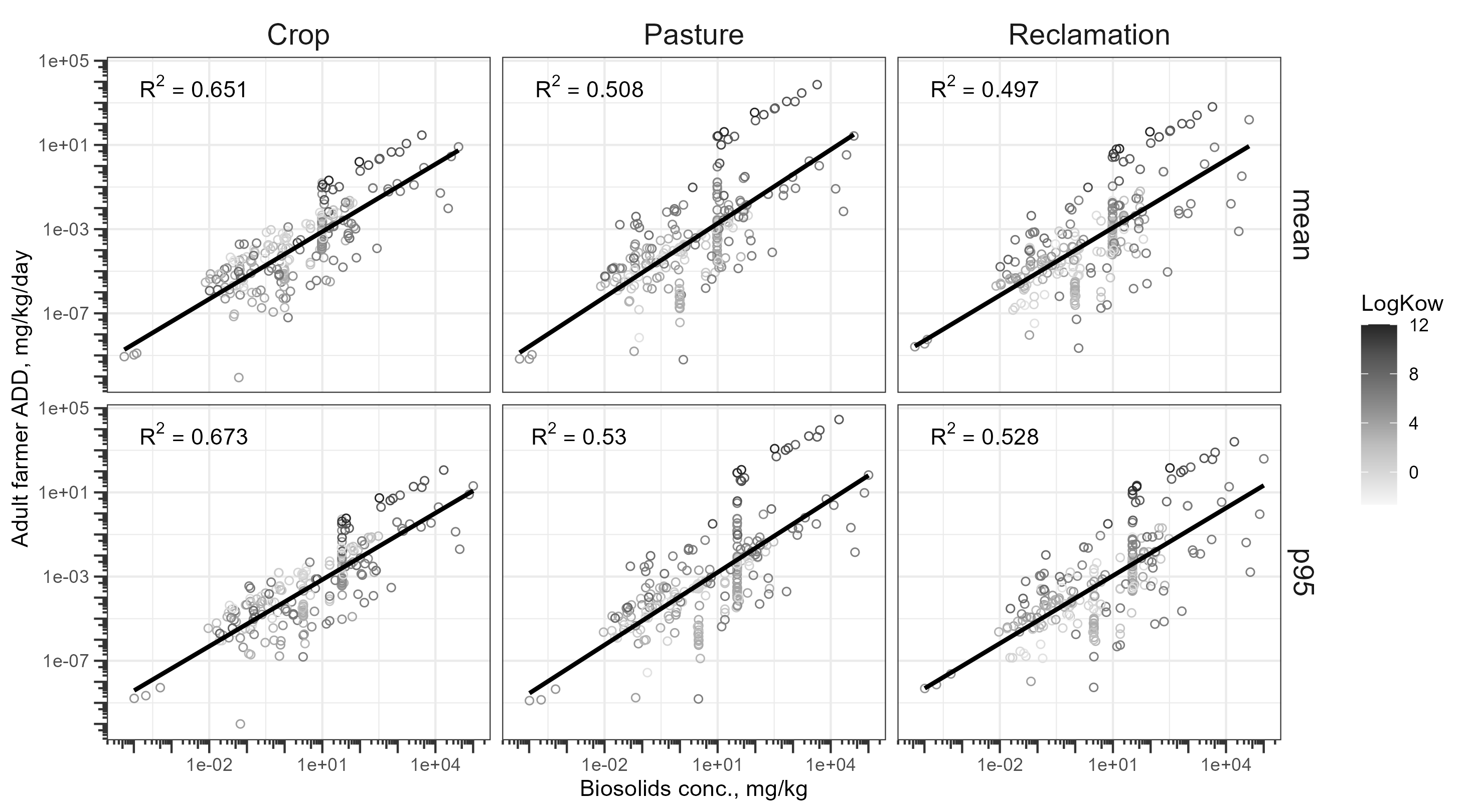

### BST_ADD_freqpoly.pdf

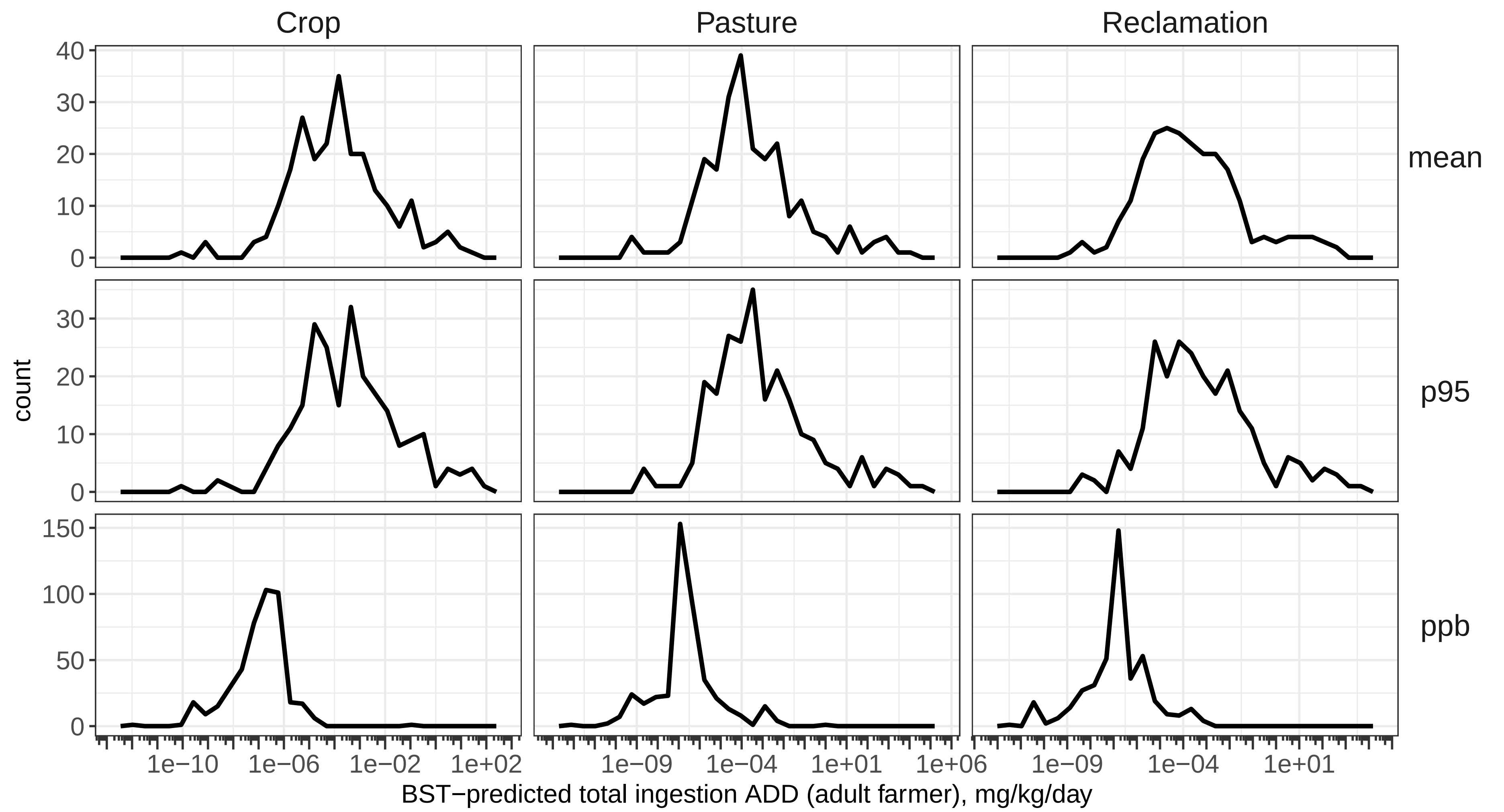

### BST_ADD_freqpoly.png

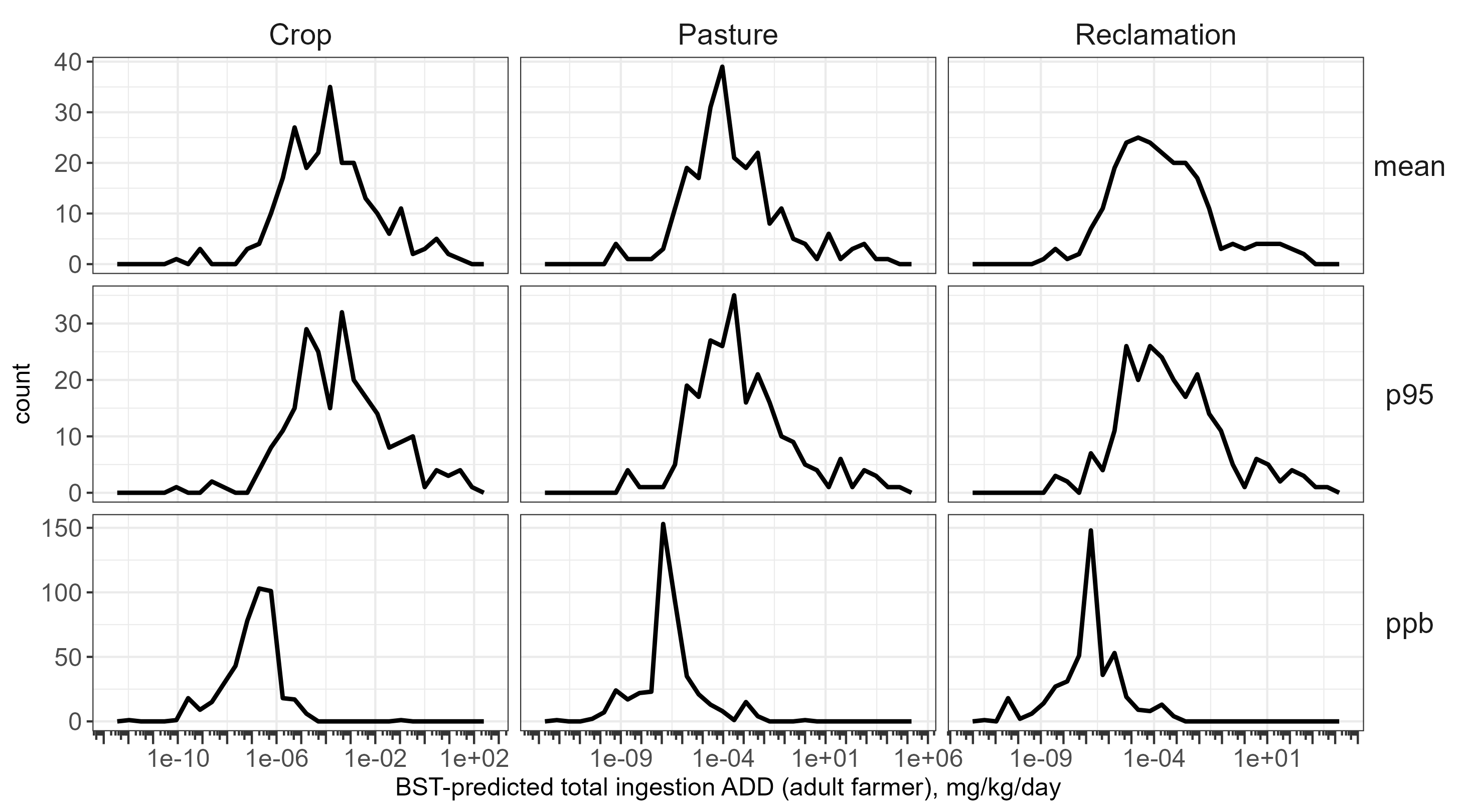

### BST_sankey_1ppb.pdf

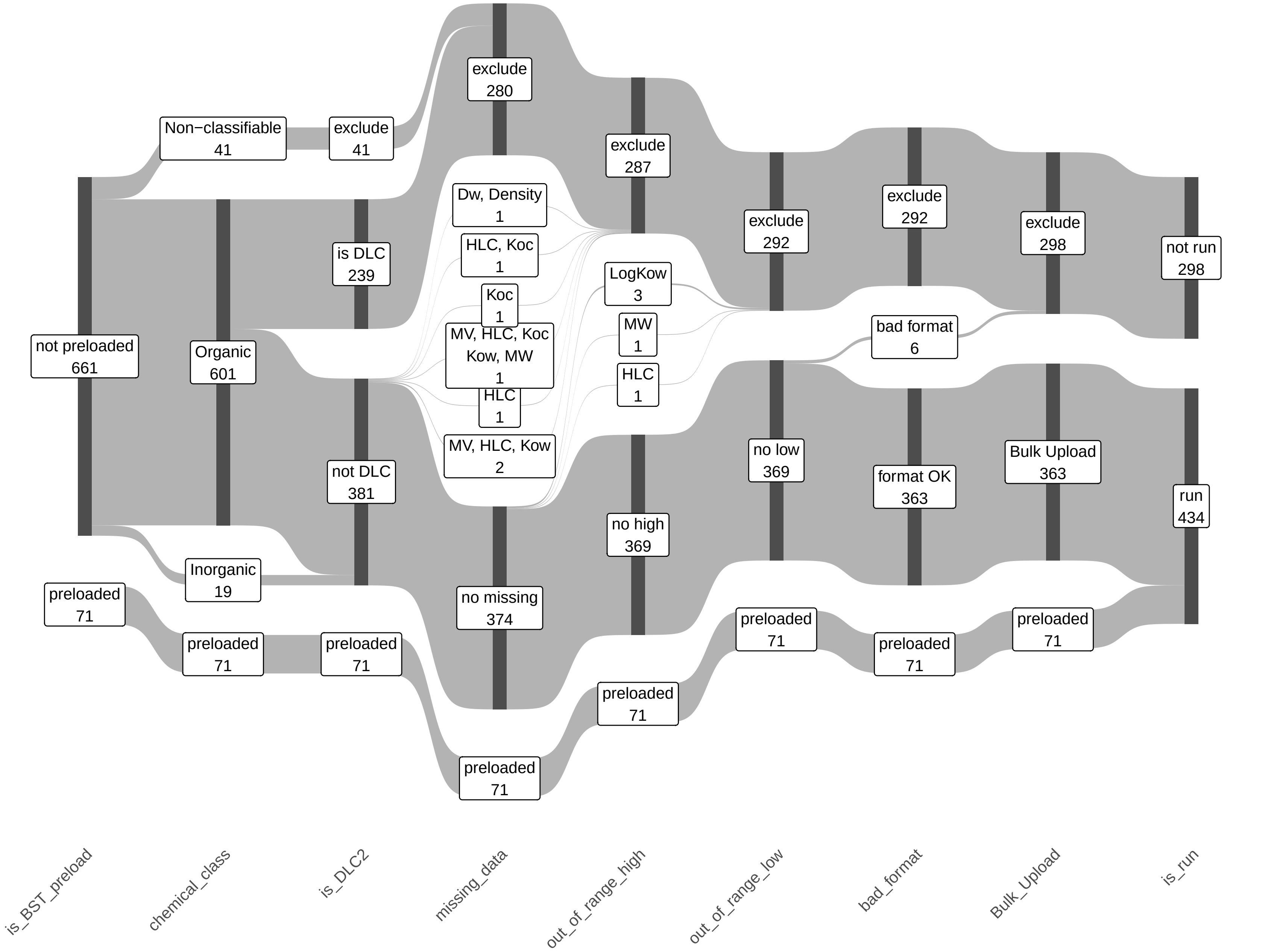

### BST_sankey_1ppb.png

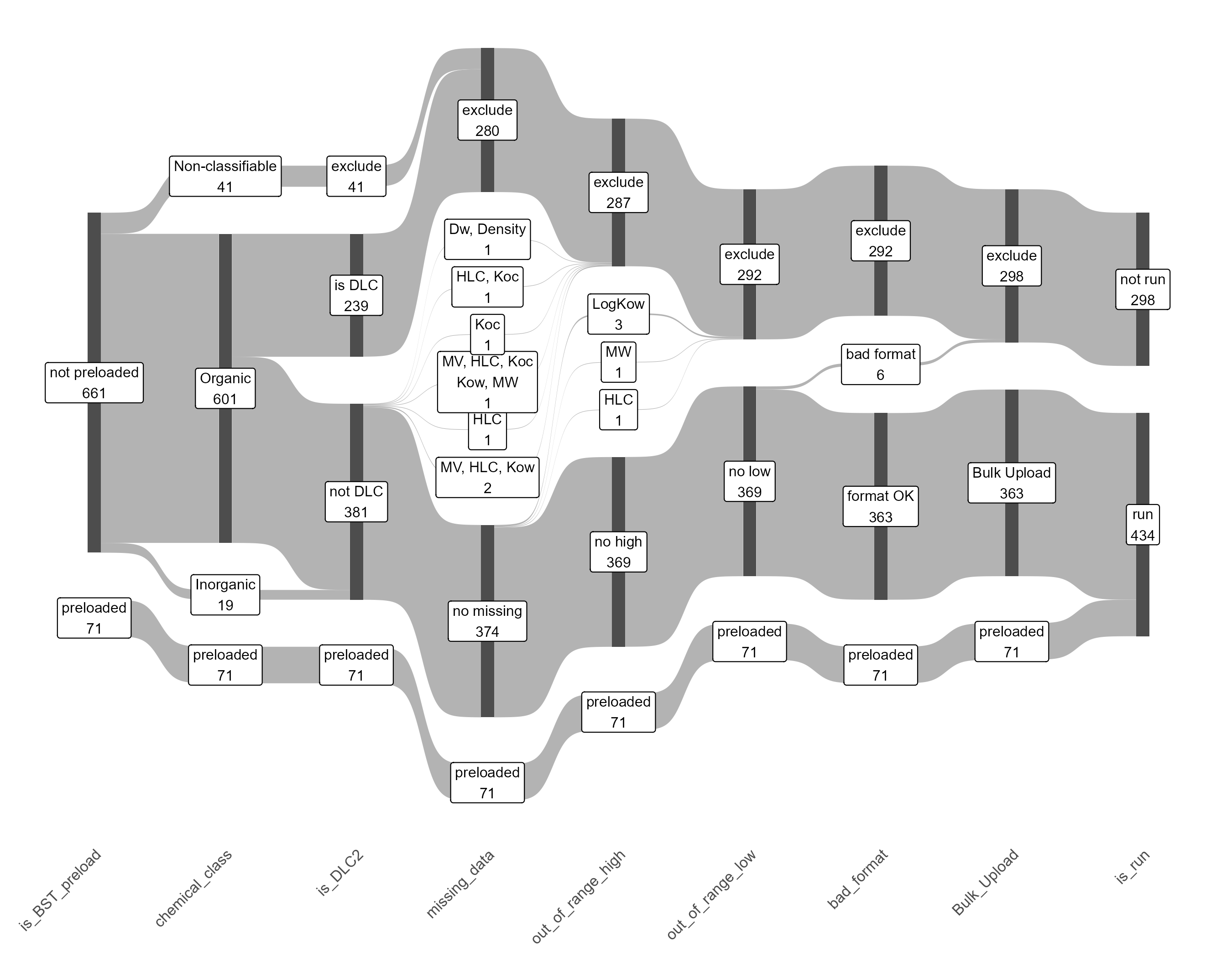

### BST_sankey_NSSS95th.pdf

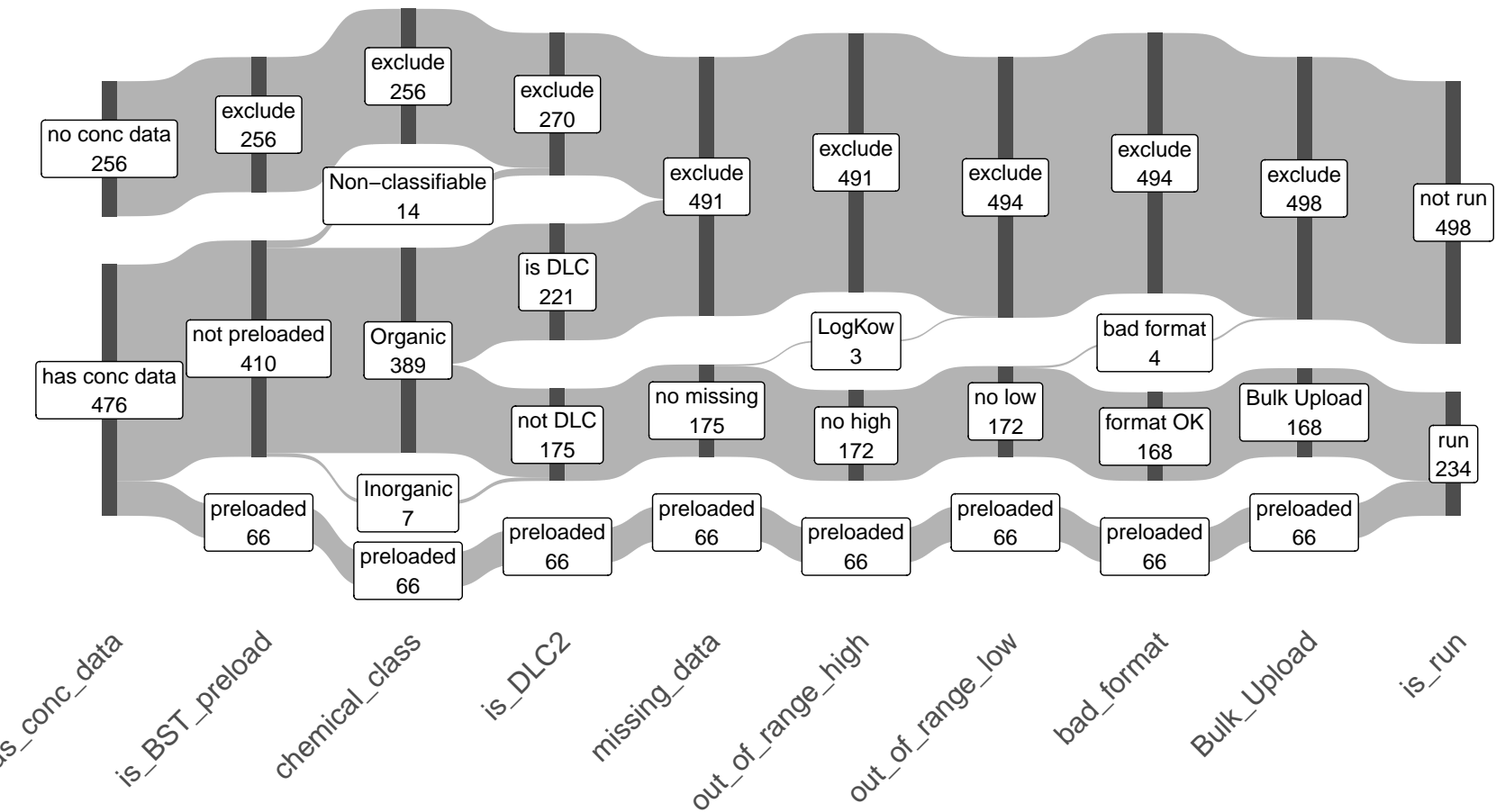

### BST_sankey_NSSS95th.png

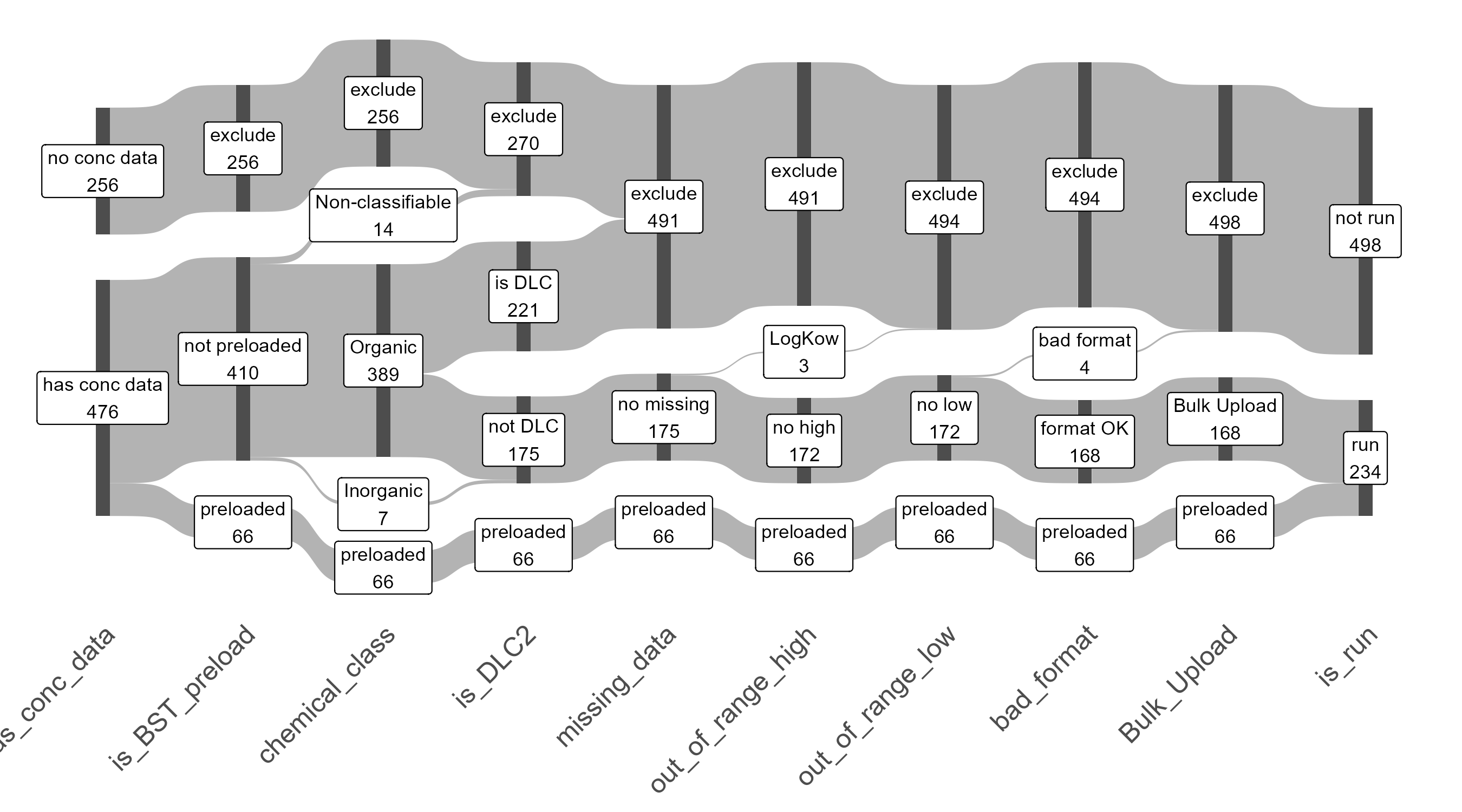

### BST_sankey_NSSSmean.pdf

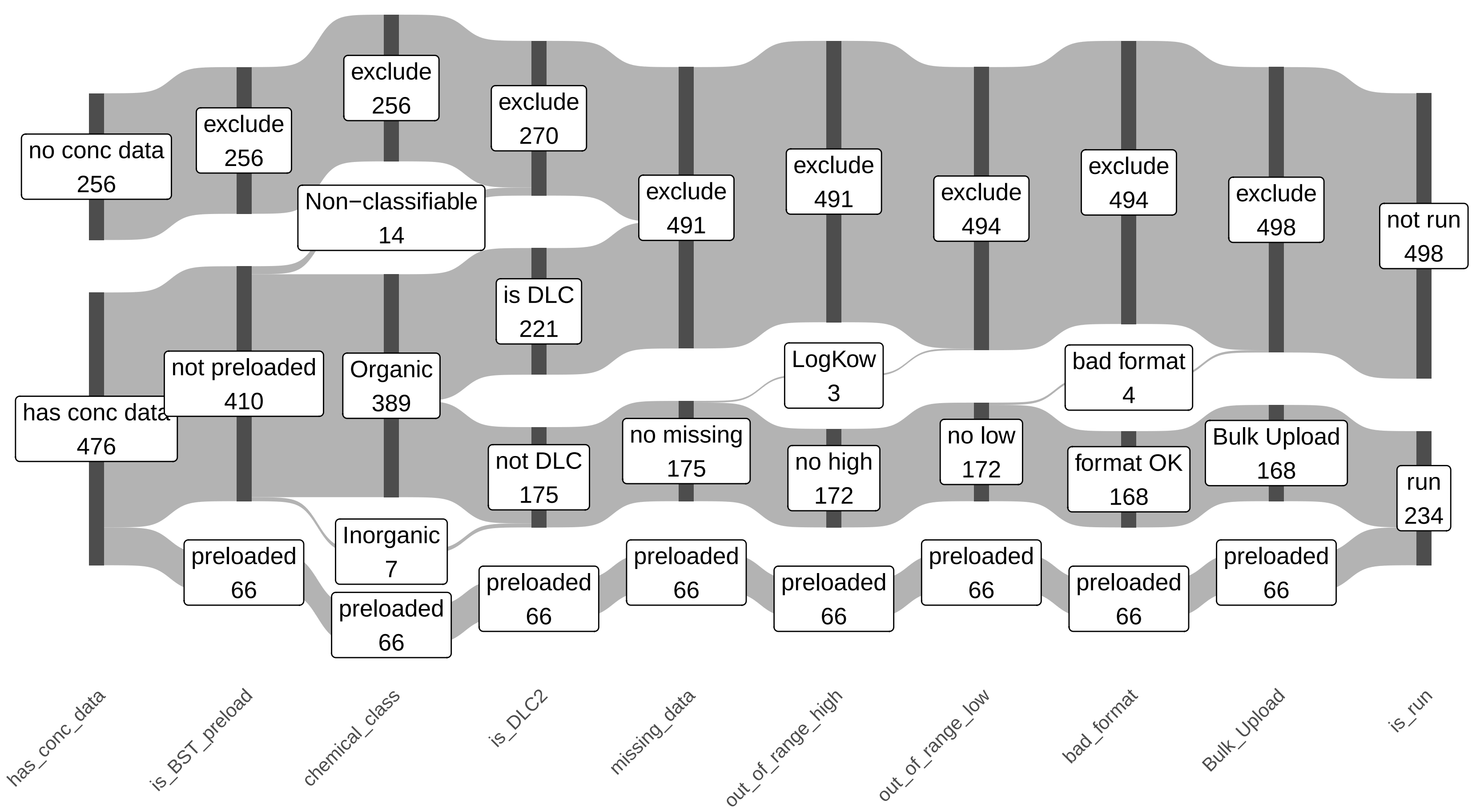

### BST_sankey_NSSSmean.png

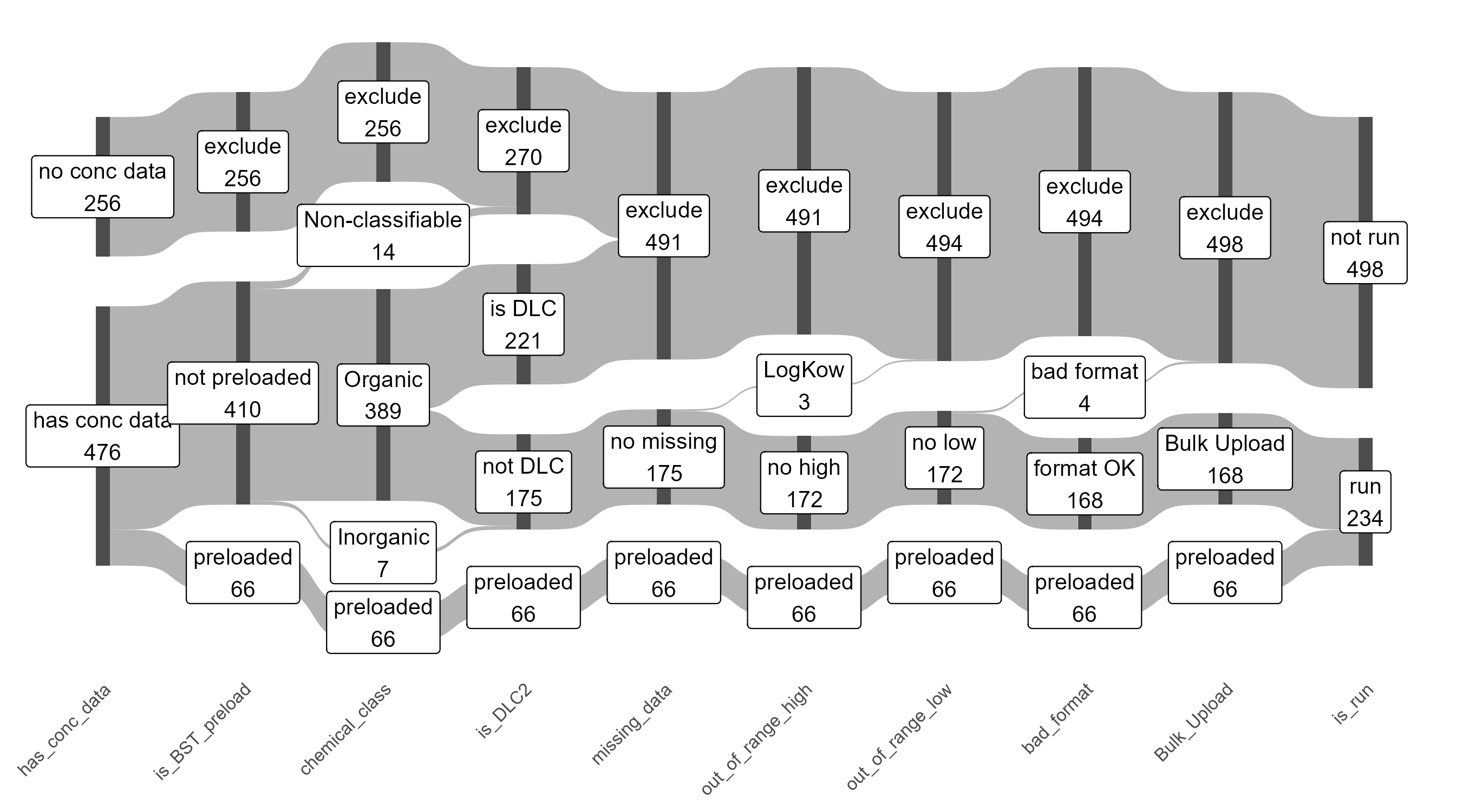
